# Time-resolved volatile organic compound profiling enables non-invasive detection of phenological progression in soybean

**DOI:** 10.64898/2026.08.28.747781

**Authors:** Ryu Nakata, Susumu Hiraga, Masao Ishimoto

## Abstract

**Background and aims:** Plant volatile organic compounds (VOCs) change dynamically with plant development and in response to environmental conditions. However, their potential as non-invasive indicators of phenological progression remains poorly explored. In this study, we developed a framework integrating automated VOC sampling, time-resolved VOC profiling, and machine-learning analysis for the non-invasive assessment of plant phenology. Using soybean (*Glycine max* (L.) Merr.), we investigated whether development-associated temporal variation in VOC emissions could delineate and predict developmental phases.

**Methods:** We collected VOCs daily under controlled environmental conditions from 16 to 43 days after sowing, spanning the transition from vegetative to reproductive stages, using an automated sampling system coupled with thermal desorption-gas chromatograph-mass spectrometer (TD-GC-MS). To characterise temporal changes in VOC profiles associated with phenological progression, we analysed the daily VOC data using a multi-step pipeline combining statistical filtering and similarity-based network analysis. We defined VOC-derived developmental phases from similarity patterns in the VOC profiles, then developed and evaluated machine-learning models to predict these phases.

**Key results:** Seven VOCs exhibited distinct phase-dependent dynamics, including green leaf volatiles and monoterpenes showing characteristic temporal changes during phenological progression. Network-based clustering of VOC profiles resolved five developmental phases closely aligned with conventional developmental stages. A machine-learning model predicted these phases from the VOC profiles with high predictive accuracy on independent test data, demonstrating that phenological progression could be quantitatively inferred from VOC emission patterns.

**Conclusions:** Our findings support VOC profiling as a reliable and non-invasive approach for assessing phenological progression in soybean. By extracting temporally structured VOC signals, this framework captures developmental information that may be difficult to obtain through visual observation alone, particularly after canopy closure. VOC profiling offers a practical tool for monitoring crop developmental dynamics and has broader potential for plant phenotyping and precision crop management.

## Introduction

Plants produce and emit a chemically diverse array of volatile organic compounds (VOCs) throughout their development. More than 1,700 volatile compounds have been identified across 90 plant families (Knudsen *et al*., 2006), with terpenoids, benzenoid/phenylpropanoids, and fatty acid derivatives representing the major classes (Dudareva *et al*., 2013). Although VOC biosynthesis is genetically regulated, VOC emission patterns are highly plastic and respond dynamically to developmental and environmental cues. Both constitutive emissions and rapid VOC bursts induced by biotic and abiotic stressors have been well documented (Kesselmeier and Staudt, 1999; Dudareva *et al*., 2006; Holopainen and Gershenzon, 2010; Loreto and Schnitzler, 2010; Murali-Baskaran *et al*., 2022).

Rather than the presence or absence of individual VOCs, their relative abundances and compositional patterns often convey physiologically relevant information (Niederbacher *et al*., 2015). Accordingly, VOC profiles represent an integrated chemical signature of plant physiological status. Emissions of methanol, ethylene, and related small compounds are closely associated with processes such as cell expansion, maturation, and senescence, as reported in species including common bean, cotton, and soybean (Aharoni *et al*., 1979; Nemecek-Marshall *et al*., 1995; Hüve *et al*., 2007). Phenological progression, including leaf aging, bud maturation, and flowering, is accompanied by coordinated shifts in terpene and benzenoid composition, as observed in wheat, peppermint, sweet orange, and water lily (Batten *et al*., 1995; Gershenzon *et al*., 2000; Killiny and Jones, 2017; Liu *et al*., 2023). In addition to development-associated variation, VOC profiles are strongly shaped by biotic and abiotic stressors. Herbivore feeding and pathogen infection induce characteristic VOC emissions across diverse plant species, such as barrel medic, maize, rice, and tomato (Turlings *et al*., 1990; Takabayashi *et al*., 1991; Obara *et al*., 2002; Sharifi *et al*., 2018; Shiojiri *et al*., 2018; Dreher *et al*., 2019). Abiotic environmental factors such as temperature variation, drought, and nutrient limitation also modulate VOC profiles, as reported in apple and rapeseed (Ebel *et al*., 1995; Gouinguené and Turlings, 2002; Höfer *et al*., 2022).

Given the strong association between VOC profiles and plant physiological state, non-invasive VOC sensing is increasingly being investigated as a means of assessing crop status and informing management decisions related to pest control, fertilisation, and irrigation (Jansen *et al*., 2011; Niederbacher *et al*., 2015). While VOC-based monitoring of biotic and abiotic stresses has advanced considerably, its use for phenological assessment remains limited. Existing examples include field-scale analyses of VOC fluxes in relation to growth dynamics in winter wheat (Loubet *et al*., 2022) and phenology-resolved analyses of VOC composition and exchange in maize (Wiß *et al*., 2017). Temporal variation in VOC profiles during plant development reflects underlying physiological processes, including shifts in specialised metabolism and cell wall remodelling (Laothawornkitkul *et al*., 2009; Dudareva *et al*., 2013). Such biochemical signals may provide complementary phenotypic information that is not fully captured by image-based phenotyping approaches alone, such as hyperspectral imaging (Mishra *et al*., 2017). Therefore, VOC-based sensing has considerable potential as a non-invasive approach for tracking phenological progression in plants.

In this study, we present a framework that integrates analytical chemistry and machine-learning approaches to capture plant developmental dynamics from time-resolved VOC profiles. Specifically, we extracted and interpreted temporally structured VOC signals to delineate VOC-derived developmental phases, which were subsequently used as prediction targets for developing and evaluating machine-learning models. As a proof of concept, we applied this framework to soybean (*Glycine max* (L.) Merr.), one of the most important legume crops, and investigated whether temporal variation in VOC emissions could serve as a non-invasive biochemical fingerprint of phenological progression. In soybean, the transition from vegetative to reproductive growth is central to both photoperiodic flowering research and yield determination (Watanabe et al., 2012; Lin et al., 2021; Vogel et al., 2021). However, direct observation of flowers and developing pods becomes increasingly difficult after canopy closure, as reproductive organs develop at nodes across the plant and are often obscured by dense foliage, thereby limiting conventional phenological assessments. We hypothesised that VOC profiling could provide an alternative approach for phenological assessment by capturing physiological changes associated with plant development that are not readily accessible through visual observation. To test this hypothesis, we collected time-resolved VOC profiles from soybean plants at 16 to 43 days after sowing, spanning developmental stages from V3 (third node) to R5 (beginning seed) (Fehr and Caviness, 1977). We then determined developmentally informative VOC features and derived VOC-based developmental phases from similarity patterns in VOC profiles. Finally, we developed and evaluated a machine-learning model to predict these phases from VOC profiles (Fig. 1). Our results demonstrate that phenological information can be inferred from VOC emission patterns and highlight the potential of VOC profiling as a practical, non-invasive approach for monitoring crop developmental dynamics.

**Figure 1.**
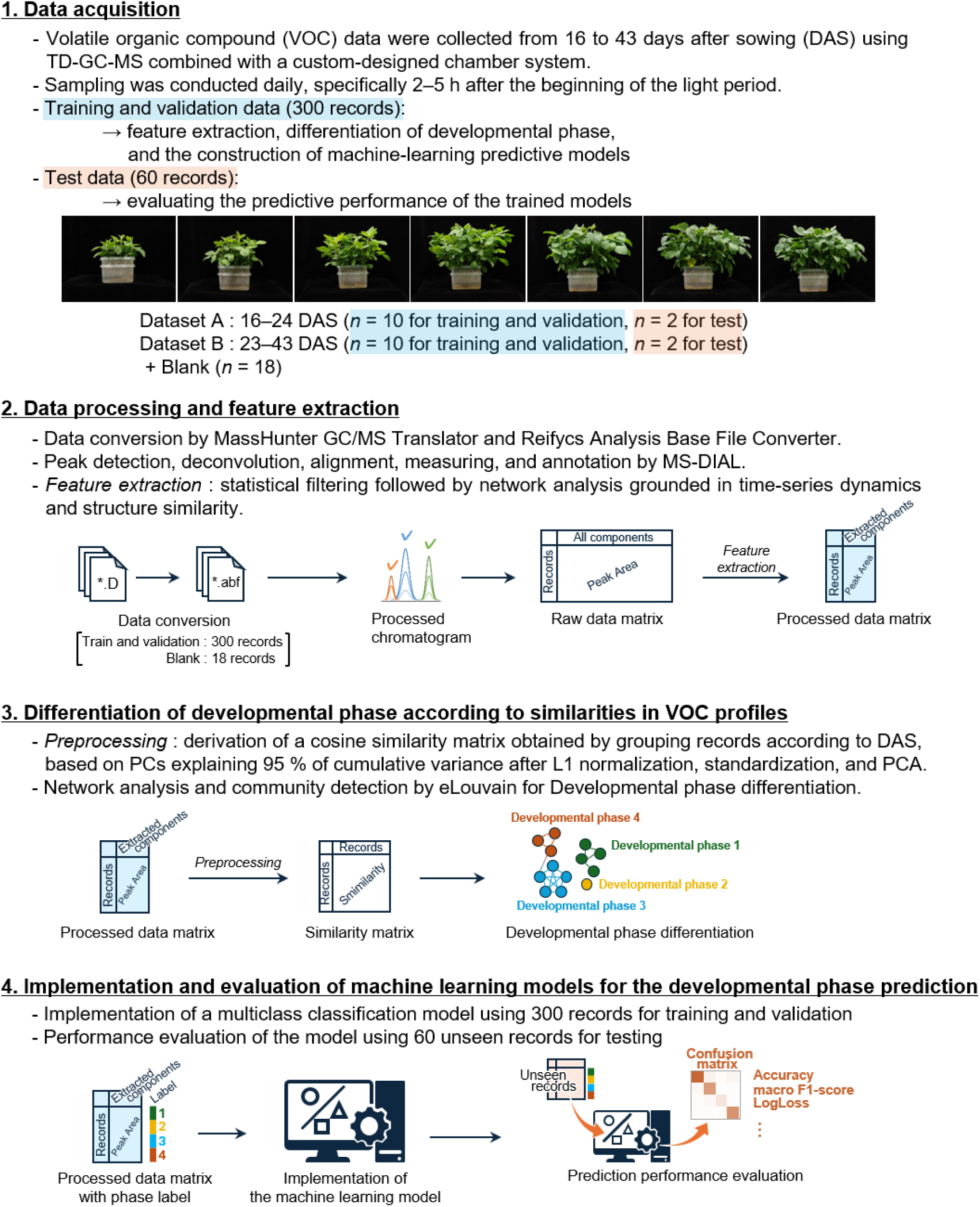
Schematic overview of the experimental workflow for VOC acquisition and data analysis.

## Materials and methods

### Plants and growth conditions

Twelve soybean seeds (cv. Enrei) were sown in a cell tray (Shinwa Co., Ltd.) containing 35-mL cells filled with vermiculite GL (NITTAI Co., Ltd.). Four days after sowing, eight uniformly germinated seedlings were selected and transplanted into double-stacked 4.3-L polypropylene containers (inner dimensions: 167 × 230 × 120 mm; ASVEL Co., Ltd.). Each upper container was filled with approximately 3 L of vermiculite GL, with a bottom opening fitted with polyethene mesh (2.4-mm mesh size; Watanabetai Co., Ltd.). Each lower container was filled with 2 L of nutrient solution and wrapped in aluminium foil to block the light. The nutrient solution was prepared according to a previously reported composition (Sugiyama *et al*., 2016) with slight modifications as follows: 3.0 mM MgSO_4_, 6.3 mM KNO_3_, 0.87 mM KCl, 1.4 mM KH_2_PO_4_, 0.019 mM Fe(III)-EDTA, 2.5 mM Ca(NO_3_)_2_, 4.5 μM KI, 0.028 mM MnCl_2_, 0.019 mM H_3_BO_3_, 2.3 μM ZnSO_4_, 0.50 μM CuSO_4_, and 0.30 nM Na_2_MoO_4_; pH adjusted to 6.8. To prevent depletion of the nutrient solution, the lower container was replaced with a new one containing 2 L of fresh nutrient solution before the solution level dropped below approximately one-third. The germination and growth conditions were maintained at 26 ± 3 °C and 65 ± 15 % relative humidity (RH) under a 12-h light/12-h dark photoperiod. A plant growth LED (VEFA160WRD; ALTRADER Co., Ltd.) was used to provide a photosynthetic photon flux density (PPFD) of approximately 1,750 μmol m^−2^ s^−1^ at the top of the container, measured using a Light Analyzer (LA-105; Nippon Medical & Chemical Instruments Co., Ltd.). Soybean developmental stages were assigned according to the conventional staging system described by Fehr and Caviness (1977).

### VOC sampling system

Air containing VOCs emitted from soybean plants was drawn from a transparent acrylic chamber and introduced into a thermal desorption-gas chromatograph-mass spectrometer (TD-GC-MS) system. An overview of the automated VOC sampling system is shown in Fig. 2, and its components are described below. The transparent acrylic chamber (internal dimensions: 430 mm width × 380 mm depth × 700 mm height; Fig. 2A-a) was equipped with a quantum sensor (SQ- 110-SS; Apogee Instruments Inc.; Fig. 2A-b) and a thermo-hygrometer (HygroVUE10; Campbell Scientific Inc.; Fig. 2A-c). Sensor outputs were recorded and monitored using a data logger (CR1000X; Campbell Scientific Inc.) and support software (LoggerNet ver. 4.5; Campbell Scientific Inc.). The acrylic chamber was equipped with ports for sensor cable connections, an inlet for clean air filtered through activated carbon (Fig. 2A-d), and an outlet connected to the analytical system. To prevent air leakage, all chamber wall penetrations were sealed with high- airtight cable glands. The acrylic chamber was placed inside a plant growth chamber (LPH- 411PFQDT-SPC; Nippon Medical & Chemical Instruments Co., Ltd.; Fig. 2A-e). Before reaching the switching valve, air from the chamber outlet passed through a moisture-removal line comprising two 500-mL polycarbonate gas-wash bottles connected in series (Kokugo Co., Ltd.; Fig. 2A-f). To remove moisture, these bottles were cooled to approximately –15 °C using a refrigerant (SNOWPACK; Mie Chemical Industry Co., Ltd.) placed in a 3 L vacuum-insulated vessel (AS ONE Co., Ltd.) and were further covered with a thermal insulation bag (Fig. 2B). The two bottles were connected via a membrane-type tube dryer (AQUADRIVE SWT-1.3-060; AGC Engineering Co., Ltd.; Fig. 2A-g). The outlet of the second bottle was connected through a second membrane-type dryer to the IN port (green arrow in Fig. 2A) of a normally closed (N.C.) PTFE three-way solenoid valve (FSM-0306Y-TTG; FLON INDUSTRY Co., Ltd.; Fig. 2A-h). The normally open (N.O.) port, indicated by the orange arrow in Fig. 2A, was connected downstream to a flow meter (RK1200-15-SS-1/4-Air-500 mL/min-0.1 MPa-0 MPa-D; KOFLOC Co., Ltd.; Fig. 2A-i) and an air pump (CM-15-12; Enomoto Micro Pump Mfg. Co., Ltd.; Fig. 2A-j). The N.C. port, indicated by the blue arrow in Fig. 2A, was connected to the TD-GC-MS system (Fig. 2A- k–m). The solenoid valve and the air pump were controlled by a relay controller (SDM- CD16ACA; Campbell Scientific Inc.) operated via LoggerNet software. Soybean plants grown in double-stacked containers were placed inside the acrylic chamber within the plant growth chamber (Fig. 2C and D). Except during sampling periods, the sample gas was vented through the N.O. port. During sampling, the solenoid valve was activated to direct the gas into the TD-GC-MS system. Three sampling lines, each configured as shown in Fig. 2A-a–j, were connected to the TD-GC-MS and operated sequentially under valve and pump control. Each sampling cycle consisted of flow-path equilibration, sample collection, and flow-path purging.

**Figure 2.**
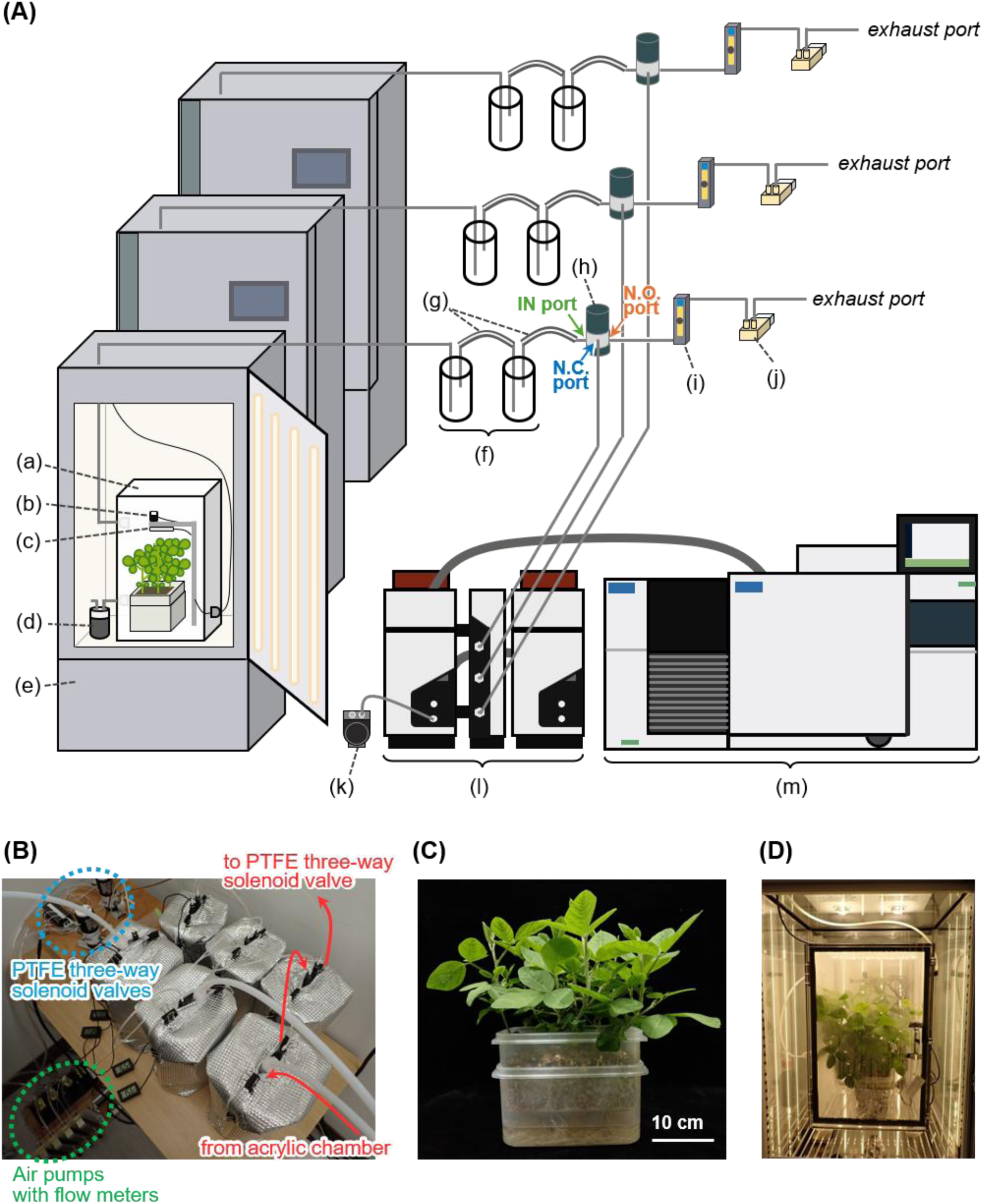
Overview of the automated VOC sampling system. (A) Schematic diagram of the experimental system. (a) acrylic chamber, (b) quantum sensor, (c) thermo-hygrometer, (d) active carbon, (e) plant growth chamber, (f) cold trap, (g) membrane-type tube dryer, (h) PTFE three- way solenoid valve, (i) flow meter, (j) air pump, (k) external pump, (l) thermal desorption (TD) device, and (m) GC-MS apparatus. PTFE or PFA tubing (6 mm i.d.) used between components (d) and (h). PU tubing (6 mm i.d.) downstream of (h) and for the exhaust line, and Sulfinert tubing (2.16 mm i.d.) between (h) and the inlet of the TD device (l). Green, blue, and orange arrows indicate the IN, N.C., and N.O. ports of the solenoid valve, respectively. (B) Cold traps consisting of 500 mL polycarbonate gas washing bottles, refrigerants, 3 L vacuum-insulated vessel enclosed by thermal insulation bags. Pairs of bottles are connected by membrane-type tube dryers. Red arrows indicate one representative flow path from the acrylic chamber through the cold trap to the PTFE three-way solenoid valve. Blue and green dashed lines enclosed the PTFE three-way solenoid valves and air pumps with flow meters, respectively. (C) Soybean plants cultivated in double-stacked containers. (D) Interior view of the plant growth chamber with an acrylic chamber containing soybean plants. Three sampling lines, each consisting of components (a–j), were connected to the TD-GC-MS and operated under valve and pump control in a sequential cycle comprising flow-path equilibration, sample collection, and flow-path purging.

### VOC data acquisition

To enable reproducible and time-resolved acquisition of plant-emitted VOCs, we established an automated multiplexed sampling system integrating controlled-environment chambers with online TD-GC-MS analysis. The system was designed to perform sequential sampling from multiple airflow lines under programmable control of valves and pumps, allowing continuous VOC monitoring with minimal manual intervention.

A double-stacked container with soybean plants was placed in an acrylic chamber (Fig. 2A). The light intensity of the growth chamber was set to 100 %, providing a PPFD of approximately 1,500 μmol m^−2^ s^−1^ as measured at the top of the container. Temperature and relative humidity inside the acrylic chamber were maintained at approximately 26 °C and 60– 80 % RH, respectively. Soybean plants were placed in the acrylic chamber 30 min before sampling. Each sample was collected for 20 min within 2–5 h after the onset of the light period, corresponding to mid-morning. This time window was selected to minimise variability by avoiding transient emission bursts after lights-on and shifts in VOC emission rates later in the photoperiod (Hu *et al*., 2025; Webster *et al*., 2010), thereby enabling the acquisition of more stable VOC emission profiles. Sample gas was introduced directly into an online TD-GC-MS system consisting of a thermal desorption unit equipped with an automated online sampling module and a moisture removal unit (UNITY-xr / UNITY-Air Server-xr / Kori-xr; Markes International Ltd.), connected to an 8890 GC system coupled with a 5977B Inert Plus MSD with Turbo EI and a flame ionisation detector (FID) (Agilent Technologies Inc.). High-purity gases were used, including He (> 99.99995 %), N_2_ (> 99.9995 %), and air supplied from compressed gas cylinders. H_2_ (> 99.9995 %) was generated using a hydrogen generator (H2PEM-100; Parker Hannifin Co.). The sampling and thermal desorption conditions were as follows: standby split flow of 5 mL min^−1^; flow path temperature of 160 °C; sample purge at 100 mL min^−1^ for 2 min; sampling of 2 L at 100 mL min^−1^; line flushing at 50 mL min^−1^ for 1 min. To remove moisture from the sample gas, the Kori unit was operated at –30 °C to trap water and then heated to 300 °C to purge the retained moisture. The sample trap was maintained at –30 °C, followed by trap purging at 50 mL min^−1^ for 2 min. The sample trap was desorbed at 300 °C for 3 min using the maximum heating rate, with a split flow of 3 mL min^−1^. Tenax GR was used as the adsorbent in the sample trap. A diaphragm- type vacuum pump (DAP-6D; ULVAC KIKO Inc.) was used as an external pump for sampling and sample purging. In the GC system, the desorbed sample was split using a pressure control module and introduced into two separate columns: a DB-HeavyWAX (60 m × 0.25 mm i.d., 0.25 µm film thickness; Agilent Technologies Inc.) and a GS-GasPro (60 m × 0.32 mm i.d.; Agilent Technologies Inc.). The DB-HeavyWAX column, operated in constant-flow mode at 1.2 mL min^−1^ and 19.433 psi, was connected to the mass spectrometer. The GS-GasPro column, operated in constant-pressure mode at 2.7 mL min^−1^ and 19.433 psi, was connected to the FID; however, the FID data were not used in this study. The GC inlet conditions were as follows: temperature, 200 °C; pressure, 22.15 psi; flow rate, 12.9 mL min^−1^. The oven temperature was programmed from 40 °C, held for 5 min, and heated to 260 °C at a rate of 10 °C min^−1^. Mass spectrometry was performed in the electron ionisation (EI) mode at 70 eV, with scanning over the *m/z* range of 33–280. The solvent delay was set to 6 min. The ion source and quadrupole temperatures were maintained at 250 °C and 150 °C, respectively.

VOC emissions were measured daily from 16 to 43 days after sowing (DAS). Two datasets were acquired: dataset A covered 16–24 DAS, and dataset B covered 23–43 DAS, each with 10 biological replicates per day. In each dataset, the same biological replicates were followed across sampling days, thereby preserving the longitudinal structure of the measurements. Altogether, these datasets comprised 300 records used for feature extraction, VOC-derived developmental phase delineation, and machine-learning model development. An independent test dataset comprising 60 previously unseen records was collected under the same experimental conditions. This test dataset included two biological replicates per day over the same two sampling periods (16–24 and 23–43 DAS) and was used to evaluate the predictive performance of the model. Containers without soybean plants were analysed separately as blank controls (*n* = 18).

### GC-MS data processing

The acquired raw GC-MS data were converted to CDF format and subsequently to ABF format using MassHunter GC/MS Translator 10.0 (Agilent Technologies Inc.) and the Reifycs Analysis Base File Converter (Reifycs Inc.), respectively. The ABF files were processed using MS-DIAL (version 5.5.250221; Tsugawa *et al*., 2015). Data processing was performed over the *m/z* range of 30–400 and a retention time (RT) range of 6–32 min. Peak detection was conducted with a minimum peak height threshold of 1,000 amplitude. Smoothing was performed using a linear weighted moving average method with a smoothing level of 3 scans and an average peak width of 20 scans. MS1 deconvolution was performed with a sigma window value of 0.2 and an EI spectra cut-off of 5 amplitude. Peak annotation was conducted using the MS-DIAL metabolomics MSP spectral kit containing EI–MS reference spectra, downloaded on September 4, 2024, with a mass tolerance of 0.5 Da and an identification score threshold of 700. Peak alignment was performed using an RT tolerance of 0.5 min and an EI similarity tolerance of 70 %, with a retention time factor of 0.5 and an EI similarity factor of 0.6. Peak annotation was refined after alignment, and gap filling was applied. The peak area matrix was exported into CSV format. All EI–MS spectral data were exported into an MSP format after alignment correction using the *peak curation* function in MS-DIAL. Details of the software tools and spectral resources are provided in Supplementary Table S1.

### Detection of temporally variable VOCs by statistical filtering and network analysis

Temporally variable VOC features were extracted through a multi-step approach consisting of statistical filtering followed by network analysis. The peak area matrix obtained from the MS-DIAL analysis (Supplementary Dataset S1) was filtered to remove unreliable features. Features were retained only when both of the following criteria were met: (i) peak areas exceeded five times the mean blank value in at least 50 samples, and (ii) fewer than 50 % of the samples had peak areas less than or equal to 0.8 times the mean blank value. To identify and remove features potentially influenced by differences between datasets A and B, a linear mixed model (LMM) was fitted to each retained feature using only records from 23 and 24 DAS, which were common to datasets A and B. Datasets (A or B) and DAS were treated as fixed effects, and a replicate was included as a random intercept. P-values for the dataset effect were adjusted using the Benjamini– Yekutieli procedure, and features with a significant dataset effect (false discovery rate < 0.01) were excluded from subsequent analyses.

Network analysis was then applied to select features connected to those retained after the preceding filtering steps in spectral or temporal similarity networks. EI-MS spectra exported from MS-DIAL (Supplementary Dataset S2) were parsed, and fragment-ion intensities within the *m/z* range of 0–500 were binned using a bin width of 0.3 Da to generate normalised intensity vectors for all alignment features. Pairwise spectral similarities were calculated as cosine similarities between these vectors, representing spectral relatedness. Temporal similarities among the features were evaluated by applying soft-dynamic time warping (Soft-DTW; γ = 0.01) to standardised time-series profiles. The resulting distances were converted into similarities with a radial basis function kernel. Spectral and temporal similarity matrices were normalised and thresholded to construct two undirected networks: an EI-MS cosine-similarity network and a Soft-DTW network. In the Soft-DTW network, an edge was retained when the Soft-DTW similarity ranked within the top one-eighth (12.5 %) of all pairwise values. In the EI-MS cosine-similarity network, an edge was retained when the cosine similarity was ≥ 0.60. Features were selected for subsequent analysis if they were connected to at least one other feature in either network and their peak areas were at least five times the mean blank value. All data processing and network integration steps were implemented using Python scripts. The scripts and corresponding outputs are provided in Supplementary Data S1–4.

The selected interconnected features were manually curated using MRMPROBS (ver. 3.66; Tsugawa *et al*., 2013), including peak verification and reintegration. The curated peak area data were used for downstream analyses. The following parameters were applied in MRMPROBS: smoothing method, linear weighted moving average; smoothing level, 3 scans; minimum peak width, 5 scans; minimum peak height, 50 amplitudes; RT tolerance, 1 min; amplitude tolerance, 10 %; minimum posterior, 70 %; compound library file, Supplementary Dataset S3. Putative structural annotation of each peak was performed using the Agilent MassHunter Unknown Analysis software (ver. 10.1) with the NIST17 mass spectral library. For this analysis, *m/z* 28 was excluded, and RT window size factors of 25, 50, 100, and 200 were applied for peak deconvolution. All other settings were kept at their default values. MRMPROBS version and source information are provided in Supplementary Table S1.

### Delineation of developmental phases based on VOC-profile similarities

The peak area matrix of VOCs curated using the MRMPROBS software (Supplementary Dataset S4) was subjected to L1 normalisation followed by z-score standardisation. Principal component analysis (PCA) was performed, and daily centroids were calculated in the PCA space defined by principal components cumulatively explaining 95 % of the total variance. Pairwise cosine similarities between daily centroids were calculated to quantify VOC-profile similarities between DAS. The resulting similarity matrix was used to construct a weighted network, in which nodes and edges represented DAS and pairwise VOC profile similarities, respectively. Community detection was performed using the ensemble Louvain (eLouvain) method (Evkoski *et al*., 2021) to delineate VOC-derived developmental phases. To maximise the weighted modularity score, the edge threshold (0.5–0.9), resolution (0.01–10), and matching threshold (0.6–1.0) were optimised using Optuna with the Tree-structured Parzen Estimator sampler over 200 trials (Akiba *et al*., 2019). All analyses were implemented using Python scripts, and the scripts and corresponding outputs are provided in Supplementary Data S5.

### Implementation and evaluation of machine-learning models for developmental phase prediction

To assess whether soybean developmental phases could be predicted from VOC profiles, we implemented a Tabular prior-data fitted network (TabPFN)-based classification pipeline. TabPFN is a transformer-based model pretrained on synthetic tabular data and designed to perform inference without task-specific fine-tuning (Hollmann *et al*., 2025). We used AutoTabPFNClassifier, a post-hoc ensemble extension of TabPFN that selects optimal prediction heads based on cross-validated performance. The dataset comprised 300 records with selected VOC features and VOC-derived developmental phase labels assigned by community detection analysis described above (Supplementary Dataset S5) and was split into stratified training and validation subsets at a 75:25 ratio. To address class imbalance, KMeansSMOTE was applied only to the training data (Douzas *et al*., 2018). To maximise the silhouette coefficient, related hyperparameters—including the number of clusters (*n_clusters*), number of nearest neighbours (*k_neighbors*), and cluster balance threshold (*cluster_balance_threshold*)—were optimised using Optuna within the ranges of 1–15, 1–15, and 0.01–0.30, respectively. All features were standardised by z-score transformation, with the scaler fitted on the resampled training data. The AutoTabPFNClassifier was fitted to the resampled training set using the *best_quality* preset, with probability balancing enabled and model selection based on the *roc_auc_ovo_macro* metric. The ensemble was run with a time budget of 3,000 s. Model performance was evaluated using both the 25 % hold-out validation set and an independent test set comprising 60 previously unseen records acquired under the same experimental conditions. For the independent test set, peak areas of the seven selected VOCs were quantified using MRMPROBS with parameters and procedures as described above. Reference labels were assigned according to the VOC-derived developmental phases defined by the preceding community detection analysis. Accuracy, balanced accuracy, macro precision, macro recall, macro F_1_ score, macro averaged receiver operating characteristic area under the curve (ROC-AUC), and log loss were calculated. The 95 % confidence intervals were estimated by stratified bootstrapping (*n* = 1,000). All analyses were implemented using Python scripts, and the scripts and corresponding outputs are provided in Supplementary Data S6.

### Statistical analysis and software environment

Statistical analyses, network analyses, and machine-learning modelling were performed in a Python 3.13.5 environment managed with Conda (version 25.7.0), with scripts developed executed using JupyterLab (version 4.4.9). The main Python packages were as follows: numpy 2.2.6, scipy 1.16.2, pandas 2.3.3, statsmodels 0.14.5, scikit-learn 1.6.1, imbalanced-learn 0.14.0, tslearn 0.6.4, tabpfn 2.2.1, tabpfn-extensions 0.1.6, networkx 3.5, elouvain 0.1, optuna 4.5.0, torch 2.10.0+cu128, matplotlib 3.10.7, and seaborn 0.13.2.

## Results

### VOC features associated with phenological progression

Using a multi-step analytical framework combining statistical filtering and similarity-based network analysis, we extracted VOC features associated with phenological progression from time-resolved VOC datasets. MS-DIAL analysis of all samples (300 records) detected 250 peaks (Supplementary Dataset S1). After blank-based filtering, 11 peaks were retained based on peak area values. Statistical filtering using LMM to remove features exhibiting significant dataset effects further reduced this set to three peaks: Alignment ID (AID)_28, 128, and 153 (Supplementary Data S1). These three peaks were considered statistically supported VOC features associated with phenological progression. To uncover additional VOC features that might be related to phenological progression but were not selected by the statistical filtering, we constructed a multi-layer similarity network using Soft-DTW and cosine similarity. This analysis revealed four additional peaks, AID_122, 142, 143, and 154, which were directly connected to at least one of the statistically selected peaks, suggesting that they shared similar temporal dynamics and/or spectral characteristics. Thus, the network analysis expanded the set of candidate VOC features from three statistically supported peaks to seven candidate VOC features potentially associated with phenological progression (Supplementary Fig. S1).

The seven peaks were tentatively annotated using the NIST17 mass spectral library. In the order of retention time, they corresponded to 3-methylfuran, (*E*)-2-hexenal, *β* -ocimene, (*E*)- 3-hexenyl acetate, (*Z*)-3-hexenyl acetate, *trans*-alloocimene, and (*Z*)-3-hexen-1-ol (Supplementary Fig. S2 and Table 1). AID_122, 142, 143, and 154 displayed increased emission from approximately 23 DAS, followed by a decline after approximately 32 DAS (Fig. 3). Based on tentative annotation, we classified these compounds as C6 aliphatic alcohols and their oxygenated derivatives, likely originating from the lipoxygenase pathway and commonly referred to as green leaf volatiles. In contrast, AID_128 and 153 exhibited delayed induction beginning around 26 DAS, followed by a decrease after approximately 35 DAS (Fig. 3). They were provisionally assigned as monoterpenes derived from the terpenoid biosynthetic pathway. AID_28 was tentatively classified as a furan compound and showed a gradual increase in abundance from the vegetative stage through flowering and seed filling, consistent with previous observations in wheat (Batten *et al*., 1995), although its biosynthetic origin remains unclear. Overall, the proposed framework extracted a compact set of VOC features associated with phenological progression, capturing coordinated metabolic shifts across multiple biosynthetic pathways.

**Figure 3.**
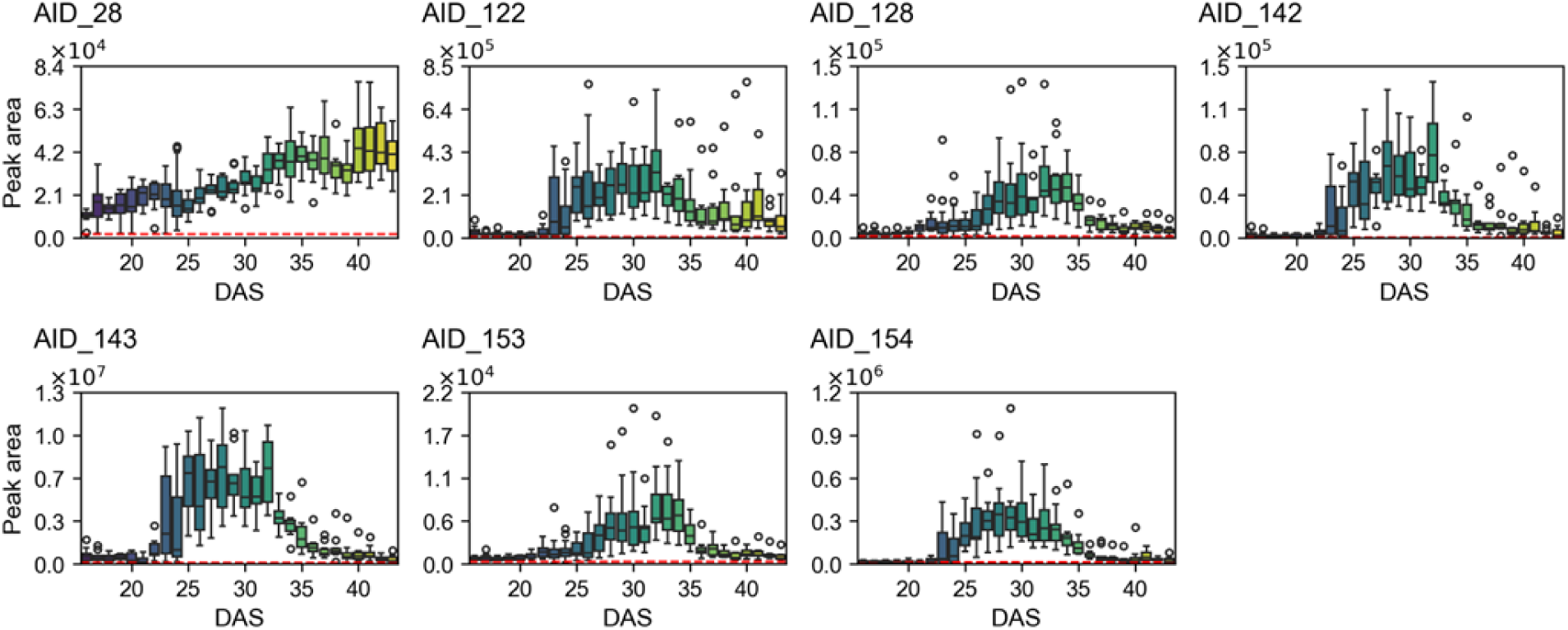
Temporal variations in VOC emission profiles during soybean development (*n* =10–20). Red dashed lines represent mean level in blank records. In the box plots, outliers outside 1.5 × the interquartile range (IQR) are shown as open dots, and the whiskers indicate the minimum and maximum values within this range.

**Table 1.** VOCs identified through feature extraction combining blank-based filtering followed by network analysis using temporal dynamics and structure similarity.

| Alignment ID | Retention time (min) | Tentative compound name | Match Factor |
| --- | --- | --- | --- |
| AID_28 | 7.16 | 3-Methylfuran | 96.3 |
| AID_122 | 13.96 | (*E*)-2-Hexenal | 98.2 |
| AID_128 | 14.49 | β-Ocimene | 97.8 |
| AID_142 | 15.46 | (*E*)-3-Hexenyl acetate | 94.6 |
| AID_143 | 15.62 | (*Z*)-3-Hexenyl acetate | 98.3 |
| AID_153 | 16.43 | *trans*-Alloocimene | 88.4 |
| AID_154 | 16.50 | (*Z*)-3-Hexen-1-ol | 97.5 |

### Developmental phase delineation from VOC profile similarity

To investigate whether developmental structure was embedded in the extracted VOC features, we applied a network-based clustering framework to delineate developmental phases based on VOC profile similarities. After manual peak curation using MRMPROBS (Supplementary Dataset S4), the peak area matrix comprising 300 records was L1-normalised per sample and standardised per variable. The processed data were projected into the PCA space, and the first four principal components explaining over 95 % of the cumulative variance (Supplementary Fig. S3A) were used to calculate centroids for each DAS. Pairwise cosine similarities between daily centroids were computed, and the resulting similarity matrix was visualised as a heatmap in Fig. 4A. A network was then constructed using cosine similarity as edge weights, followed by community detection using the eLouvain method. Key parameters, including the edge selection threshold, resolution parameter, and co-occurrence matching threshold, were optimised using Optuna to maximise weighted modularity (Supplementary Fig. S3B). This analysis enabled the identification of five distinct communities based on VOC profile similarities (Supplementary Table S2), which were interpreted in relation to the observed phenological progression: vegetative growth (16–19 and 22–24 DAS: phase 1), bud maturation (20–21 DAS: phase 2), flowering and pod emergence (25–32 DAS: phase 3), pod elongation (33–35 DAS: phase 4), and seed filling (36–43 DAS: phase 5) (Fig. 4B). These VOC-derived phases were broadly consistent with the developmental stages assigned using the conventional soybean staging system (Supplementary Table S3). According to conventional staging, the R1 stage (beginning bloom) occurs at 25 DAS and earlier time points are generally assigned as the vegetative stage. However, the VOC-derived phase delineation resolved the 20–21 DAS period as a distinct phase corresponding to bud maturation (phase 2), not represented as a separate stage in the conventional staging system. During this phase, B1-stage floral buds become visible (Supplementary Fig. S4), characterised by the banner petal extending beyond the calyx tip (Peterson *et al*., 1992). Following the R1 stage, plants progressed through the R2, R3, R4, and R5 (full bloom, beginning pod, full pod, and beginning seed) stages at 27, 30, 33, and 36 DAS, respectively. These conventional stages corresponded well with the VOC-derived phase delineation, with R1–R3, R4, and R5 corresponding to phases 3, 4, and 5, respectively (Supplementary Table S3).

**Figure 4.**
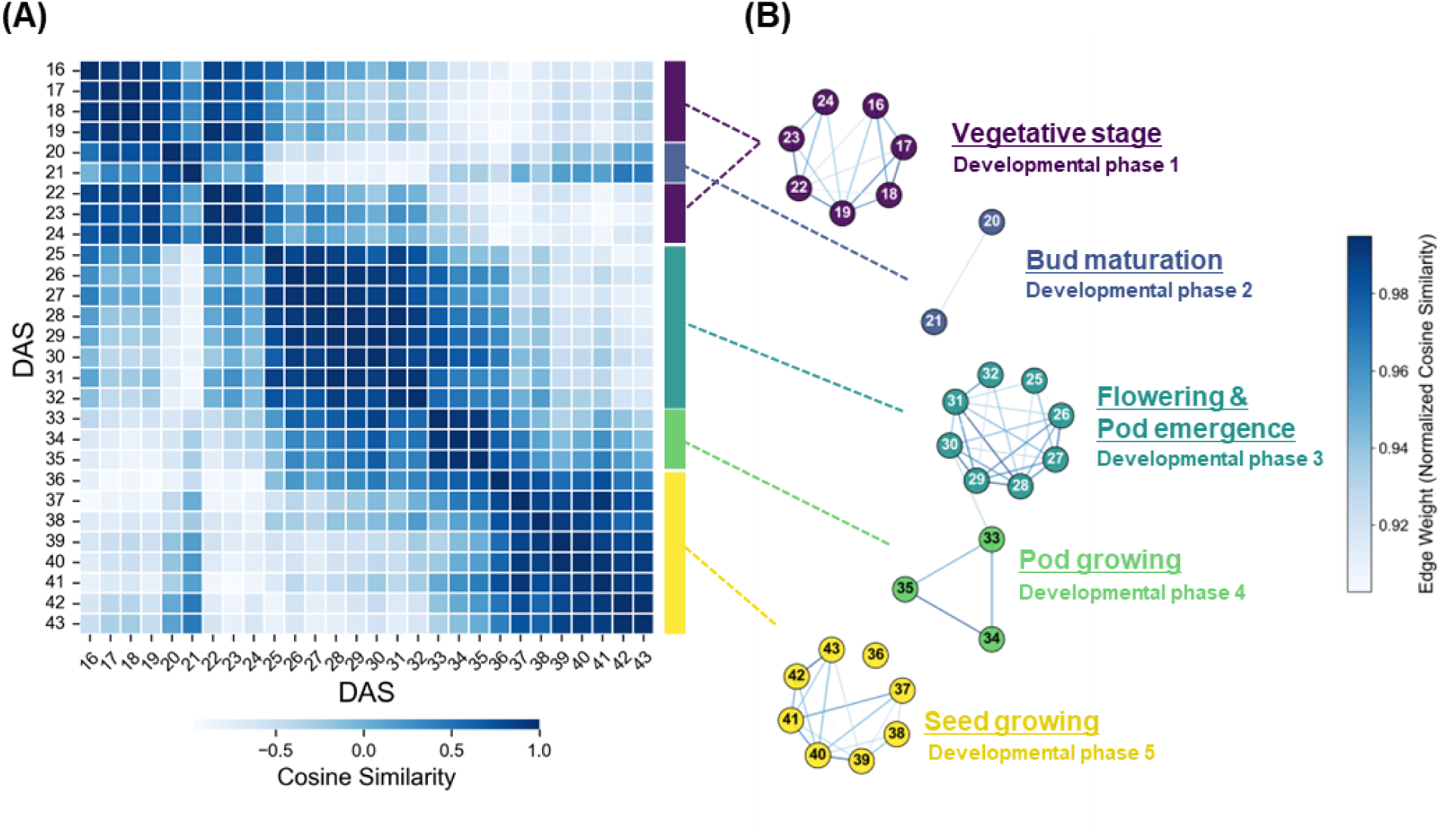
Differentiation of developmental phase based on VOC profiles. (A) Heatmap of the similarity matrix calculated from VOC profiles across days after sowing (DAS). (B) Cosine similarity-weighted network with communities detected using the eLouvain method, annotated with developmental phases assigned to each community based on a conventional staging method. Node labels represent DAS.

These results indicate that VOC profiles capture coordinated physiological and metabolic transitions during soybean development, consistent with previous studies reporting stage-dependent shifts in VOC emission, including phenology-resolved profiles in wheat (Batten *et al*., 1995), developmentally regulated floral volatiles in snapdragon (Dudareva *et al*., 2000), and stage-specific VOC signatures in developing soybean seeds (Boué *et al*., 2003). However, these studies were based mainly on broadly defined developmental stages or on VOCs derived from specific tissues. In contrast, our results demonstrated that whole-plant VOC profiles measured daily from the vegetative (V3) to seed filling (R5) stage closely track soybean phenological progression, thereby supporting headspace VOC profiling as a sensitive and non-invasive phenological marker.

### VOC profile-based developmental phase prediction using machine learning

To assess whether VOC profiles could be used to predict the developmental phases, we developed machine-learning models and evaluated their performance using an independent test dataset, comprising 60 previously unseen records collected independently and not used for VOC feature selection, phase delineation, or model development (Fig. 1). Accurate prediction of developmental phases in this dataset would support the validity of VOC-derived phase delineation and the feasibility of non-invasive phenological assessment using VOC profiles. The model-development dataset comprising 300 records was split into training (75 %) and validation (25 %) sets using stratified sampling. Class imbalance in the training data was addressed using KMeansSMOTE, and the resulting features were standardised by z-score transformation. We then fitted a TabPFN model on the processed training data and evaluated the predictive performance using the hold-out validation set and the independent test dataset. On the validation set, the model achieved an accuracy of 0.893, a macro F_1_-score of 0.858, and a macro ROC-AUC of 0.982, indicating robust generalisation across the developmental phases (Supplementary Fig. S5). Performance remained high on the independent test dataset, with an accuracy of 0.917 and consistently high values for the other evaluation metrics (Fig. 5A). Furthermore, the confusion matrix and class-wise ROC curves showed clear separation among the developmental phases (Fig. 5B and C). However, predictive performance for developmental phase 2 was comparatively lower, likely reflecting its limited representation in the training data (Fig. 5B and D). These results support the validity of VOC-based classification of soybean developmental phases and further demonstrate the potential of this approach as a practical and non-invasive tool for monitoring phenological progression.

**Figure 5.**
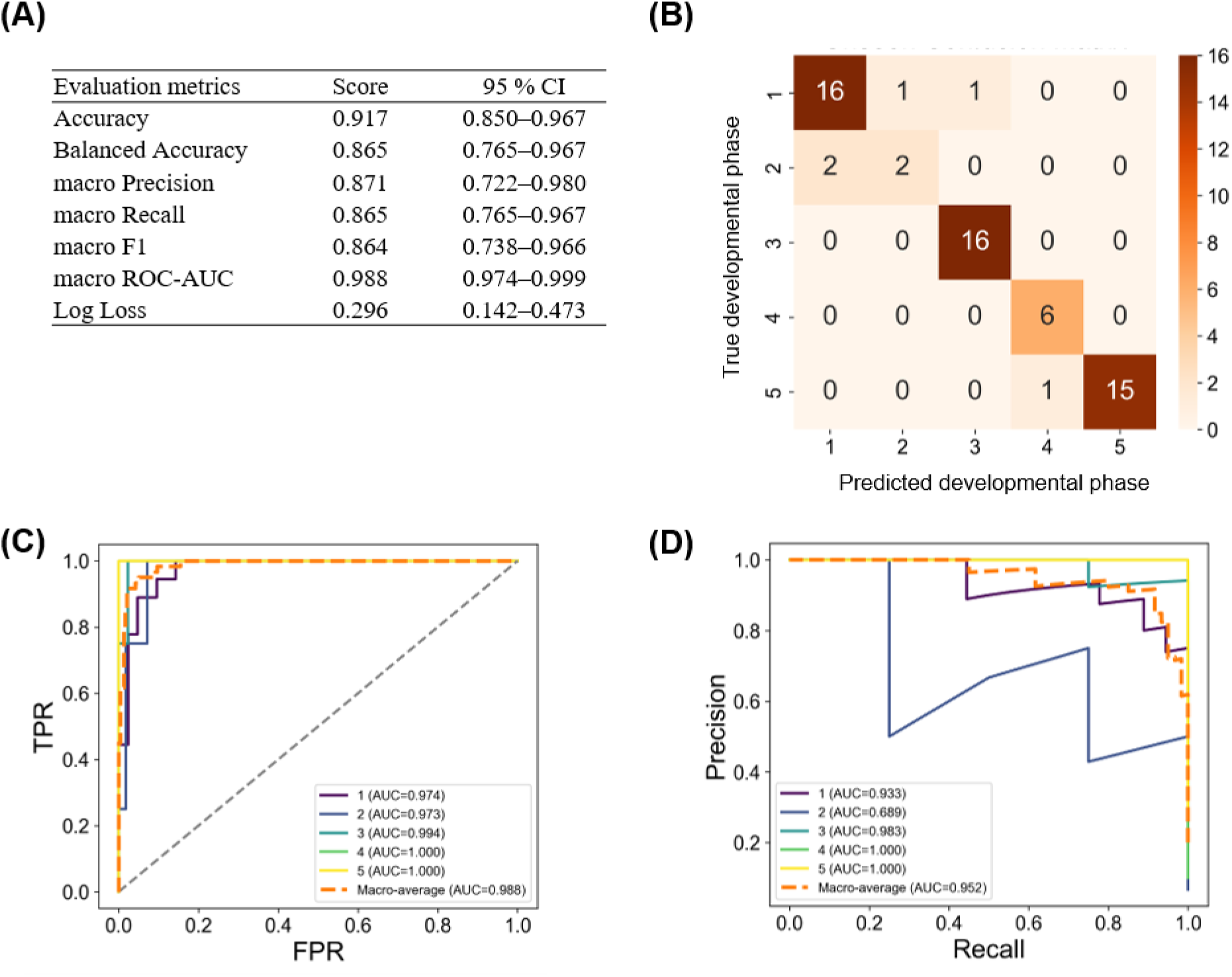
Predictive performances of the TabPFN model for soybean developmental phase classification using independent test data. (A) Evaluation metric scores with 95 % confidence intervals (95 % CI) from 1,000 bootstrap resamples. (B) Heatmap of the confusion matrix. (C) Receiver operating characteristic (ROC) curves. (D) Precision–Recall curves.

## Discussion

### Effectiveness of VOC-based phenological assessment in soybean

VOC-based approaches offer distinct advantages for phenological assessment in soybean. Because the period from flowering onset (R1) to seed filling (R6–R7) is a critical phase for yield determination (Vogel *et al*., 2021), stage-specific management practices and the development of cultivars adapted to local growing environments are particularly important. However, accurate assessment of phenological progression during this period remains challenging. As reproductive development proceeds, canopy closure can increasingly obscure flowers and developing pods formed at nodes throughout the plant. This makes conventional visual assessment during reproductive stages labour-intensive, error-prone, and potentially damaging to plants. Although optical sensing approaches are promising for identifying specific developmental stages (Sakamoto *et al*., 2010; Pan *et al*., 2023), continuous and non-invasive monitoring of phenological progression throughout the reproductive cycle remains challenging. By capturing physiological and metabolic changes that are not readily detectable visually, VOC profiling offers a valuable complementary approach. However, its application to continuous phenological monitoring remains underexploited, and the links between time-resolved VOC profiles and phenological events are poorly characterised.

In this study, we performed time-resolved VOC profiling of soybean from 16 to 43 DAS (V3–R5) using an automated TD-GC-MS platform, followed by untargeted data analysis (Figs. 1 and 2). Long-term GC-MS measurements are susceptible to instrumental drift, including retention time shifts and signal intensity variations, which can lead to erroneous peak detection, alignment, and integration (Koh *et al*., 2010; Watrous *et al*., 2017; Han and Li, 2020). To mitigate these effects, we combined statistical filtering with a network-based strategy inspired by feature-based molecular networking (Nothias *et al*., 2020). By integrating information on chemical relatedness and temporal co-variation, this framework expanded the feature set and enhanced biological interpretability, consistent with previous applications of network analysis in metabolomics (Perez de Souza *et al*., 2020; Amara *et al*., 2022). Using this approach, we obtained seven VOCs with clear temporal dynamics (Fig. 3), enabling the delineation of five developmental phases that were broadly consistent with conventional developmental stages (Fig. 4). Compared with analysis using only the three statistically retained VOCs, which produced only three coarse groups (Supplementary Fig. S6), the multi-step pipeline comprising statistical filtering and similarity- based network analysis provided finer resolution of developmental phases. These results indicate that our approach enable more sensitive detection of physiological changes associated with phenological progression. Furthermore, we developed a machine-learning model to predict these developmental phases based on the VOC profiles and achieved high predictive performance on an independent dataset (Fig. 5). Overall, our results demonstrate the effectiveness of combining VOC profiling with machine learning for accurate, time-resolved, and non-invasive assessment of soybean phenological progression.

### VOC profile dynamics reveal distinct phenological structure in soybean

We obtained seven soybean-emitted VOCs that exhibited distinct temporal dynamics during continuous sampling from 16 to 43 DAS, covering the transition from vegetative growth (V3) to reproductive development (R5). These VOCs were putatively associated with chemical groups including terpenes, fatty-acid derivatives, and furans, which have been commonly reported across plant species. Unlike studies focusing on specific developmental stages or narrowly defined periods, our analysis captured time-resolved VOC profile dynamics across an extended vegetative-to-reproductive developmental window. To the best of our knowledge, this study is among the first to explicitly link such continuous VOC dynamics to plant phenology. Previous studies have characterised VOC profiles in soybean seeds during late reproductive stages (R6–R8) (Boué *et al*., 2003), reported stage-specific emissions during anthesis in maize (Wiß *et al*., 2017), or observed stage-dependent VOC patterns in wheat, sorghum, and rice based on temporally sparse sampling (Batten *et al*., 1995; Hinge *et al*., 2016; Manco *et al*., 2021; Loubet *et al*., 2022). In contrast, our data-driven analysis of time-resolved VOC profiles resolved five distinct developmental phases across vegetative-to-reproductive progression (Fig. 4). Developmental phases 3, 4, and 5 corresponded closely to reproductive stages, R1–R3, R4, and R5, respectively. The correspondence between VOC-derived developmental phases and conventional developmental stages suggests that VOC profiles contain information relevant to established developmental transitions. Notably, the vegetative stage (V stage) was subdivided into two distinct phases (phases 1 and 2), with the latter corresponding to the onset of bud maturation (B1 stage) (Supplementary Fig. S4). This suggests that VOC profiles may capture subtle developmental features, which remain unresolved by conventional staging methods. Although budding-associated VOCs have been reported in species such as *Chrysanthemum indicum* (Zhu *et al*., 2022), comparable observations have not been described in soybean or other major crops. These results suggest that VOC profile dynamics reflect coordinated physiological changes across phenological progression. Unlike morphology-based conventional staging, which relies on visible developmental traits, VOC-based phenological assessment captures underlying physiological changes that are not directly represented by external morphology.

### Machine learning implementation for VOC-based developmental phase prediction

The concept that plant-emitted VOCs could serve as non-invasive biomarkers of plant physiological and environmental states has been recognised for decades (Niederbacher *et al*., 2015). Recent studies have demonstrated practical applications of VOC-based sensing, including disease detection in citrus (Aksenov *et al*., 2014), fruit maturity and cultivar classification in strawberry (Huang *et al*., 2025), water stress detection in soybean (Herrmann *et al*., 2024), mycotoxin contamination monitoring in wheat (Leggieri *et al*., 2022), and barley variety differentiation (Jud *et al*., 2018). In this study, we demonstrated that machine-learning models trained on VOC profiles could predict soybean developmental phases (Fig. 5). TabPFN represents a probabilistic transformer model pretrained on many synthetic tasks and can achieve high predictive performance on tabular data without extensive hyperparameter tuning specific to each task (Hollmann et al., 2025). This characteristic is well suited to VOC datasets obtained by GC/MS analysis, where sample sizes often remain limited due to the cost and complexity of the experiments. Consistent with this advantage, the TabPFN model achieved high predictive performance even without model-specific hyperparameter tuning, attaining an accuracy of 0.917 and a macro ROC-AUC of 0.988 on the independent test dataset (Fig. 5). Despite the high overall predictive performance, prediction of the underrepresented phase 2 remained comparatively less accurate (Fig. 5B and D), suggesting that class imbalance among developmental phases was not fully mitigated by KMeansSMOTE. This limitation is likely inherent to longitudinal phenological datasets, where developmental phases vary in duration and continuous sampling consequently yields uneven class representation. Further improvements would require more effective strategies for handling imbalanced data, including increased sampling of underrepresented phases, refinement of preprocessing and feature engineering, class-weighted or cost-sensitive learning, and ensemble strategies designed for imbalanced data (Krawczyk, 2016; Chen et al., 2024; Galar et al., 2012).

### Applicability and limitations of VOC-based phenological assessment

The findings of this study highlight the potential of VOC profiling for phenological assessment in soybean. The machine-learning model achieved high predictive performance on an independent dataset, indicating that VOC profiles contain developmentally informative physiological signals that may support decision-making in crop monitoring and management. Previous studies have demonstrated the potential of VOC profiling for diverse agricultural applications, including detecting water stress in soybean (Herrmann *et al*., 2024), assessing seed germination in maize and wheat (Zhang *et al*., 2022; Liu *et al*., 2024), monitoring postharvest quality in strawberry (Do *et al*., 2024), and identifying pest infestation in maple (Vermeeren *et al*., 2025). However, its application to continuous monitoring of phenological progression remains largely unexplored. The present study extends the utility of VOC profiling to phenological assessment, demonstrating its potential for non-invasive and time-resolved monitoring of phenological progression.

Integration of VOC sensing with other sensing modalities offers further opportunities for crop monitoring, particularly in combination with established optical sensing approaches. Optical and image-based methods have been used for phenological prediction in crops such as strawberry and wheat (Darlan *et al*., 2025; Naseer *et al*., 2025), and hyperspectral imaging has been widely applied to assess crop physiological and biochemical traits (Lu et al., 2020; Sarić et al., 2022). Combining VOC profiling with imaging and spectral sensing approaches could integrate morphological, spectral, and volatile chemical signals associated with crop developmental and physiological status. Such multimodal approaches are increasingly recognised as promising strategies for crop-state assessment because they can complement the strengths and compensate for the limitations of individual sensing modalities and improve the robustness of monitoring (Karmakar *et al*., 2024). Despite this potential, VOC sensing remains underutilised in multimodal frameworks for agricultural monitoring. Further integration of VOC-based approaches with established sensing technologies could contribute to the development of more sensitive, robust, and non-invasive systems for monitoring crop developmental and physiological status.

However, several limitations should be considered for broader application. First, the present study was conducted using a single soybean cultivar. As demonstrated in maize, barley, and Lamiaceae taxa, genetic background can influence VOC profiles (Ibrahim *et al*., 2022; Jud *et al*., 2018; Pretorius *et al*., 2022), suggesting that cultivar-specific variation should be incorporated into model development. Luo and Xu (2025) reported that VOC-based models achieved high accuracy for maturity assessment (96.7 %) and cultivar discrimination (98.3 %) using VOC profiles from multiple strawberry cultivars, suggesting that similar approaches could be extended to soybean and other crops. Second, environmental conditions, including temperature and humidity, can substantially affect VOC emissions (Niederbacher *et al*., 2015). Although the present study was conducted under controlled conditions, practical deployment will require analytical frameworks that remain robust to fluctuating environmental conditions (Gan *et al*., 2023). Third, field implementation will require VOC-sensing platforms that are portable, robust, and suitable for high-throughput monitoring. Technologies such as gas chromatography- differential mobility spectrometry (Aksenov *et al*., 2014) and field asymmetric ion mobility spectrometry (Zhang *et al*., 2023) provide promising options for rapid and portable VOC detection while retaining some degree of chemical separation or specificity. In contrast, odour-sensing platforms such as the electronic nose (eNose) generally capture overall VOC or odour patterns rather than resolving individual compounds, but may offer greater portability and lower cost option when combined with AI-driven analytical approaches (Seesaard *et al*., 2022; Marcillo *et al*., 2023; Haque *et al.,* 2026). Although sensor drift and reproducibility remain major challenges, continued advances in eNose technologies may facilitate the practical development of VOC-based crop monitoring systems in agricultural settings (Rabehi *et al*., 2024; Wörner *et al*., 2025).

As a proof-of-concept study, the present work focused on evaluating the feasibility of VOC-based phenological assessment rather than on definitive identification of VOC features or mechanistic analysis of VOC emission. Future studies using authentic standards for chemical identification, together with complementary omics approaches such as transcriptomics, metabolomics, or hormone profiling, would help clarify how VOC dynamics are linked to developmental regulation, metabolic reprogramming, and physiological transitions during phenological progression.

## Conclusion

In this study, we demonstrated that time-resolved VOC profiling combined with machine learning enables the accurate and non-invasive assessment of soybean phenological progression. By capturing temporal patterns in VOC emissions, the proposed framework extracts physiologically informative signals that complement conventional morphology-based staging and enable the detection of subtle developmental features, including early reproductive stages that are difficult to monitor visually. Although further validation across diverse genotypes and under field conditions will be required, these findings highlight the potential of VOC-based phenological assessment as a promising approach for crop monitoring. Continued advances in VOC sensing technology, together with integration into multimodal sensing frameworks, could further expand its application in plant phenotyping and crop management.

## Supplemental data

**Figure S1**: Network analysis based on time-series dynamics and structure relationships following statistical filtering.

**Figure S2**: Peak analysis performed using MassHunter software.

**Figure S3**: Data preprocessing and hyperparameter optimisation for community detection.

**Figure S4**: Time-course of bud numbers at or beyond the B1 stage.

**Figure S5**: Prediction performance of the TabPFN on validation data for the developmental phases.

**Figure S6**: Differentiation of developmental phase based on VOC profile derived from only the three features retained after statistical filtering.

**Table S1**: Software tools and spectral resources used for GC/MS data analysis.

**Table S2**: Detected communities and quality assessment in a VOC profile-based similarity network.

**Table S3**: Summary of developmental phase according to VOC profile and conventional staging. **Dataset S1**: Peak area values and related metadata for 250 alignment features detected across 300 records obtained from the MS-DIAL analysis.

**Dataset S2**: EI-MS spectra of 250 alignment features obtained from the MS-DIAL analysis. **Dataset S3**: Compound library file for the seven VOCs used in MRMPROBS peak curation. **Dataset S4**: Peak area values for the seven VOCs obtained from MRMPROBS analysis and related metadata across 300 records used to discriminate developmental phases through network analysis and community detection.

**Dataset S5**: Peak area values for the seven VOCs obtained from MRMPROBS analysis, related metadata, and corresponding class labels across 300 records used to build the predictive model with TabPFN.

**Dataset S6**: Peak area values for the seven VOCs obtained from MRMPROBS analysis and ground-truth class labels across 60 records, which served as previously unseen data for evaluating the predictive model built with TabPFN.

**Data S1**: Script and outputs for statistical filtering.

**Data S2**: Script and outputs for cosine similarity-based EI spectra similarity analysis.

**Data S3**: Script and outputs for Soft-DTW-based time-series similarity analysis.

**Data S4**: Script and outputs for multi-layer similarity network construction and subnetwork extraction using cosine and Soft-DTW similarity.

**Data S5**: Script and outputs for classifying soybean developmental phases using network analysis and community detection.

**Data S6**: Script and outputs for predicting soybean developmental phases from VOC profiles using TabPFN.

## Data availability

All data are included within the manuscript and supplementary material.

## Author contributions

RN, SH and MI: conceptualisation, funding acquisition and writing—original draft, and writing— review and editing. RN and SH: methodology and resources. RN: data curation, formal analysis, investigation, validation, and visualisation.

## AI Assistance

Microsoft Copilot was used to assist with English-language editing and improvement of manuscript readability. All scientific interpretations, conclusions, and final wording were reviewed and approved by the authors, who take full responsibility for the content of the manuscript.

## Acknowledgements

We thank Editage (www.editage.jp) for English language editing.

