## Supplementary material for "Time-resolved volatile organic compound profiling enables non-invasive detection of phenological progression in soybean": Supplemetal Materials: Supplementary Data S1.pdf

### Supplementary Data S1 The script and outputs for blank-based filtering.

This Python script implements a data preprocessing and statistical filtering pipeline for GC-MS peak area data. After importing all data (`All_Data.csv`), blank samples are identified where both `RepeatID` and `TimeID` equal 99, and are separated from non-blank samples. The script computes variable-wise mean values in blanks and filters compounds that exceed the upper blank threshold (blank mean × 5) in at least 50 samples and remain below the lower threshold (blank mean × 0.8) in less than 50 % of samples. The retained variables are then analyzed by a linear mixed model (LMM) including a fixed effect of `GroupID` and a random intercept per `RepeatID`, using only the overlapping time points (`TimeID` = 23 or 24). P-values for `GroupID` are adjusted using the Benjamini–Yekutieli method (FDR < 0.01), and significant variables are reported in `overlap_group_effect.csv`.

In [1]:

```
import warnings
import pandas as pd
import numpy as np
from statsmodels.formula.api import mixedlm
from statsmodels.stats.multitest import multipletests
from statsmodels.tools.sm_exceptions import ConvergenceWarning
warnings.filterwarnings("ignore", category=ConvergenceWarning)
```

In [2]:

```
# ------------------------------------------------------
# 1. Data Loading
# ------------------------------------------------------
# Read the CSV file and use all columns except the first two
data = pd.read_csv('All_Data.csv')
df = data.iloc[:, 2:]
print(f"Shape of df: {df.shape}")
```

```
Shape of df: (318, 253)
```

In [3]:

```
# ------------------------------------------------------
# 2. Extract and Save Blank and Non-Blank Data
# ------------------------------------------------------
# Blank data: records where both 'RepeatID' and 'TimeID' are 99
blank_condition = (df['RepeatID'] == 99) & (df['TimeID'] == 99)
blank_data = df[blank_condition].copy()
blank_data.to_csv('blank_data.csv', index=False)
print(f"Shape of blank_data: {blank_data.shape}")

# Non-blank data: records that do not satisfy the blank condition
non_blank_data = df[~blank_condition].copy()
non_blank_data.to_csv('non_blank_data.csv', index=False)
print(f"Shape of non_blank_data: {non_blank_data.shape}")
```

```
Shape of blank_data: (18, 253)
Shape of non_blank_data: (300, 253)
```

In [4]:

```
# ------------------------------------------------------
# 3. Filter Variables Based on Blank Data Criteria
# ------------------------------------------------------
# Parameter settings:
u = 5       # Multiplier for the blank mean for the upper threshold
n = 50       # Minimum number of values exceeding the upper threshold

l = 0.8     # Multiplier for the blank mean for the lower threshold
m = 50     # Maximum percentage of values below the lower threshold

# Target variables: columns from the 4th column onward in non_blank_data
variable_list = non_blank_data.iloc[:, 3:].columns.tolist()
# Compute the mean of the blank data for the target variables
blank_means = blank_data[variable_list].mean()

# Define thresholds: upper threshold = blank mean * u, lower threshold = blank mean * l
upper_thresholds = blank_means * u
lower_thresholds = blank_means * l

# Filter variables based on the criteria
filtered_vars = []
for var in variable_list:
    count_high = (non_blank_data[var] >= upper_thresholds[var]).sum()
    percentage_low = (non_blank_data[var] <= lower_thresholds[var]).mean() * 100
    if (count_high >= n) and (percentage_low < m):
        filtered_vars.append(var)

print(f"Filtered variables: {len(filtered_vars)} / {len(variable_list)}")
print(f"List of filtered variables: {filtered_vars}")

# Create and save the filtered non-blank data
filtered_non_blank_data = non_blank_data[['GroupID', 'RepeatID', 'TimeID'] + filtered_vars]
filtered_non_blank_data.to_csv('filtered_non_blank_data.csv', index=False)
```

```
Filtered variables: 11 / 250
List of filtered variables: ['AID_28', 'AID_55', 'AID_122', 'AID_128', 'AID_134', 'AID_142', 'AID_143', 'AID_149', 'AID_153', 'AID_154', 'AID_181']
```

In [5]:

```
# ------------------------------------------------------
# 4. LMM for Selecting Significant Variables (with random intercept and random slope for TimeID)
# ------------------------------------------------------
# Target variables: columns starting with "AID_"
variable_cols = [
    col for col in filtered_non_blank_data.columns
    if col.startswith('AID_')
]

# Analysis data: select records where TimeID is 23 or 24
overlap = filtered_non_blank_data[
    filtered_non_blank_data['TimeID'].isin([23, 24])
].copy()


# Convert 'GroupID' and 'RepeatID' to categorical type
```

```python
overlap.loc[:, 'GroupID']  = overlap['GroupID'].astype('category')
overlap.loc[:, 'RepeatID'] = overlap['RepeatID'].astype('category')

# Run LMM for each target variable
vars_ok   = []
p_group   = []

for var in variable_cols:
    formula = f"{var} ~ C(GroupID) + TimeID"
    try:
        m   = mixedlm(
            formula,
            overlap,
            groups=overlap["RepeatID"],
            re_formula="1"
        )
        res = m.fit()
        p = res.pvalues.get('C(GroupID)[T.2]', None)
        if p is not None:
            vars_ok.append(var)
            p_group.append(p)
        else:
            print(f"[WARN] {var}: p-value for GroupID not found")
    except Exception as e:
        print(f"[ERROR] {var}: {e}")

# Apply multiple testing correction using FDR (Benjamini-Hochberg method)
reject, p_group_bh, _, _ = multipletests(
    p_group, alpha=0.01, method='fdr_by'
)

results_overlap = pd.DataFrame({
    'Variable':   vars_ok,
    'p_group':    p_group,
    'p_group_BY': p_group_bh,
    'Reject':     reject
})
results_overlap.to_csv('overlap_group_effect.csv', index=False)

# Output variable names
keep_vars = results_overlap.loc[
    results_overlap['Reject'] == False,
    'Variable'
].tolist()

print(f"Variables: {len(keep_vars)} / {len(variable_cols)}")
print(keep_vars)
```

```
Variables: 3 / 11
['AID_28', 'AID_128', 'AID_153']
```
