## Supplementary material for "Time-resolved volatile organic compound profiling enables non-invasive detection of phenological progression in soybean": Supplemetal Materials: Supplementary Data S2.pdf

### Supplementary Data S2 The script and outputs for cosine similarity-based EI spectra similarity analysis.

This Python script provides a reproducible workflow for parsing GC–MS spectral files ( `All_spectra.msp` ) exported by MS-DIAL, normalizing peak intensities, binning mass spectra, and computing pairwise cosine similarity. The workflow consists of: (1) parsing `.msp` files line by line with error handling, extracting metadata and peak information, and saving the parsed spectra as a pickle file ( `All_spectra.pkl` ); (2) converting each spectrum to a normalized binned intensity vector using a defined bin width (0.3 m/z) and normalization method ( `max` , `l1` , or `l2` ); and (3) calculating a cosine similarity matrix among all spectra and visualizing the similarity structure as a heatmap. The resulting cosine similarity matrix is saved as `cosine_sim_max.csv` , and the visualization figure as `cosine_similarity_heatmap_max.png` .

In [1]:
```
import os
import time
import pickle
import pandas as pd
import numpy as np
import matplotlib.pyplot as plt
import seaborn as sns
from tqdm import tqdm
from sklearn.metrics.pairwise import cosine_similarity

plt.rcParams["font.family"] = "Arial"
```

In [2]:
```
# ----------------------------------------------------
# 1. Parse MSP File and Save as Pickle
# ----------------------------------------------------
msp_file_path = 'All_spectra.msp'
spectra = []              # List to store all spectrum entries
current_spectrum = {}     # Dictionary for the current spectrum entry
peaks_remaining = 0       # Number of peaks remaining for the current spectrum entry
filename = os.path.basename(msp_file_path)
value_error_count = 0     # Counter for ValueErrors
unrecognized_indices = [] # List to record indices with unrecognized line formats

start_time = time.perf_counter()

# Count total number of lines for progress tracking
with open(msp_file_path, 'r', encoding='utf-8') as f:
    total_lines = sum(1 for _ in f)

# Parse the MSP file line by line using tqdm for progress bar
with open(msp_file_path, 'r', encoding='utf-8') as f:
    for line_num, line in enumerate(tqdm(f, total=total_lines, desc="Parsing MSP file"), start=1):
        line = line.strip()
        if not line:
            # Empty line indicates the end of current spectrum entry.
            if current_spectrum:
                # Compute normalized intensities for peaks.
                spec = current_spectrum.get('Spectrum', [])
                intensities = [p['Intensity'] for p in spec]
                max_intensity = max(intensities) if intensities else 0
                sum_intensity = sum(intensities) if intensities else 0
                l1_norm = sum(abs(i) for i in intensities)
                l2_norm = np.sqrt(sum(i**2 for i in intensities)) if intensities else 0
                for peak in spec:
                    peak['Relative Intensity'] = peak['Intensity'] / max_intensity if max_intensity > 0 else 0
                    peak['Normalized Intensity'] = peak['Intensity'] / sum_intensity if sum_intensity > 0 else 0
                    peak['L1 Normalized Intensity'] = peak['Intensity'] / l1_norm if l1_norm > 0 else 0
                    peak['L2 Normalized Intensity'] = peak['Intensity'] / l2_norm if l2_norm > 0 else 0
                current_spectrum['filename'] = filename
                spectra.append(current_spectrum)
                current_spectrum = {}
            continue

        if peaks_remaining > 0:
            # Parse a peak line (should contain m/z and intensity).
            try:
                parts = line.split()
                if len(parts) < 2:
                    raise ValueError("Insufficient peak data.")
                mz = float(parts[0])
                intensity = float(parts[1])
                current_spectrum.setdefault('Spectrum', []).append({
                    'm/z': mz,
                    'Intensity': intensity
                })
                peaks_remaining -= 1
            except ValueError:
                value_error_count += 1
                current_index = len(spectra)
                if current_index not in unrecognized_indices:
                    unrecognized_indices.append(current_index)
        else:
            # Parse key-value line (e.g., NAME, NUM_PEAKS)
            if ':' in line:
                key, value = line.split(':', 1)
                key = key.strip().upper().replace(' ', '_')
                value = value.strip()
                if key == 'NAME':
                    parts = value.split('|')
                    current_spectrum['Name'] = parts[0].strip()
                    for part in parts[1:]:
                        if '=' in part:
                            sub_key, sub_value = part.split('=', 1)
                            sub_key = sub_key.strip().upper().replace(' ', '_')
                            sub_value = sub_value.strip()
                            try:
                                if sub_key == 'ID':
```

```python
                                    current_spectrum['ID'] = sub_value
                                else:
                                    current_spectrum[sub_key] = float(sub_value)
                            except ValueError:
                                current_spectrum[sub_key] = sub_value
                        else:
                            current_spectrum.setdefault('ADDITIONAL_INFO', '')
                            current_spectrum['ADDITIONAL_INFO'] += part + ' | '
                            current_spectrum['ADDITIONAL_INFO'] = current_spectrum['ADDITIONAL_INFO'].strip(' |')
                elif key == 'NUM_PEAKS':
                    try:
                        peaks_remaining = int(value)
                        current_spectrum['Num Peaks'] = peaks_remaining
                    except ValueError:
                        value_error_count += 1
                        current_index = len(spectra)
                        if current_index not in unrecognized_indices:
                            unrecognized_indices.append(current_index)
                else:
                    try:
                        current_spectrum[key] = float(value)
                    except ValueError:
                        current_spectrum[key] = value
            else:
                current_index = len(spectra)
                if current_index not in unrecognized_indices:
                    unrecognized_indices.append(current_index)

# Append the last spectrum entry if it exists.
if current_spectrum:
    spec = current_spectrum.get('Spectrum', [])
    intensities = [p['Intensity'] for p in spec]
    max_intensity = max(intensities) if intensities else 0
    sum_intensity = sum(intensities) if intensities else 0
    l1_norm = sum(abs(i) for i in intensities)
    l2_norm = np.sqrt(sum(i**2 for i in intensities)) if intensities else 0
    for peak in spec:
        peak['Relative Intensity'] = peak['Intensity'] / max_intensity if max_intensity > 0 else 0
        peak['Normalized Intensity'] = peak['Intensity'] / sum_intensity if sum_intensity > 0 else 0
        peak['L1 Normalized Intensity'] = peak['Intensity'] / l1_norm if l1_norm > 0 else 0
        peak['L2 Normalized Intensity'] = peak['Intensity'] / l2_norm if l2_norm > 0 else 0
    current_spectrum['filename'] = filename
    spectra.append(current_spectrum)

end_time = time.perf_counter()
elapsed_time = end_time - start_time
print(f"\nParsed spectra: {len(spectra)}")
print(f"Time taken: {elapsed_time:.2f} seconds")
print(f"ValueErrors: {value_error_count}")
if unrecognized_indices:
    print(f"Unrecognized format indices: {unrecognized_indices}")
else:
    print("No unrecognized line formats.")
print()

# Save parsed spectra to pickle
with open('All_spectra.pkl', 'wb') as f:
    pickle.dump(spectra, f)
```

```
Parsing MSP file: 100%|██████████████████████████████████████████████████████| 32557/32557 [00:00<00:00, 649406.52it/s]
Parsed spectra: 250
Time taken: 0.05 seconds
ValueErrors: 0
No unrecognized line formats.
```

In [3]:

```python
# -----------------------------------------------------
# 2. Load parsed spectra, bin spectra, and compute cosine similarity
# -----------------------------------------------------
with open('All_spectra.pkl', 'rb') as f:
    data = pickle.load(f)

# Binning parameters
bin_width = 0.3  # Width of each bin for m/z values
mz_min = 0       # Minimum m/z value
mz_max = 500     # Maximum m/z value
bins = np.arange(mz_min, mz_max + bin_width, bin_width)  # Create bin edges

# Select normalization method: 'max', 'l1', or 'l2'
normalization_method = 'max'

binned_spectra = []  # List to store binned intensity vectors for each spectrum
spectra_ids = []     # List for sequential spectrum IDs
spectra_names = []   # List for spectrum names

# For each spectrum entry, create a binned intensity vector
for idx, entry in enumerate(data):
    spectra_ids.append(idx)
    spectrum = entry.get('Spectrum', [])
    name = entry.get('Name', f"Spectrum_{idx}")
    spectra_names.append(name)

    # Get the list of intensities from the spectrum
    intensities = [peak['Intensity'] for peak in spectrum]
    if intensities:
        if normalization_method == 'max':
            norm_factor = max(intensities)
        elif normalization_method == 'l1':
            norm_factor = sum(abs(i) for i in intensities)
        elif normalization_method == 'l2':
            norm_factor = np.sqrt(sum(i**2 for i in intensities))
```

```python
        else:
            raise ValueError("Invalid normalization method. Choose 'max', 'l1', or 'l2'.")
    else:
        norm_factor = 1  # Avoid division by zero

    # Initialize a vector for binned intensities
    binned_intensity = np.zeros(len(bins) - 1)
    # For each peak, determine the appropriate bin and add the normalized intensity
    for peak in spectrum:
        mz = peak['m/z']
        intensity = peak['Intensity'] / norm_factor
        if mz_min <= mz < mz_max:
            bin_index = np.searchsorted(bins, mz, side='right') - 1
            binned_intensity[bin_index] += intensity
    binned_spectra.append(binned_intensity)

# Convert the list of binned spectra to a 2D numpy array
binned_matrix = np.array(binned_spectra)
# Compute the cosine similarity matrix among the binned spectra
cosine_sim_matrix = cosine_similarity(binned_matrix)
# Convert the similarity matrix to a DataFrame for better visualization
cosine_sim_df = pd.DataFrame(cosine_sim_matrix, index=spectra_ids, columns=spectra_ids)

# Save the cosine similarity matrix as a CSV file and display the first few rows
cosine_sim_df.to_csv(f"cosine_sim_{normalization_method}.csv", index=True)
```

In [4]:

```python
# ----------------------------------------------------
# 3. Visualize the Cosine Similarity Matrix as a Heatmap
# ----------------------------------------------------
# Set up the matplotlib figure
plt.figure(figsize=(10, 8))

# Plot the heatmap using seaborn
# We use a diverging colormap and display the similarity values.
sns.heatmap(cosine_sim_df,
            cmap='Blues',
            annot=False,
            fmt=".2f",
            cbar_kws={"label": "Cosine Similarity", "shrink": 0.8})

plt.xlabel("Spectrum ID")
plt.ylabel("Spectrum ID")
plt.tight_layout()
plt.savefig(f"cosine_similarity_heatmap_{normalization_method}.png", dpi=300, bbox_inches='tight')
plt.show()
```

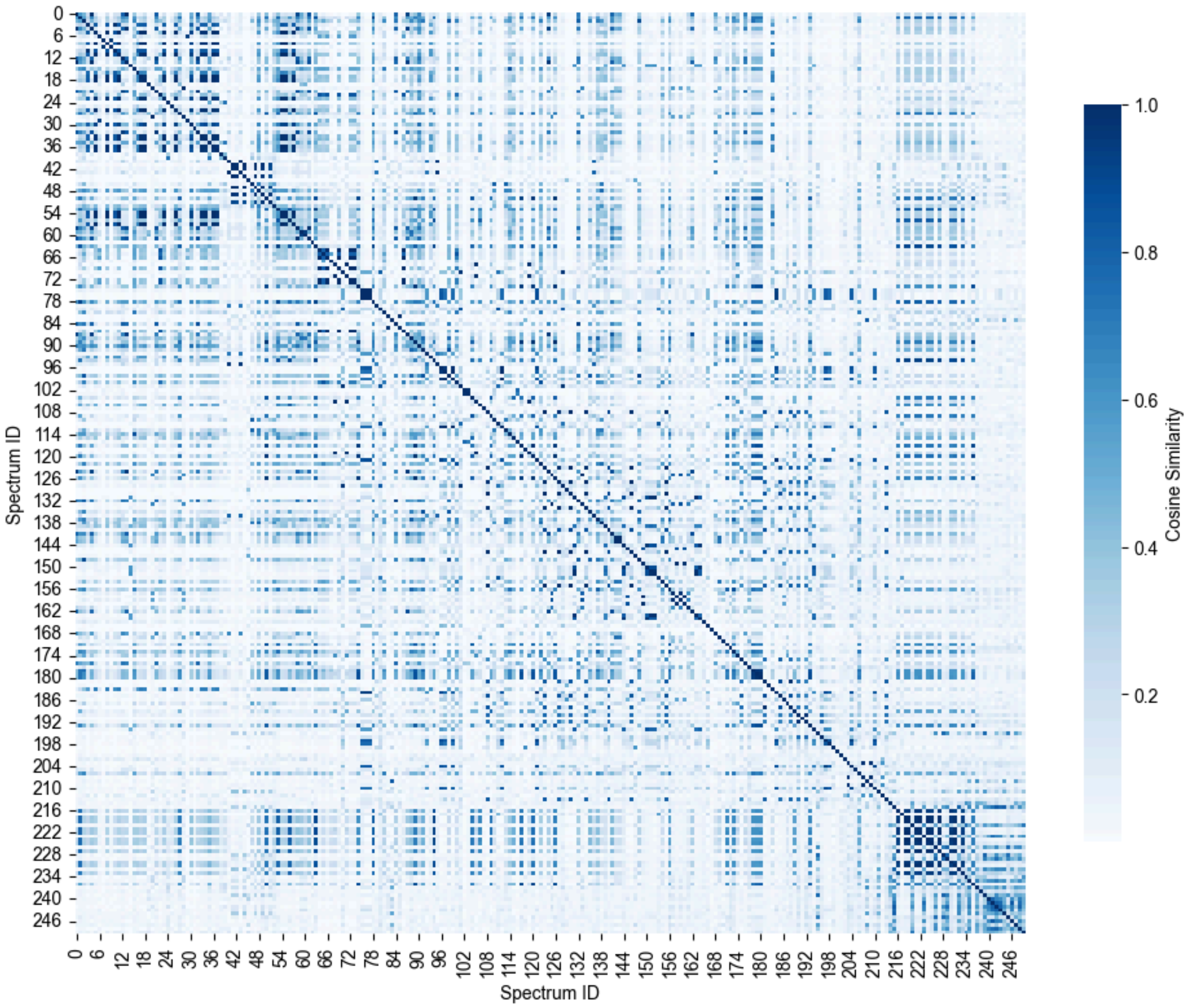
