## Supplementary material for "Time-resolved volatile organic compound profiling enables non-invasive detection of phenological progression in soybean": Supplemetal Materials: Supplementary Data S3.pdf

### Supplementary Data S3 The script and outputs for Soft-DTW-based time-series similarity analysis

This Python script provides a reproducible workflow for calculating pairwise time-series similarity among GC–MS-derived variables using the Soft Dynamic Time Warping (Soft-DTW) algorithm. The pipeline consists of: (1) loading the dataset ( `All_data.csv` ) and removing blank samples where both `RepeatID` and `TimeID` equal 99; (2) extracting variables beginning with `AID_` and computing mean time-series values across `TimeID` ; (3) standardizing each variable's mean trajectory using z-score normalization; and (4) calculating pairwise Soft-DTW distances (γ = 0.01) in parallel across all variable pairs; and The computed distance matrix is saved as `Soft_DTW_distance_matrix_gamma_0.01.csv` , and the corresponding heatmap figure as `soft_dtw_distance_heatmap_gamma_0.01.png` .

In [1]:
```
import os
import itertools
import time
import pickle
import warnings
warnings.filterwarnings("ignore", category=UserWarning)

import pandas as pd
import numpy as np
from sklearn.preprocessing import StandardScaler
from tslearn.metrics import soft_dtw
from joblib import Parallel, delayed
from tqdm.auto import tqdm
# from tqdm_joblib import tqdm_joblib
from pathlib import Path
import matplotlib.pyplot as plt
import matplotlib.colors as mcolors
import matplotlib.cm as cm
import seaborn as sns
import math

plt.rcParams["font.family"] = "Arial"
```

```
D:\miniconda3\envs\tabpfn-gpu\Lib\site-packages\tqdm\auto.py:21: TqdmWarning: IProgress not found. Please update jupyter and ipywidgets. See https://ipywi
dgets.readthedocs.io/en/stable/user_install.html
  from .autonotebook import tqdm as notebook_tqdm
```

In [2]:
```
# ------------------------------------------------------
# 1. Data Loading and Preprocessing
# ------------------------------------------------------
# Read the CSV file and create a DataFrame with all columns.
data = pd.read_csv('All_data.csv')
df = data.copy()
print(f"\nDataFrame shape: {df.shape}")

# Remove rows where both 'RepeatID' and 'TimeID' equal 99 (these are considered blank data)
if 'RepeatID' not in df.columns or 'TimeID' not in df.columns:
    raise KeyError("Columns 'RepeatID' and/or 'TimeID' are missing.")
df_cleaned = df[~((df['RepeatID'] == 99) & (df['TimeID'] == 99))].reset_index(drop=True)
print(f"\nShape after cleanup: {df_cleaned.shape}")
```

```
DataFrame shape: (318, 255)

Shape after cleanup: (300, 255)
```

In [3]:
```
# ------------------------------------------------------
# 2. Extract Variables and Time/Repeat Information
# ------------------------------------------------------
# Extract columns starting with 'AID_' which are the variables to analyze.
variables = [col for col in df_cleaned.columns if col.startswith('AID_')]
num_variables = len(variables)
print(f"\nNumber of variables: {num_variables}")

# Extract unique TimeID and RepeatID values.
time_ids = sorted(df_cleaned['TimeID'].unique())
repeats = sorted(df_cleaned['RepeatID'].unique())
print(f"\nNumber of time points: {len(time_ids)}")
print(f"Number of repeats: {len(repeats)}")
```

```
Number of variables: 250

Number of time points: 28
Number of repeats: 22
```

In [4]:
```
# ------------------------------------------------------
# 3. Compute and Standardize Mean Time Series for Each Variable
# ------------------------------------------------------
# For each variable, compute the mean time series across TimeID and then standardize it.
mean_time_series_dict = {}
for var in tqdm(variables, desc="Calculating mean time series"):
    mean_series = df_cleaned.groupby('TimeID')[var].mean().reindex(time_ids).values
    scaler = StandardScaler()
    standardized_series = scaler.fit_transform(mean_series.reshape(-1, 1)).flatten()
    mean_time_series_dict[var] = standardized_series
```

```
Calculating mean time series: 100%|██████████████████████████████████████████████████| 250/250 [00:00<00:00, 2497.96it/s]
```

In [5]:
```
# ------------------------------------------------------
# 4. Compute Soft-DTW Distance Matrix
# ------------------------------------------------------
JOBLIB_TMP = r"C:\joblib_temp"
Path(JOBLIB_TMP).mkdir(parents=True, exist_ok=True)

# Set gamma parameter for soft-DTW.
gamma = 0.01

# Function to compute soft-DTW distance between two time series.
def compute_soft_dtw(pair, variables, time_series_dict, gamma):
    i, j = pair
```

```python
    ts_i = time_series_dict[variables[i]]
    ts_j = time_series_dict[variables[j]]
    return soft_dtw(ts_i, ts_j, gamma=gamma)

# Generate all combinations with replacement of variable indices.
pairs = list(itertools.combinations_with_replacement(range(num_variables), 2))
dtw_distance_matrix = np.zeros((num_variables, num_variables))

# Compute soft-DTW distances in parallel and show progress using tqdm.
dtw_results = Parallel(n_jobs=-1, backend="loky", temp_folder=JOBLIB_TMP, max_nbytes=None)(
    delayed(compute_soft_dtw)(pair, variables, mean_time_series_dict, gamma)
    for pair in tqdm(pairs, desc="Computing Soft-DTW distances")
)

# Fill the symmetric distance matrix.
for idx, (i, j) in enumerate(pairs):
    distance = dtw_results[idx]
    dtw_distance_matrix[i, j] = distance
    dtw_distance_matrix[j, i] = distance
np.fill_diagonal(dtw_distance_matrix, 0)

# Convert the distance matrix to a DataFrame and save as CSV.
dtw_df = pd.DataFrame(dtw_distance_matrix, index=range(num_variables), columns=range(num_variables))
output_filename = f"Soft_DTW_distance_matrix_gamma_{gamma}.csv"
dtw_df.to_csv(output_filename, index=True)
```

```
Computing Soft-DTW distances: 100%|████████████████████████████████████████| 31375/31375 [00:11<00:00, 2772.57it/s]
```

In [6]:

```python
# ----------------------------------------------------
# 5. Visualize Soft-DTW Distance Matrix as a Heatmap
# ----------------------------------------------------
# Create a figure for the heatmap
plt.figure(figsize=(10, 8))

# Plot the heatmap for the Soft-DTW distance matrix using a red colormap
sns.heatmap(dtw_df,
            cmap='Reds',        # Red colormap for distances
            annot=False,        # Do not annotate cells with values
            fmt=".2f",          # Format numbers with 2 decimal places
            cbar_kws={"label": "Soft-DTW Distance", "shrink": 0.8})

plt.xlabel("Variable Index")
plt.ylabel("Variable Index")
plt.tight_layout()
plt.savefig(f"soft_dtw_distance_heatmap_gamma_{gamma}.png", dpi=300, bbox_inches='tight')
plt.show()
```

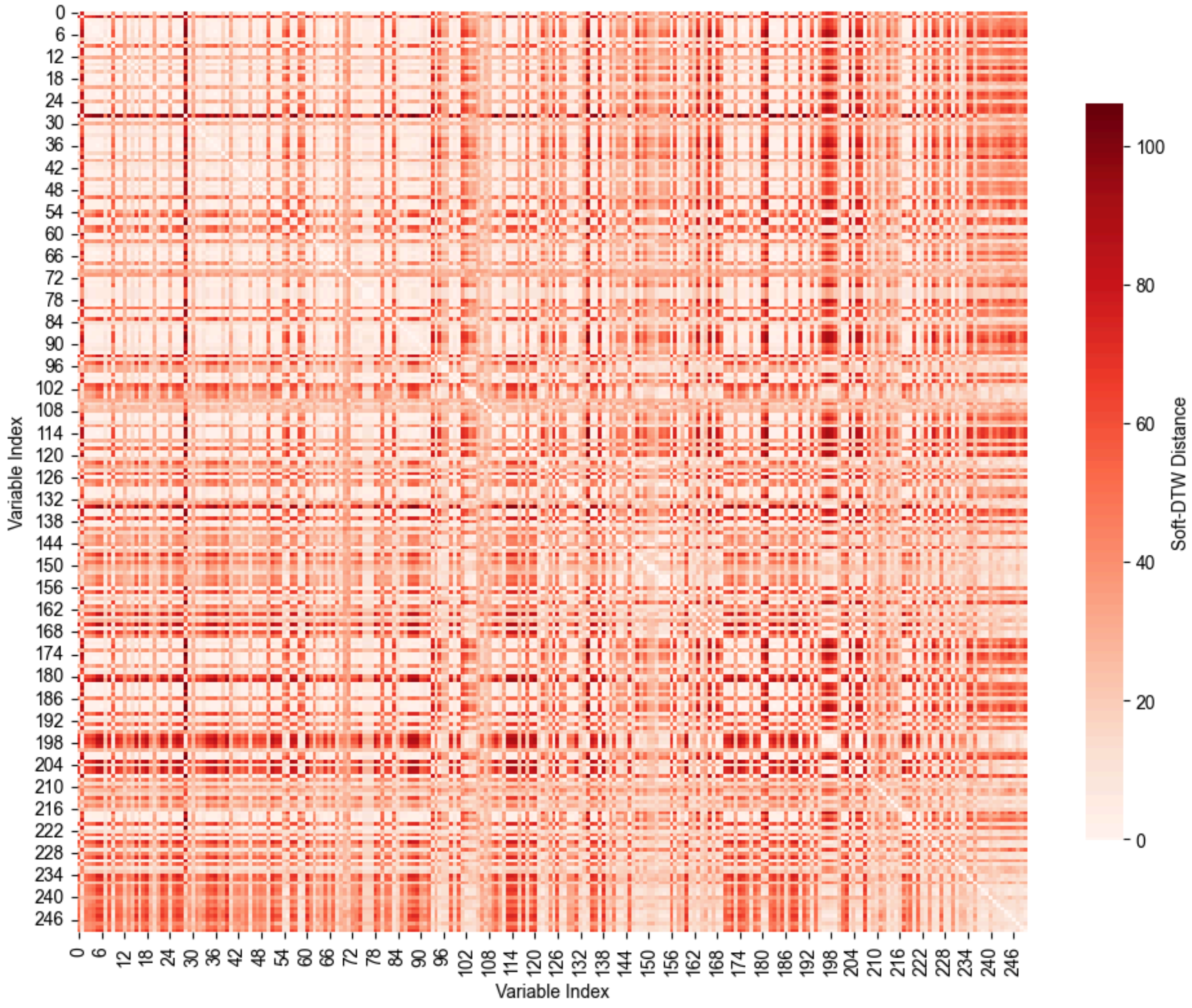
