## Supplementary material for "Time-resolved volatile organic compound profiling enables non-invasive detection of phenological progression in soybean": Supplemetal Materials: Supplementary Data S4.pdf

### Supplementary Data S4 The script and outputs for multi-layer similarity network construction and subnetwork extraction using cosine and Soft-DTW similarity

This Python script builds a multi-layer similarity network by integrating Soft-DTW–based temporal similarity and cosine spectral similarity, and then extracts a size-filtered subnetwork around pre-specified core nodes. The workflow comprises: (1) computing node "Size_Ratio" from `All_Data.csv` as the mean of non-baseline samples (rows with `RepeatID` & `TimeID` ≠ 99) divided by the baseline mean (rows with `RepeatID` & `TimeID` = 99), saved as `node_sizes.csv`; (2) loading Soft-DTW distances (`Soft_DTW_distance_matrix_gamma_0.01.csv`), converting them to similarities via an RBF kernel, and thresholding at 87.5%; (3) loading cosine similarities (`cosine_sim_max.csv`) and applying a fixed threshold of 0.60; (4) visualizing similarity distributions as histograms (`soft_dtw_histogram.png`, `cosine_similarity_histogram.png`);(5) constructing a multi-layer network (Soft-DTW layer in red, cosine layer in blue), highlighting user-defined core nodes (`CORE_NODE_IDS = [28, 128, 153]`) and their one-hop neighbors across layers; and (6) extracting a subnetwork consisting of the union of core nodes and their one-hop neighbors filtered by `Size_Ratio ≥ 5`, exporting node/edge lists (`n_hop_nodes_multi.csv`, `n_hop_edges_multi.csv`) and figures of the full network and subnetwork (`multi_layer_network.png`, `subnetwork.png`).

In [1]:
```
import numpy as np
import pandas as pd
import matplotlib.pyplot as plt
import networkx as nx
from matplotlib.colors import Normalize
from matplotlib.lines import Line2D
import math

plt.rcParams["font.family"] = "Arial"
```

In [2]:
```
# ----------------------------------------
# 1. Parameter Settings
# ----------------------------------------
# Imput files
INPUT_CSV_SIZES = 'All_Data.csv'
INPUT_CSV_DTW_DISTANCE = 'Soft_DTW_distance_matrix_gamma_0.01.csv'
INPUT_CSV_COSINE_SIMILARITY = 'cosine_sim_max.csv'

# Core Node setting
CORE_NODE_IDS = [28, 128, 153]
```

In [3]:
```
# ----------------------------------------
# 2. Node Size Calculation (Calculate Size_Ratio)
# ----------------------------------------
# For each column starting with "AID_", compute the ratio of the mean of non-baseline
# rows (RepeatID & TimeID != 99) to the mean of baseline rows (RepeatID & TimeID == 99).
# This ratio (Size_Ratio) is later used as a filtering criterion for subnetwork extraction.
OUTPUT_CSV_NODE_SIZES = 'node_sizes.csv'
size_df = pd.read_csv(INPUT_CSV_SIZES)
aid_columns = [col for col in size_df.columns if col.startswith('AID_')]

node_sizes = []
for aid in aid_columns:
    # Extract node ID from column name (e.g., "AID_123" -> 123)
    node_id = int(aid.split('_')[1])
    # Define baseline: rows where RepeatID and TimeID equal 99
    baseline_mask = (size_df['RepeatID'] == 99) & (size_df['TimeID'] == 99)
    baseline_mean = size_df.loc[baseline_mask, aid].mean()
    # Define non-baseline: all other rows
    non_baseline_mask = ~baseline_mask
    non_baseline_mean = size_df.loc[non_baseline_mask, aid].mean()
    # Calculate Size_Ratio; if baseline_mean is zero, use 1 as default
    size_ratio = non_baseline_mean / baseline_mean if baseline_mean != 0 else 1
    node_sizes.append({'Node_ID': node_id, 'Size_Ratio': size_ratio})

node_sizes_df = pd.DataFrame(node_sizes)
node_sizes_df.to_csv(OUTPUT_CSV_NODE_SIZES, index=False, encoding='utf-8-sig')

# Create a dictionary for later filtering (subnetwork extraction)
node_size_dict = pd.Series(node_sizes_df.Size_Ratio.values, index=node_sizes_df.Node_ID).to_dict()
```

In [4]:
```
# ----------------------------------------
# 3. Load Similarity Matrices and Compute Thresholds
# ----------------------------------------
# Read the Soft-DTW distance matrix and cosine similarity matrix,
# convert distances to similarities, and determine thresholds for edge inclusion.
#
# --- Soft-DTW Similarity ---
dtw_distance_df = pd.read_csv(INPUT_CSV_DTW_DISTANCE, index_col=0)
# Convert indices and columns to integers
dtw_distance_df.index = dtw_distance_df.index.astype(int)
dtw_distance_df.columns = dtw_distance_df.columns.astype(int)
spectra_ids = dtw_distance_df.index.tolist()

distance_matrix = dtw_distance_df.values
# Shift the distance matrix so that the minimum becomes 0
shifted_distance = distance_matrix - distance_matrix.min()
# Compute sigma as the median of the off-diagonal elements
mask = ~np.eye(dtw_distance_df.shape[0], dtype=bool)
sigma = np.median(shifted_distance[mask])

# Compute similarity matrix using an RBF kernel; set diagonal to 1
similarity_matrix_dtw = np.exp(- (shifted_distance ** 2) / (2 * sigma ** 2))
np.fill_diagonal(similarity_matrix_dtw, 1.0)
similarity_df_dtw = pd.DataFrame(similarity_matrix_dtw, index=spectra_ids, columns=spectra_ids)
# Flatten the similarity values (excluding self-similarity)
flattened_dtw = similarity_df_dtw.values.flatten()
flattened_dtw = flattened_dtw[flattened_dtw != 1.0]
# Set threshold at n%
threshold_sim_dtw = np.percentile(flattened_dtw, 87.5)
selected_count_dtw = np.sum(flattened_dtw >= threshold_sim_dtw)
```

```
percentage_selected_dtw = (selected_count_dtw / len(flattened_dtw)) * 100
print(f"Soft-DTW Threshold: {threshold_sim_dtw:.4f}, Selected: {percentage_selected_dtw:.2f}%")

# --- Cosine Similarity ---
cosine_similarity_df = pd.read_csv(INPUT_CSV_COSINE_SIMILARITY, index_col=0)
# Ensure the matrix is square
if cosine_similarity_df.shape[0] != cosine_similarity_df.shape[1]:
    raise ValueError("Cosine similarity matrix must be square.")
cosine_similarity_df.index = cosine_similarity_df.index.astype(int)
cosine_similarity_df.columns = cosine_similarity_df.columns.astype(int)
cosine_ids = cosine_similarity_df.index.tolist()
# Set self-similarity to 1
np.fill_diagonal(cosine_similarity_df.values, 1.0)
flattened_cos = cosine_similarity_df.values.flatten()
flattened_cos = flattened_cos[flattened_cos != 1.0]
# Use a fixed threshold for cosine similarity
threshold_cos = 0.6
selected_count_cos = np.sum(flattened_cos >= threshold_cos)
percentage_selected_cos = (selected_count_cos / len(flattened_cos)) * 100
print(f"Cosine Threshold: {threshold_cos:.4f}, Selected: {percentage_selected_cos:.2f}%")
```

```
Soft-DTW Threshold: 0.9870, Selected: 12.50%
Cosine Threshold: 0.6000, Selected: 4.51%
```

In [5]:
```
# ---------------------------------------
# 4. Plot Similarity Histograms
# ---------------------------------------
# Visualize the distribution of similarity values for both Soft-DTW and Cosine similarity.
OUTPUT_PNG_HISTOGRAM_DTW = 'soft_dtw_histogram.png'
fig_hist_dtw, ax_hist_dtw = plt.subplots(figsize=(6, 3))
ax_hist_dtw.hist(flattened_dtw, bins=50, color='lightcoral', edgecolor='black')
ax_hist_dtw.axvline(x=threshold_sim_dtw, color='red', linestyle='--',
                    label=f"Threshold: {threshold_sim_dtw:.4f}\nSelected: {percentage_selected_dtw:.2f}%")
ax_hist_dtw.set_title("Soft-DTW Similarity Histogram")
ax_hist_dtw.set_xlabel("Similarity")
ax_hist_dtw.set_ylabel("Frequency")
ax_hist_dtw.legend(loc='upper left', bbox_to_anchor=(1, 1), frameon=False)
plt.tight_layout()
plt.savefig(OUTPUT_PNG_HISTOGRAM_DTW, dpi=300, bbox_inches='tight')
plt.show()
plt.close()

OUTPUT_PNG_HISTOGRAM_COS = 'cosine_similarity_histogram.png'
fig_hist_cos, ax_hist_cos = plt.subplots(figsize=(6, 3))
ax_hist_cos.hist(flattened_cos, bins=50, color='lightblue', edgecolor='black')
ax_hist_cos.axvline(x=threshold_cos, color='blue', linestyle='--',
                    label=f"Threshold: {threshold_cos:.4f}\nSelected: {percentage_selected_cos:.2f}%")
ax_hist_cos.set_title("Cosine Similarity Histogram")
ax_hist_cos.set_xlabel("Similarity")
ax_hist_cos.set_ylabel("Frequency")
ax_hist_cos.legend(loc='upper left', bbox_to_anchor=(1, 1), frameon=False)
plt.tight_layout()
plt.savefig(OUTPUT_PNG_HISTOGRAM_COS, dpi=300, bbox_inches='tight')
plt.show()
plt.close()
```

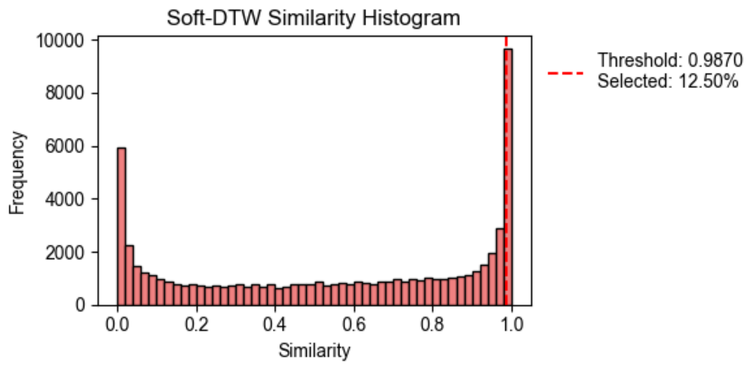


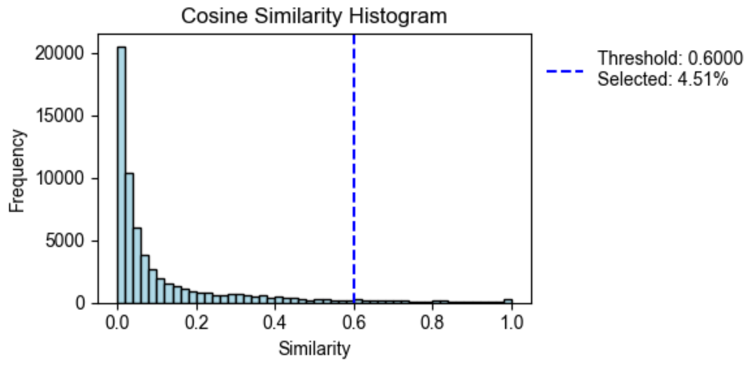


In [6]:
```
# ---------------------------------------
# 5. Build Multi-Layer Network
# ---------------------------------------
# Construct a network graph by adding edges for pairs of nodes
# if their similarity meets the defined thresholds.
OUTPUT_PNG_NETWORK = 'multi_layer_network.png'
# Define edge drawing parameters inline.
CMAP_DTW = plt.cm.Reds
CMAP_COS = plt.cm.Blues
EDGE_WIDTH = 1
```

```python
EDGE_ALPHA = 1

all_nodes = set(spectra_ids).union(set(cosine_ids))
G_multi = nx.Graph()
G_multi.add_nodes_from(all_nodes)

# Add edges based on Soft-DTW similarity threshold.
for i in spectra_ids:
    for j in spectra_ids:
        if i < j:
            sim_value = similarity_df_dtw.at[i, j]
            if sim_value >= threshold_sim_dtw:
                G_multi.add_edge(i, j, Soft_DTW=sim_value)

# Add edges based on Cosine similarity threshold.
for i in cosine_ids:
    for j in cosine_ids:
        if i < j:
            sim_value = cosine_similarity_df.at[i, j]
            if sim_value >= threshold_cos:
                if G_multi.has_edge(i, j):
                    G_multi.edges[i, j]['Cosine'] = sim_value
                else:
                    G_multi.add_edge(i, j, Cosine=sim_value)

print(f"Multi-layer network: {G_multi.number_of_nodes()} nodes, {G_multi.number_of_edges()} edges.")
```

Multi-layer network: 250 nodes, 4992 edges.

In [7]:
```python
# ----------------------------------------
# 6. Determine Core Nodes and 1-Hop Neighbors
# ----------------------------------------
# Identify core nodes (from CORE_NODE_IDS) that are present in the network,
# and then extract their 1-hop neighbors based on both Soft-DTW and Cosine edges.
# Define node color parameters inline.
CORE_NODE_COLOR = 'gold'
ONE_HOP_DTW_COLOR = '#FF6666'
ONE_HOP_COS_COLOR = '#66B2FF'
BOTH_HOP_COLOR = 'plum'
OTHER_NODES_COLOR = '#A9A9A9'

G_core = [node for node in CORE_NODE_IDS if node in spectra_ids and node in cosine_ids]
print("Core node IDs:", G_core)

# Extract 1-hop neighbors using Soft-DTW edges.
G_dtw = nx.Graph((u, v, d) for u, v, d in G_multi.edges(data=True) if 'Soft_DTW' in d)
n_hop_nodes_dtw = set()
for core in G_core:
    if core in G_dtw:
        n_hop_nodes_dtw.update(nx.single_source_shortest_path_length(G_dtw, core, cutoff=1).keys())

# Extract 1-hop neighbors using Cosine edges.
G_cos = nx.Graph((u, v, d) for u, v, d in G_multi.edges(data=True) if 'Cosine' in d)
n_hop_nodes_cos = set()
for core in G_core:
    if core in G_cos:
        n_hop_nodes_cos.update(nx.single_source_shortest_path_length(G_cos, core, cutoff=1).keys())
```

Core node IDs: [28, 128, 153]

In [8]:
```python
# ----------------------------------------
# 7. Subnetwork Extraction
# ----------------------------------------
# From the union of core nodes and their 1-hop neighbors,
# include only nodes with Size_Ratio >= 5.
candidate_nodes = set(G_core).union(n_hop_nodes_dtw).union(n_hop_nodes_cos)
subnetwork_nodes = {node for node in candidate_nodes if (node in G_core or node_size_dict.get(node, 0) >= 5)}
print(f"Subnetwork: {len(subnetwork_nodes)} nodes.")

G_sub = G_multi.subgraph(subnetwork_nodes).copy()
subnetwork_node_names = sorted(list(G_sub.nodes()))
print("Subnetwork node names:", subnetwork_node_names)

# Save subnetwork nodes and edges to CSV.
OUTPUT_CSV_SUBNETWORK_NODES = 'n_hop_nodes_multi.csv'
df_sub_nodes = pd.DataFrame({'Node_ID': sorted(list(subnetwork_nodes))})
df_sub_nodes.to_csv(OUTPUT_CSV_SUBNETWORK_NODES, index=False, encoding='utf-8-sig')

OUTPUT_CSV_SUBNETWORK_EDGES = 'n_hop_edges_multi.csv'
subnetwork_edges = []
for u, v, d in G_multi.edges(data=True):
    if u in subnetwork_nodes and v in subnetwork_nodes:
        subnetwork_edges.append({
            'Node1_ID': u,
            'Node2_ID': v,
            'Soft_DTW_Similarity': d.get('Soft_DTW', np.nan),
            'Cosine_Similarity': d.get('Cosine', np.nan)
        })
df_sub_edges = pd.DataFrame(subnetwork_edges)
df_sub_edges.to_csv(OUTPUT_CSV_SUBNETWORK_EDGES, index=False, encoding='utf-8-sig')
```

Subnetwork: 7 nodes.
Subnetwork node names: [28, 122, 128, 142, 143, 153, 154]

In [9]:
```python
# ----------------------------------------
# 8. Main Network Visualization
# ----------------------------------------
# Visualize the full multi-layer network with fixed node sizes and color coding.
# Compute a fixed layout.
pos = nx.spring_layout(G_multi, k=0.8, iterations=150, seed=42)

# Define fixed drawing parameters inline.
```

```python
FIXED_NODE_SIZE = 175
FIXED_NODE_EDGE_COLOR = 'black'
FIXED_NODE_EDGE_WIDTH = 0.8
NODE_LABEL_FONT_SIZE = 7
SHOW_NODE_LABELS = True
NODE_ALPHA = 0.60

node_colors = []
for node in G_multi.nodes():
    if node in G_core:
        node_colors.append(CORE_NODE_COLOR)
    elif node in n_hop_nodes_dtw and node in n_hop_nodes_cos:
        node_colors.append(BOTH_HOP_COLOR)
    elif node in n_hop_nodes_dtw:
        node_colors.append(ONE_HOP_DTW_COLOR)
    elif node in n_hop_nodes_cos:
        node_colors.append(ONE_HOP_COS_COLOR)
    else:
        node_colors.append(OTHER_NODES_COLOR)

fig = plt.figure(figsize=(12, 12))
ax_main = fig.add_axes([0.05, 0.01, 0.60, 0.5])
ax_cbar_dtw = fig.add_axes([0.68, 0.01, 0.015, 0.35])
ax_cbar_cos = fig.add_axes([0.76, 0.01, 0.015, 0.35])
ax_legend = fig.add_axes([0.6, 0.45, 0.25, 0.05])
ax_legend.axis('off')

# Draw nodes with fixed size and edge color.
nx.draw_networkx_nodes(G_multi, pos, node_size=FIXED_NODE_SIZE, node_color=node_colors,
                       edgecolors=FIXED_NODE_EDGE_COLOR, linewidths=FIXED_NODE_EDGE_WIDTH,
                       alpha=NODE_ALPHA, ax=ax_main)

# Draw Soft-DTW edges.
dtw_edges = [(u, v) for u, v, d in G_multi.edges(data=True) if 'Soft_DTW' in d]
dtw_weights = [G_multi.edges[u, v]['Soft_DTW'] for u, v in dtw_edges]
nx.draw_networkx_edges(G_multi, pos, edgelist=dtw_edges, edge_color=dtw_weights, edge_cmap=CMAP_DTW,
                       edge_vmin=threshold_sim_dtw, edge_vmax=similarity_df_dtw.values.max(),
                       width=EDGE_WIDTH, alpha=EDGE_ALPHA, style=(0, (2, 2)), ax=ax_main)

# Draw Cosine edges.
cos_edges = [(u, v) for u, v, d in G_multi.edges(data=True) if 'Cosine' in d]
cos_weights = [G_multi.edges[u, v]['Cosine'] for u, v in cos_edges]
nx.draw_networkx_edges(G_multi, pos, edgelist=cos_edges, edge_color=cos_weights, edge_cmap=CMAP_COS,
                       edge_vmin=threshold_cos, edge_vmax=cosine_similarity_df.values.max(),
                       width=EDGE_WIDTH, alpha=EDGE_ALPHA, style=(2, (2, 2)), ax=ax_main)

# Display labels only for core nodes.
if SHOW_NODE_LABELS:
    labels = {node: str(node) for node in G_multi.nodes() if node in G_core}
    nx.draw_networkx_labels(G_multi, pos, labels, font_size=NODE_LABEL_FONT_SIZE, ax=ax_main)

# Add colorbars for the edge weight colormaps.
sm_dtw = plt.cm.ScalarMappable(cmap=CMAP_DTW, norm=Normalize(vmin=threshold_sim_dtw, vmax=similarity_df_dtw.values.max()))
sm_dtw.set_array([])
fig.colorbar(sm_dtw, cax=ax_cbar_dtw, label='Soft-DTW Similarity')
sm_cos = plt.cm.ScalarMappable(cmap=CMAP_COS, norm=Normalize(vmin=threshold_cos, vmax=cosine_similarity_df.values.max()))
sm_cos.set_array([])
fig.colorbar(sm_cos, cax=ax_cbar_cos, label='Cosine Similarity')

# Create a legend for node colors.
custom_lines = [
    Line2D([0], [0], marker='o', color='w', label='Core Node', markerfacecolor=CORE_NODE_COLOR, markersize=10),
    Line2D([0], [0], marker='o', color='w', label='1-Hop Soft-DTW', markerfacecolor=ONE_HOP_DTW_COLOR, markersize=10),
    Line2D([0], [0], marker='o', color='w', label='1-Hop Cosine', markerfacecolor=ONE_HOP_COS_COLOR, markersize=10),
    Line2D([0], [0], marker='o', color='w', label='Both 1-Hop', markerfacecolor=BOTH_HOP_COLOR, markersize=10),
    Line2D([0], [0], marker='o', color='w', label='Other Nodes', markerfacecolor=OTHER_NODES_COLOR, markersize=10)
]
ax_legend.legend(handles=custom_lines, loc='upper center', frameon=False)

ax_main.axis('off')
plt.savefig(OUTPUT_PNG_NETWORK, dpi=300, bbox_inches='tight')
plt.show()
plt.close()
```

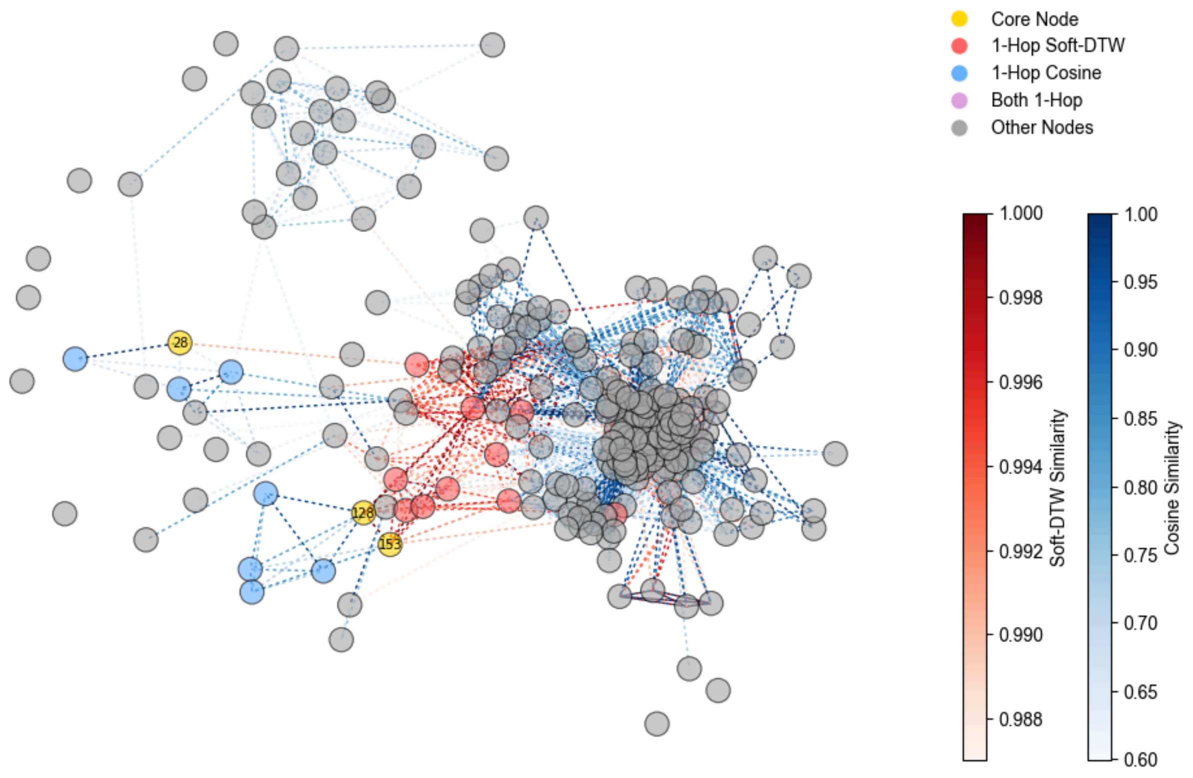


In [10]:

```
# ----------------------------------------
# 8. Subnetwork Visualization
# ----------------------------------------
# Visualize the subnetwork (candidate nodes with Size_Ratio >= 2) and display all node labels.
OUTPUT_PNG_SUBNETWORK = 'subnetwork.png'
G_sub = G_multi.subgraph(subnetwork_nodes).copy()
# Use the same layout as the main network.
pos_sub = {node: pos[node] for node in G_sub.nodes()}

node_colors_sub = []
for node in G_sub.nodes():
    if node in G_core:
        node_colors_sub.append(CORE_NODE_COLOR)
    elif node in n_hop_nodes_dtw and node in n_hop_nodes_cos:
        node_colors_sub.append(BOTH_HOP_COLOR)
    elif node in n_hop_nodes_dtw:
        node_colors_sub.append(ONE_HOP_DTW_COLOR)
    elif node in n_hop_nodes_cos:
        node_colors_sub.append(ONE_HOP_COS_COLOR)
    else:
        node_colors_sub.append(OTHER_NODES_COLOR)

fig_sub = plt.figure(figsize=(6, 4.5))
ax_main_sub = fig_sub.add_axes([0.05, 0.01, 0.60, 0.5])
# ax_cbar_sub_dtw = fig_sub.add_axes([0.68, 0.01, 0.015, 0.35])
# ax_cbar_sub_cos = fig_sub.add_axes([0.76, 0.01, 0.015, 0.35])
# ax_legend_sub = fig_sub.add_axes([0.6, 0.45, 0.25, 0.05])
# ax_legend_sub.axis('off')

nx.draw_networkx_nodes(G_sub, pos_sub, node_size=FIXED_NODE_SIZE, node_color=node_colors_sub,
                       edgecolors=FIXED_NODE_EDGE_COLOR, linewidths=FIXED_NODE_EDGE_WIDTH,
                       alpha=NODE_ALPHA, ax=ax_main_sub)

sub_dtw_edges = [(u, v) for u, v, d in G_sub.edges(data=True) if 'Soft_DTW' in d]
sub_dtw_weights = [G_sub.edges[u, v]['Soft_DTW'] for u, v in sub_dtw_edges]
nx.draw_networkx_edges(G_sub, pos_sub, edgelist=sub_dtw_edges, edge_color=sub_dtw_weights, edge_cmap=CMAP_DTW,
                       edge_vmin=threshold_sim_dtw, edge_vmax=similarity_df_dtw.values.max(),
                       width=EDGE_WIDTH, alpha=EDGE_ALPHA, style=(0, (1, 2)), ax=ax_main_sub)

sub_cos_edges = [(u, v) for u, v, d in G_sub.edges(data=True) if 'Cosine' in d]
sub_cos_weights = [G_sub.edges[u, v]['Cosine'] for u, v in sub_cos_edges]
nx.draw_networkx_edges(G_sub, pos_sub, edgelist=sub_cos_edges, edge_color=sub_cos_weights, edge_cmap=CMAP_COS,
                       edge_vmin=threshold_cos, edge_vmax=cosine_similarity_df.values.max(),
                       width=EDGE_WIDTH, alpha=EDGE_ALPHA, style=(2, (2, 2)), ax=ax_main_sub)

# Display labels for all nodes in the subnetwork.
if SHOW_NODE_LABELS:
    sub_labels = {node: str(node) for node in G_sub.nodes()}
    nx.draw_networkx_labels(G_sub, pos_sub, sub_labels, font_size=NODE_LABEL_FONT_SIZE, ax=ax_main_sub)

# sm_sub_dtw = plt.cm.ScalarMappable(cmap=CMAP_DTW, norm=Normalize(vmin=threshold_sim_dtw, vmax=similarity_df_dtw.values.max()))
# sm_sub_dtw.set_array([])
# fig_sub.colorbar(sm_sub_dtw, cax=ax_cbar_sub_dtw, label='Soft-DTW Similarity')
# sm_sub_cos = plt.cm.ScalarMappable(cmap=CMAP_COS, norm=Normalize(vmin=threshold_cos, vmax=cosine_similarity_df.values.max()))
# sm_sub_cos.set_array([])
# fig_sub.colorbar(sm_sub_cos, cax=ax_cbar_sub_cos, label='Cosine Similarity')

# custom_lines_sub = [
#     Line2D([0], [0], marker='o', color='w', label='Core Node', markerfacecolor=CORE_NODE_COLOR, markersize=10),
#     Line2D([0], [0], marker='o', color='w', label='1-Hop Soft-DTW', markerfacecolor=ONE_HOP_DTW_COLOR, markersize=10),
#     Line2D([0], [0], marker='o', color='w', label='1-Hop Cosine', markerfacecolor=ONE_HOP_COS_COLOR, markersize=10),
#     Line2D([0], [0], marker='o', color='w', label='Both 1-Hop', markerfacecolor=BOTH_HOP_COLOR, markersize=10),
#     Line2D([0], [0], marker='o', color='w', label='Other Nodes', markerfacecolor=OTHER_NODES_COLOR, markersize=10)
```

```python
# ]
# ax_legend_sub.legend(handles=custom_lines_sub, loc='upper center', frameon=False)

ax_main_sub.axis('off')
OUTPUT_PNG_SUBNETWORK = 'subnetwork.png'
plt.savefig(OUTPUT_PNG_SUBNETWORK, dpi=300, bbox_inches='tight')
plt.show()
plt.close()
```

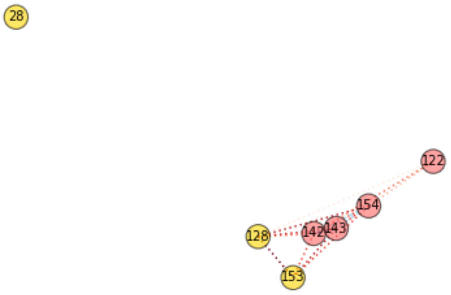
