## Supplementary material for "Time-resolved volatile organic compound profiling enables non-invasive detection of phenological progression in soybean": Supplemetal Materials: Supplementary Data S5.pdf

### Supplementary Data S5 The script and outputs for classifying soybean developmental stages using network analysis and community detection.

This Python script implements a reproducible workflow to summarize high-dimensional features by PCA, compare stage-wise centroids via cosine similarity, and construct a community-optimized network using ensemble Louvain with Optuna-based hyperparameter search. The workflow comprises: (1) loading `Train and Validation_2.csv`, indexing samples, selecting `AID_` features, and excluding baseline rows (`TimeID` = 99); (2) L1-normalizing each sample followed by feature standardization, performing PCA, plotting the cumulative explained variance (target ≥ 95%) and a PC1–PC2 projection (`cumulative_explained_variance_ratio.png`, `pca_projection.png`); (3) creating a pairwise PCA plot using the top components required to reach 95% variance (`pairwise_pca.png`); (4) computing stage-wise centroids in the selected PCA space, evaluating centroid-to-centroid cosine similarity, and visualizing a heatmap (`cosine_similarity_heatmap.png`); (5) building a centroid similarity network with edge weights from normalized cosine similarity; optimizing `edge_threshold`, `resolution`, and `matching_threshold` via Optuna (200 trials) using ensemble Louvain modularity as the objective; saving optimization history and plot (`optuna_optimization_history.csv`, `optimization_history_cosine.png`); (6) drawing the best network with community labels (`network_graph_cosine.png`) and exporting node/edge tables (`network_graph_nodes.csv`, `network_graph_edges.csv`); and (7) computing overall and per-community metrics—average clustering coefficient, modularity contribution, and conductance—exported as `network_metrics.csv` and summarized in `community_metrics_grid.png`.

In [1]:

```
import os
import pandas as pd
import numpy as np
import warnings
warnings.filterwarnings("ignore")
from tqdm import TqdmWarning
warnings.filterwarnings("ignore", category=TqdmWarning)
from sklearn.preprocessing import Normalizer, StandardScaler
from sklearn.decomposition import PCA
import matplotlib.pyplot as plt
import matplotlib.colors as mcolors
from matplotlib.cm import ScalarMappable
from matplotlib.lines import Line2D
from matplotlib.colors import Normalize
import seaborn as sns
import math
from sklearn.metrics.pairwise import cosine_similarity
import networkx as nx
from elouvain.ensemble_louvain import detect
import optuna
from collections import defaultdict
import logging

plt.rcParams["font.family"] = "Arial"
optuna.logging.set_verbosity(optuna.logging.WARNING)
```

In [2]:

```
# -------------------------------
# 1. Create output directories
# -------------------------------
output_dir = 'results'
plots_dir = os.path.join(output_dir, 'plots')
csv_dir = os.path.join(output_dir, 'csv')
os.makedirs(plots_dir, exist_ok=True)
os.makedirs(csv_dir, exist_ok=True)
```

In [3]:

```
# -------------------------------
# 2. Data Loading and Preprocessing
# -------------------------------
# Load data from CSV file.
original_df = pd.read_csv('Train and Validation_2.csv')

# Data pPreprocessing
original_df = original_df.reset_index().rename(columns={'index': 'instance_id'})
a_id_cols = [col for col in original_df.columns if col.startswith('AID_')]
df = original_df[['instance_id', 'TimeID'] + a_id_cols]
preprocessed_df = df[df['TimeID'] != 99].reset_index(drop=True)

# Separate features and labels.
X = preprocessed_df.drop(['instance_id', 'TimeID'], axis=1)
y = preprocessed_df['TimeID'].values

# Normalize each record using L1 norm, then standardize features (mean=0, std=1).
X_normalized = Normalizer(norm='l1').fit_transform(X)
X_standardized = StandardScaler().fit_transform(X_normalized)
```

In [4]:

```
# -------------------------------
# 3. PCA and Cumulative Explained Variance Plot
# -------------------------------
# Perform PCA on the standardized data.
pca = PCA()
X_pca_full = pca.fit_transform(X_standardized)

# Plot cumulative explained variance.
cumulative_variance = np.cumsum(pca.explained_variance_ratio_)
plt.figure(figsize=(4.5, 4))
plt.plot(range(1, len(cumulative_variance) + 1), cumulative_variance,
         marker='o', linestyle='--', label='Cumulative Variance')
plt.axhline(y=0.95, color='red', linestyle='--', label='95% Threshold')
plt.xlabel('Number of Principal Components', fontsize=14)
plt.ylabel('Cumulative Explained Variance', fontsize=14)
plt.xticks(fontsize=12)
plt.yticks(fontsize=12)
plt.grid(False)
plt.legend(fontsize=12)
plt.tight_layout()
plt.savefig(os.path.join(plots_dir, 'cumulative_explained_variance_ratio.png'), dpi=300)
plt.show()
plt.close()
```

```python
# Calculate the number of principal components needed to exceed 95% cumulative variance.
n_components_needed = np.argmax(cumulative_variance >= 0.95) + 1
print(f"Number of principal components required to reach 95% cumulative variance: {n_components_needed}")
```

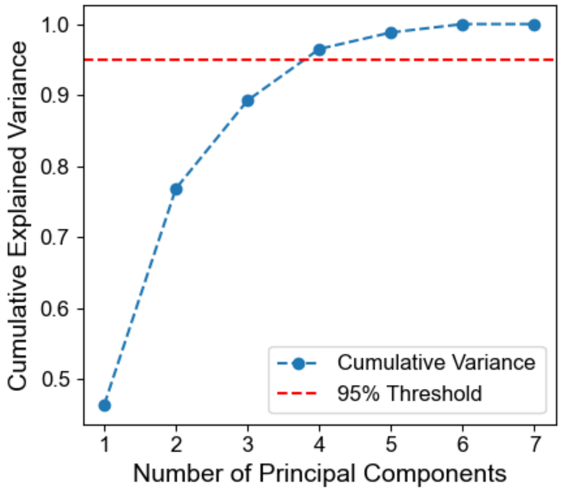


```
Number of principal components required to reach 95% cumulative variance: 4
```

In [5]:
```python
# ------------------------------
# 4. PCA Projection (First 2 Components)
# ------------------------------
# Extract the first two principal components for plotting.
X_pca = X_pca_full[:, :2]
# Calculate explained variance percentages for PC1 and PC2.
pc1_ratio = pca.explained_variance_ratio_[0] * 100
pc2_ratio = pca.explained_variance_ratio_[1] * 100
# Create a DataFrame with PCA scores and include the TimeID column.
pca_df = pd.DataFrame(X_pca, columns=[f'PC1 ({pc1_ratio:.1f}%)', f'PC2 ({pc2_ratio:.1f}%)'])
pca_df['TimeID'] = y

# Plot PCA projection scatter plot.
plt.figure(figsize=(5, 4))
scatter = plt.scatter(pca_df.iloc[:, 0], pca_df.iloc[:, 1], c=pca_df['TimeID'],
                      cmap='viridis', alpha=0.7)
plt.xlabel(f'PC1 ({pc1_ratio:.1f}%)', fontsize=12)
plt.ylabel(f'PC2 ({pc2_ratio:.1f}%)', fontsize=12)
cbar = plt.colorbar(scatter)
cbar.set_label('DAS', fontsize=12)
plt.xticks(fontsize=10)
plt.yticks(fontsize=10)
plt.grid(False)
plt.tight_layout()
plt.savefig(os.path.join(plots_dir, 'pca_projection.png'), dpi=300)
plt.show()
plt.close()
```

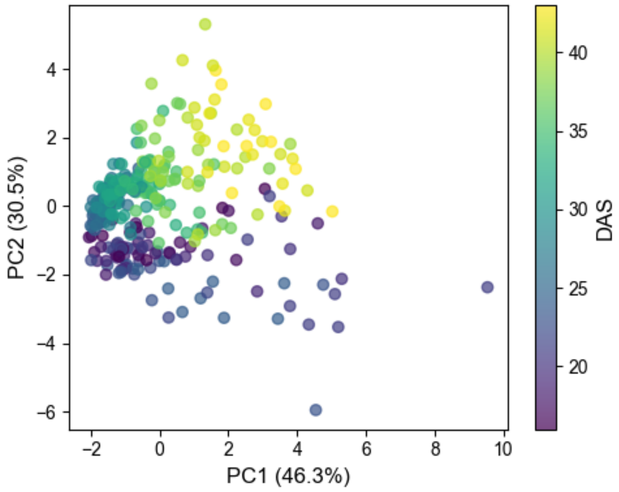


In [6]:
```python
# ------------------------------
# 5. Pairwise Plot
# ------------------------------
# Create a pairplot.
pca_n_components = n_components_needed
pca_columns = [f'PC{i+1}' for i in range(pca_n_components)]
pca_df_full = pd.DataFrame(X_pca_full[:, :pca_n_components], columns=pca_columns)
pca_df_full['TimeID'] = y

# Use seaborn to create the pairplot with KDE on the diagonal.
g = sns.pairplot(pca_df_full, vars=pca_columns, hue='TimeID', palette='viridis',
                 diag_kind='kde', height=1.2, aspect=1.1, plot_kws={"s": 8})
for ax in g.axes.flatten():
    ax.grid(False)
if g._legend is not None:
    g._legend.remove()

# Set up the colorbar.
norm = mcolors.Normalize(pca_df_full['TimeID'].min(), pca_df_full['TimeID'].max())
sm = ScalarMappable(cmap='viridis', norm=norm)
```

```
sm.set_array([])

g.fig.subplots_adjust(right=0.85)
cbar_ax = g.fig.add_axes([0.88, 0.15, 0.03, 0.7])
g.fig.colorbar(sm, cax=cbar_ax, label='DAS')

plt.savefig(os.path.join(plots_dir, 'pairwise_pca.png'), dpi=300)
plt.show()
plt.close()
```

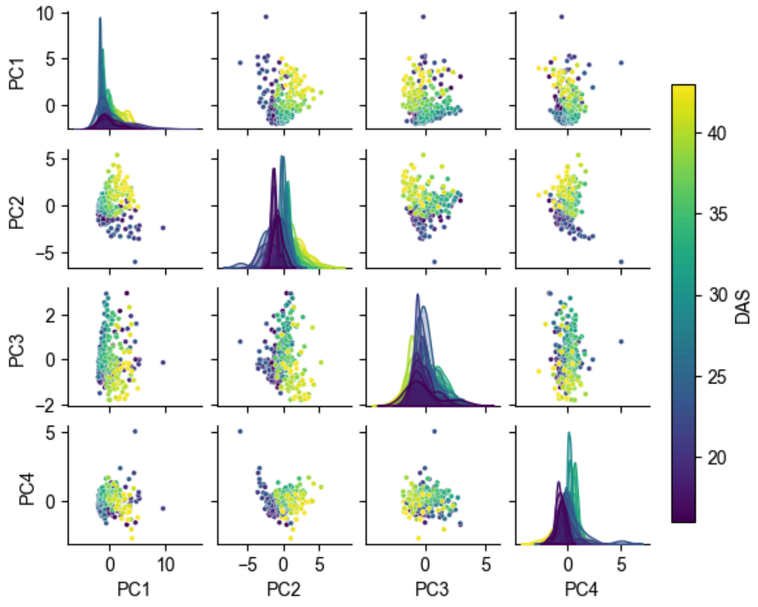


In [7]:

```
# -------------------------------
# 6. Cosine Similarity Heatmap of TimeID Centroids
# -------------------------------
# Calculate centroids using only the principal components required to reach 95% cumulative variance.
X_pca_selected = X_pca_full[:, :n_components_needed]
pca_columns_selected = [f'PC{i+1}' for i in range(n_components_needed)]
pca_df_selected = pd.DataFrame(X_pca_selected, columns=pca_columns_selected)
pca_df_selected['TimeID'] = y

# Compute the centroid for each TimeID by taking the mean.
centroids = pca_df_selected.groupby('TimeID')[pca_columns_selected].mean()

# Compute cosine similarity among centroids.
cosine_sim_matrix = cosine_similarity(centroids.values)
cosine_sim_df = pd.DataFrame(cosine_sim_matrix, index=centroids.index, columns=centroids.index)

# Plot the cosine similarity heatmap.
fig, ax = plt.subplots(figsize=(6, 4.8))
sns.heatmap(
    cosine_sim_df,
    cmap='Blues',
    annot=False,
    square=True,
    linewidths=0.5,
    cbar=False,
    ax=ax
)

# Add a horizontal colorbar below the heatmap, half width.
cax = fig.add_axes([0.3, 0.001, 0.4, 0.015])
cb = fig.colorbar(
    ax.collections[0],
    cax=cax,
    orientation='horizontal'
)
cb.set_label('Cosine Similarity', fontsize=12)
cb.ax.tick_params(labelsize=9)
cb.outline.set_visible(False)

# Center tick marks in each cell
n = cosine_sim_df.shape[1]
ax.set_xticks(np.arange(n) + 0.5)
ax.set_yticks(np.arange(n) + 0.5)
ax.set_xticklabels(cosine_sim_df.columns, rotation=45, ha='center', fontsize=9)
ax.set_yticklabels(cosine_sim_df.index,   rotation=0,  va='center', fontsize=9)
ax.set_xlim(0, n)
ax.set_ylim(n, 0)

ax.set_xlabel('DAS', fontsize=14)
ax.set_ylabel('DAS', fontsize=14)

plt.tight_layout()
plt.savefig(
    os.path.join(plots_dir, 'cosine_similarity_heatmap.png'),
    dpi=300,
    bbox_inches='tight',
    pad_inches=0.02
)
plt.show()
plt.close()
```

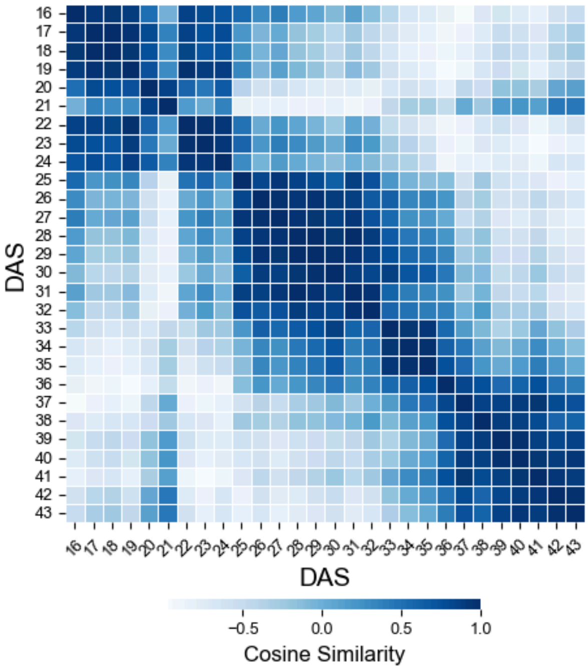


In [8]:
```python
# ------------------------------
# 7. Define Network Graph Functions
# ------------------------------
def create_network_graph(matrix_df, edge_threshold=0.8, resolution=1.0, matching_threshold=0.99, n_runs=100):
    """
    Create a network graph based on cosine similarity.

    Parameters:
        matrix_df (DataFrame): A DataFrame containing the cosine similarity matrix (correlations between centroids).
        edge_threshold (float): Only add an edge if the normalized weight is above this threshold.
        resolution (float): Resolution parameter used in community detection.
        matching_threshold (float): Matching threshold for community detection.
        n_runs (int): Number of runs for community detection.

    Returns:
        tuple: (modularity, G, partition)
            modularity: The modularity score of the graph.
            G: The created network graph (networkx.Graph).
            partition: The community partitioning result (as a dictionary) or None.
    """
    # Create a graph object and add nodes based on centroid TimeIDs.
    G = nx.Graph()
    G.add_nodes_from(centroids.index)

    # Get the minimum and maximum values of the cosine similarity matrix for normalization.
    values = matrix_df.values
    max_value = values.max()
    min_value = values.min()
    denom = max(max_value - min_value, 1e-12)

    # Add an edge between nodes if the normalized weight exceeds the edge_threshold.
    for i, time_id1 in enumerate(centroids.index):
        for j, time_id2 in enumerate(centroids.index):
            if i < j:
                value = matrix_df.loc[time_id1, time_id2]
                weight = (value - min_value) / (max_value - min_value)
                if weight >= edge_threshold:
                    G.add_edge(time_id1, time_id2, weight=weight, value=value)

    # If there are edges, perform community detection and calculate modularity.
    if G.number_of_edges() == 0:
        return 0.0, G, None

    # ensemble Louvain
    partition = None
    try:
        partition = detect(G, partitions=n_runs, matching_threshold=matching_threshold, resolution=resolution)
    except Exception as e:
        print(f"Error during community detection: {e}")
        return 0.0, G, None

    all_nodes = set(G.nodes())
    assigned = set(partition.keys()) if partition is not None else set()
    if assigned != all_nodes:
        next_id = (max(partition.values()) if partition else -1) + 1
        for u in (all_nodes - assigned):
            partition[u] = next_id
            next_id += 1

    community_dict = defaultdict(list)
    for node, comm in partition.items():
        community_dict[comm].append(node)
    communities = list(community_dict.values())
    modularity = nx.algorithms.community.modularity(G, communities, weight='weight')

    return modularity, G, partition

def draw_network_graph(
    G_final,
```

```
    partition,
    edge_threshold,
    apply_community_detection=True,
    edge_threshold_type='absolute',
    edge_threshold_value=None,
    save_path=None,
    start_angle=np.pi/2,
    end_angle=-np.pi/2,
    community_scale=3.0,
    node_radius=1.0,
    community_order=None,
    label_fontweight='800'
):
    """
    Draw the network graph.

    Parameters:
        G_final (networkx.Graph): The network graph to be drawn.
        partition (dict or None): The community partitioning result. If None, treat all nodes as one group.
        edge_threshold (float): The edge threshold (displayed in the title).
        apply_community_detection (bool): Whether to reflect the community detection results.
        edge_threshold_type (str): The type of threshold (currently fixed as 'absolute').
        edge_threshold_value (float): The value of the threshold (displayed in the title).
        save_path (str or None): Path to save the figure. If None, the figure is not saved.
        start_angle (float): The starting angle for community layout.
        end_angle (float): The ending angle for community layout.
        community_scale (float): Scale factor for spacing between communities.
        node_radius (float): Radius for spacing nodes within each community.
        community_order (list or None): The order of communities for display. If None, sort the communities.
        label_fontweight (str): The font weight for node labels.

    Returns:
        None
    """
    # If no partition information is provided, treat all nodes as one community.
    if partition is None:
        community_labels = {node: 0 for node in G_final.nodes()}
    else:
        community_labels = {node: partition[node] for node in G_final.nodes()}

    # Create a dictionary grouping nodes by their community labels.
    community_dict = defaultdict(list)
    for node, community in community_labels.items():
        community_dict[community].append(node)

    # Sort communities if no specific order is provided.
    if community_order is None:
        sorted_communities = sorted(community_dict.keys())
    else:
        sorted_communities = [comm for comm in community_order if comm in community_dict]

    num_communities = len(sorted_communities)
    reds = plt.get_cmap('viridis', num_communities)
    color_map = {comm: reds(i) for i, comm in enumerate(sorted_communities)}

    # Calculate angles for arranging communities from start_angle to end_angle.
    community_centers = {}
    if num_communities > 1:
        step = (start_angle - end_angle) / (num_communities - 1)
        for i, comm in enumerate(sorted_communities):
            angle = start_angle - i * step
            community_centers[comm] = (
                np.cos(angle) * community_scale,
                np.sin(angle) * community_scale
            )
    else:
        community_centers[sorted_communities[0]] = (
            np.cos(start_angle) * community_scale,
            np.sin(start_angle) * community_scale
        )

    # Determine positions for each node based on its community center.
    pos = {}
    for comm, nodes in community_dict.items():
        center_x, center_y = community_centers[comm]
        nodes_sorted = sorted(nodes)
        community_pos = {}
        if len(nodes_sorted) > 1:
            n = len(nodes_sorted)
            angles = start_angle - np.arange(n) * (2 * np.pi / n)
            for node, ang in zip(nodes_sorted, angles):
                community_pos[node] = (
                    np.cos(ang) * node_radius,
                    np.sin(ang) * node_radius
                )
        else:
            community_pos[nodes_sorted[0]] = (0, 0)
        for node, (x, y) in community_pos.items():
            pos[node] = (x + center_x, y + center_y)

    # Set up the figure and axes for drawing the graph.
    fig = plt.figure(figsize=(16, 12))
    ax_main = fig.add_axes([0, 0.05, 0.5, 0.9])
    ax_cbar = fig.add_axes([0.50, 0.20, 0.02, 0.4])
    ax_comm_legend = fig.add_axes([0.50, 0.55, 0.05, 0.3])
    ax_comm_legend.axis('off')

    # Draw edges with widths proportional to their weights.
    edges = G_final.edges(data=True)
    weights = [data['weight'] for _, _, data in edges]
    if weights:
```

```python
        edge_cmap = plt.get_cmap('Blues')
        edge_norm = mcolors.Normalize(vmin=min(weights), vmax=max(weights))
        sm = ScalarMappable(cmap=edge_cmap, norm=edge_norm)
        sm.set_array([])
        nx.draw_networkx_edges(G_final, pos, edgelist=edges, width=[w * 1.5 for w in weights],
                               edge_color=weights, edge_cmap=edge_cmap, edge_vmin=min(weights),
                               edge_vmax=max(weights), alpha=0.7, ax=ax_main)
    else:
        nx.draw_networkx_edges(G_final, pos, alpha=0.7, ax=ax_main)

    # Draw nodes with colors based on their community.
    node_colors = [color_map[community_labels[node]] for node in G_final.nodes()]
    nx.draw_networkx_nodes(G_final, pos, node_color=node_colors, node_size=600,
                           edgecolors='k', linewidths=1, alpha=0.9, ax=ax_main)

    # Add labels to nodes.
    labels = {node: str(node) for node in G_final.nodes()}
    for node, label in labels.items():
        x, y = pos[node]
        r, g, b, _ = color_map[community_labels[node]]
        brightness = 0.299 * r + 0.587 * g + 0.114 * b
        label_color = 'white' if brightness < 0.5 else 'black'
        ax_main.text(x, y, label, fontsize=15, color=label_color,
                     ha='center', va='center', fontweight=label_fontweight)

    # Create legend entries for communities.
    community_handles = [Line2D([0], [0], marker='o', color='w', label=f'{comm+1}',
                                markerfacecolor=color_map[comm], markersize=14, markeredgecolor='k')
                         for comm in sorted_communities]
    ax_comm_legend.legend(handles=community_handles, title='Community ID',
                          loc='center', ncol=1, frameon=False, fontsize=15, title_fontsize=18)

    # Add a colorbar for edge weights
    if weights:
        cbar = fig.colorbar(sm, cax=ax_cbar, orientation='vertical',
                            label='Edge Weight (Normalized Cosine Similarity)')
        cbar.ax.tick_params(labelsize=15)
        cbar.ax.yaxis.label.set_size(18)
        cbar.ax.yaxis.set_label_coords(3, 0.5)

    ax_main.set_aspect('equal')
    ax_main.axis('off')
    ax_comm_legend.axis('off')

    # Save the figure
    if save_path:
        plt.savefig(save_path, dpi=300, bbox_inches='tight')
    plt.show()
    plt.close()

    # Save node positions and community labels
    node_info = []
    for node in G_final.nodes():
        x, y = pos[node]
        node_info.append({'node': node, 'x': x, 'y': y, 'community': community_labels[node] + 1})
    node_info_df = pd.DataFrame(node_info)
    node_info_csv_path = os.path.join(csv_dir, 'network_graph_nodes.csv')
    node_info_df.to_csv(node_info_csv_path, index=False)

    # Save edge information
    edge_info = []
    for u, v, data in G_final.edges(data=True):
        edge_info.append({'u': u, 'v': v, 'weight': data['weight'], 'value': data.get('value', None)})
    edge_info_df = pd.DataFrame(edge_info)
    edge_info_csv_path = os.path.join(csv_dir, 'network_graph_edges.csv')
    edge_info_df.to_csv(edge_info_csv_path, index=False)
```

In [9]:
```python
# -------------------------------
# 8. Optimize Cosine Similarity-based Network
# -------------------------------
def objective(trial, matrix_df):
    """
    Objective function for Optuna optimization.

    Parameters:
        trial: An Optuna Trial object.
        matrix_df (DataFrame): A DataFrame containing the cosine similarity matrix (correlations between centroids).

    Returns:
        float: The modularity value to be optimized.
    """
    # Suggest parameter values within specified ranges.
    edge_threshold = trial.suggest_float('edge_threshold', 0.5, 0.9)
    resolution = trial.suggest_float('resolution', 0.01, 10)
    matching_threshold = trial.suggest_float('matching_threshold', 0.6, 1)
    modularity, G_tmp, _ = create_network_graph(
        matrix_df=matrix_df,
        edge_threshold=edge_threshold,
        resolution=resolution,
        matching_threshold=matching_threshold
    )

    if G_tmp.number_of_edges() == 0:
        return -1e6

    return modularity

# Create an Optuna Study object and optimize the objective function.
study_cosine = optuna.create_study(direction='maximize', sampler=optuna.samplers.TPESampler(),
                                   pruner=optuna.pruners.MedianPruner())
study_cosine.optimize(lambda trial: objective(trial, cosine_sim_df),
```

```python
                          n_trials=200, show_progress_bar=True)

# Get the best parameters from the study.
best_params_cosine = study_cosine.best_params
print("Best Parameters:", best_params_cosine)

# Plot the optimization history and save the plot and CSV.
def plot_custom_optimization_history(study, best_params, save_path):
    """
    Plot the optimization history (modularity over iterations) and save the plot.
    Additionally, save the optimization history as CSV.

    Parameters:
        study: An Optuna Study object.
        best_params (dict): Dictionary of the best parameters.
        title (str): Title of the plot.
        save_path (str): File path to save the plot image.

    Returns:
        None
    """
    # Retrieve modularity values for each trial.
    modularities = [trial.value for trial in study.trials]
    iterations = list(range(1, len(modularities) + 1))
    best_trial = study.best_trial
    best_iteration = best_trial.number + 1
    best_modularity = best_trial.value
    best_params = best_trial.params

    # Plot optimization history.
    plt.figure(figsize=(8, 4))
    plt.plot(iterations, modularities, marker='o', linestyle='-', color='blue', label='Modularity')
    plt.scatter(best_iteration, best_modularity, color='red', zorder=5, label='Best Modularity')
    plt.axhline(y=best_modularity, color='red', linestyle='--', linewidth=1)
    plt.axvline(x=best_iteration, color='red', linestyle='--', linewidth=1)
    ymin, ymax = plt.ylim()
    y_offset = (ymax - ymin) * 0.9
    annotation_text = (
        f"Iteration: {best_iteration}\n"
        f"Modularity: {best_modularity:.4f}\n"
        f"Edge Threshold: {best_params['edge_threshold']:.4f}\n"
        f"Resolution: {best_params['resolution']:.4f}\n"
        f"Matching Threshold: {best_params['matching_threshold']:.4f}"
    )
    plt.annotate(
        annotation_text,
        xy=(best_iteration, best_modularity),
        xytext=(best_iteration + 5, best_modularity - y_offset),
        color='red',
        fontsize=10,
        bbox=dict(boxstyle="round,pad=0.3", edgecolor='red', facecolor='white', alpha=0.8),
        arrowprops=dict(arrowstyle="->", connectionstyle="arc3,rad=0", color='red')
    )
    plt.xlabel('Iteration', fontsize=14)
    plt.ylabel('Modularity', fontsize=14)
    plt.legend(fontsize=12,
               loc='center left',
               bbox_to_anchor=(1.02, 0.5),
               borderaxespad=0,
               frameon=False
    )
    plt.xticks(fontsize=12)
    plt.yticks(fontsize=12)
    plt.grid(False)
    plt.tight_layout(rect=[0, 0, 0.85, 1])
    plt.savefig(save_path, dpi=300, bbox_inches='tight')
    plt.show()
    plt.close()

    # Organize the optimization history into a DataFrame and output as CSV.
    history_records = []
    for trial in study.trials:
        record = {
            "Trial Number": trial.number,
            "Modularity": trial.value,
            "Edge Threshold": trial.params.get("edge_threshold"),
            "Resolution": trial.params.get("resolution"),
            "Matching Threshold": trial.params.get("matching_threshold")
        }
        history_records.append(record)
    history_df = pd.DataFrame(history_records)
    history_csv_path = os.path.join(csv_dir, "optuna_optimization_history.csv")
    history_df.to_csv(history_csv_path, index=False)

plot_custom_optimization_history(
    study=study_cosine,
    best_params=best_params_cosine,
    save_path=os.path.join(plots_dir, 'optimization_history_cosine.png')
)
```

```
Best trial: 24. Best value: 0.683718: 100%|██████████████████████████████████████████| 200/200 [00:15<00:00, 12.83it/s]
Best Parameters: {'edge_threshold': 0.8990040776133512, 'resolution': 0.9083246220350855, 'matching_threshold': 0.6024052706791009}
```

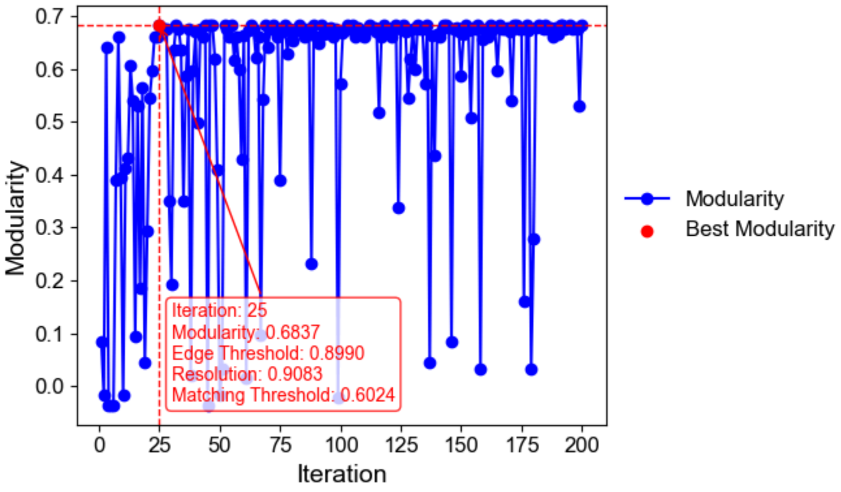


In [10]:

```
# -------------------------------
# 9. Draw Network Graph based on Cosine Similarity
# -------------------------------
# Create and draw the network graph using the best parameters from the optimization.
modularity_cosine, best_G_cosine, best_partition_cosine = create_network_graph(
    cosine_sim_df, edge_threshold=best_params_cosine['edge_threshold'],
    resolution=best_params_cosine['resolution'], matching_threshold=best_params_cosine['matching_threshold']
)
draw_network_graph(
    G_final=best_G_cosine,
    partition=best_partition_cosine,
    edge_threshold=best_params_cosine['edge_threshold'],
    apply_community_detection=True,
    edge_threshold_type='absolute',
    edge_threshold_value=best_params_cosine['edge_threshold'],
    save_path=os.path.join(plots_dir, 'network_graph_cosine.png'),
    start_angle=np.pi/3,
    end_angle=-np.pi/3,
    community_scale=2.2,
    node_radius=0.4,
    community_order=None,
    label_fontweight='800'
)
```

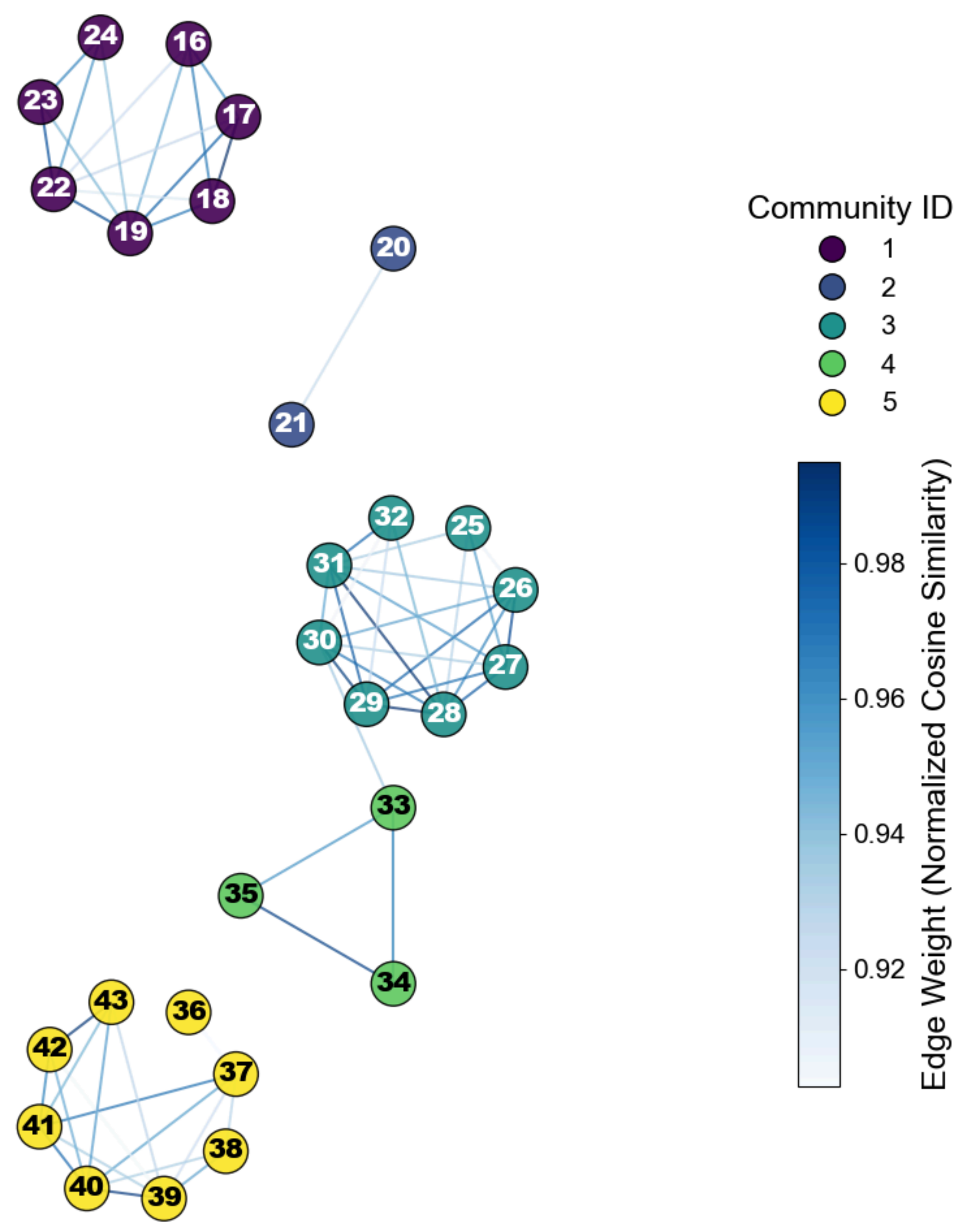


In [11]:
```
# ------------------------------
# 10. Analyze Network Metrics and Save to CSV
# ------------------------------
# Utility
def darken_color(color, factor=0.7):
    """
    Darken an RGBA color by multiplying each RGB channel.

    Parameters
    ----------
    color : tuple
        RGBA tuple (R, G, B, A) in the range [0, 1].
    factor : float, optional
        Multiplicative factor (< 1) applied to R, G, B. Default is 0.7.

    Returns
    -------
    tuple
        Darkened RGBA tuple.
    """
    r, g, b, a = color
    return (r * factor, g * factor, b * factor, a)



# Metric helper functions
def _modularity_contribution(G, nodes, total_w):
    """
    Compute the weighted modularity contribution of a community so that
    the sum over all communities matches NetworkX ``modularity``.

    Formula
    -------
    Q_c = (l_c / m) - (d_c / 2m)²
```

```python
      * l_c : total internal edge weight of the community
      * d_c : sum of weighted degrees of nodes in the community
      * m   : total edge weight of the whole graph (``total_w``)

    Parameters
    ----------
    G : networkx.Graph
        Target graph.
    nodes : Iterable
        Nodes belonging to the community.
    total_w : float
        Total edge weight ``m`` of ``G``.

    Returns
    -------
    float
        Modularity contribution of the specified community.
    """
    S = set(nodes)
    l_c = G.subgraph(S).size(weight="weight")                # internal weight
    d_c = sum(dict(G.degree(S, weight="weight")).values())   # total degree
    e_c = l_c / total_w                                      # l_c / m
    a_c = d_c / (2.0 * total_w)                              # d_c / 2m
    return e_c - a_c**2


def _conductance(G, nodes):
    """
    Calculate weighted conductance ϕ(S) of a node set S.

    ϕ(S) = cut(S, ¬S) / vol(S)

    Parameters
    ----------
    G : networkx.Graph
        Target graph.
    nodes : Iterable
        Nodes in set S.

    Returns
    -------
    float
        Conductance value. ``nan`` if ``vol(S) == 0``.
    """
    S = set(nodes)
    cut_w, vol_S = 0.0, 0.0
    for u in S:
        for v, data in G[u].items():
            w = data.get("weight", 1.0)
            if v not in S:
                cut_w += w
        vol_S += G.degree(u, weight="weight")
    return np.nan if vol_S == 0 else cut_w / vol_S


def _compute_metrics(G, nodes, total_w):
    """
    Compute three community quality metrics.

    Parameters
    ----------
    G : networkx.Graph
        Target graph.
    nodes : Iterable
        Nodes composing the community.
    total_w : float
        Total edge weight ``m`` of ``G``.

    Returns
    -------
    dict
        Keys:
          * 'Average Clustering Coefficient'
          * 'Modularity Contribution'
          * 'Conductance'
    """
    clust_vals = [nx.clustering(G, weight="weight").get(n, np.nan) for n in nodes]
    return {
        "Average Clustering Coefficient": np.nanmean(clust_vals)
        if len(clust_vals) > 0
        else np.nan,
        "Modularity Contribution": _modularity_contribution(G, nodes, total_w),
        "Conductance": _conductance(G, nodes),
    }


# Main analysis routine
def analyze_network(G, partition, csv_dir):
    """
    Calculate network/community metrics, export CSV, and return as DataFrame.

    Parameters
    ----------
    G : networkx.Graph
        Full graph.
    partition : dict or None
        Mapping ``{node: community_id}``. If ``None``, only overall metrics
        are computed.
    csv_dir : str
        Directory path to save 'network_metrics.csv'.
```

```python
    Returns
    -------
    pandas.DataFrame
        Metric table including overall and per-community rows.
    """
    total_w = G.size(weight="weight")  # m
    rows = []

    # ---- overall network ---------------------------------
    overall = _compute_metrics(G, G.nodes(), total_w)
    overall["Type"] = "Overall Network"
    overall["Community ID"] = np.nan
    rows.append(overall)

    # ---- communities -------------------------------------
    if partition is not None:
        comms = defaultdict(list)
        for n, c in partition.items():
            comms[c].append(n)

        # overall modularity (printed & stored)
        overall_mod = nx.algorithms.community.modularity(
            G, comms.values(), weight="weight"
        )
        overall["Weighted Modularity"] = overall_mod
        print(f"Overall weighted modularity: {overall_mod:.4f}")

        for cid, nodes in comms.items():
            row = _compute_metrics(G, nodes, total_w)
            row["Type"] = "Community"
            row["Community ID"] = cid + 1  # 1-indexed
            row["Weighted Modularity"] = np.nan
            rows.append(row)
    else:
        overall["Weighted Modularity"] = np.nan

    df = pd.DataFrame(rows)
    df.to_csv(os.path.join(csv_dir, "network_metrics.csv"), index=False)
    return df


# Grid visualization
def plot_metrics_grid(G, partition, metrics_df, plots_dir, weighted=True):
    """
    Draw a 1×3 grid comparing:
      1. Average clustering coefficient (bar + dots)
      2. Modularity contribution (bar)
      3. Conductance (bar)

    All visual styles (colors, axes, bar outlines) replicate the design used
    for clustering-coefficient plots elsewhere in the workflow.

    Parameters
    ----------
    G : networkx.Graph
        Full graph.
    partition : dict
        Community assignment ``{node: community_id}``.
    metrics_df : pandas.DataFrame
        Output of ``analyze_network()``.
    plots_dir : str
        Directory to save 'community_metrics_grid.png'.
    weighted : bool, optional
        Use edge weights for clustering coefficient. Default ``True``.
    """
    # ----- common data & color map ------------------------
    comm_vals_clust = defaultdict(list)
    clust_dict = nx.clustering(G, weight="weight" if weighted else None)
    for n, c in partition.items():
        if n in clust_dict:
            comm_vals_clust[c].append(clust_dict[n])

    sorted_comms = sorted(comm_vals_clust.keys())
    comm_ids = [c + 1 for c in sorted_comms]
    cmap = plt.get_cmap("viridis", len(sorted_comms))
    color_map = {c: cmap(i) for i, c in enumerate(sorted_comms)}

    comm_df = (
        metrics_df[metrics_df["Type"] == "Community"]
        .set_index("Community ID")
        .loc[comm_ids]
    )
    mod_vals = comm_df["Modularity Contribution"].values
    cond_vals = comm_df["Conductance"].values

    # ----- create grid figure ----------------------------
    fig, axes = plt.subplots(1, 3, figsize=(15, 4), sharex=False)

    # 1) Clustering
    means, errs = [], []
    for c in sorted_comms:
        vals = np.array(comm_vals_clust[c])
        means.append(np.mean(vals))
        errs.append(np.std(vals) / np.sqrt(len(vals)) if len(vals) else 0)
    axes[0].bar(
        comm_ids, means, yerr=errs, capsize=5,
        color=[color_map[c] for c in sorted_comms],
        edgecolor="black", alpha=0.7,
    )
    # dot overlay
    for c in sorted_comms:
```

```
        x = c + 1
        vals = comm_vals_clust[c]
        jitter = np.random.uniform(-0.1, 0.1, size=len(vals))
        fill = color_map[c]
        edge = darken_color(fill, 0.6)
        axes[0].scatter(
            np.full(len(vals), x) + jitter, vals,
            facecolors=fill, edgecolors=edge, linewidths=1.5,
            zorder=10, alpha=0.9,
        )
    axes[0].set_ylabel("Clustering Coefficient", fontsize=12)
    axes[0].set_title("Average Clustering", fontsize=13)

    # 2) Modularity contribution
    axes[1].bar(
        comm_ids, mod_vals,
        color=[color_map[c] for c in sorted_comms],
        edgecolor="black", alpha=0.7,
    )
    axes[1].set_ylabel("Modularity Contribution", fontsize=12)
    axes[1].set_title("Modularity Contribution", fontsize=13)

    # 3) Conductance
    axes[2].bar(
        comm_ids, cond_vals,
        color=[color_map[c] for c in sorted_comms],
        edgecolor="black", alpha=0.7,
    )
    axes[2].set_ylabel("Conductance", fontsize=12)
    axes[2].set_title("Conductance", fontsize=13)

    # common x-axis settings
    for ax in axes:
        ax.set_xlabel("Community ID", fontsize=12)
        ax.set_xticks(comm_ids)
        ax.set_xticklabels([str(cid) for cid in comm_ids], fontsize=11)
        ax.axhline(y=0, color="darkgray", linewidth=1)

    plt.tight_layout()
    plt.savefig(os.path.join(plots_dir, "community_metrics_grid.png"), dpi=300)
    plt.show()
    plt.close()


# Execute analysis & plot
metrics_df = analyze_network(
    G=best_G_cosine,
    partition=best_partition_cosine,
    csv_dir=csv_dir
)

plot_metrics_grid(
    G=best_G_cosine,
    partition=best_partition_cosine,
    metrics_df=metrics_df,
    plots_dir=plots_dir,
    weighted=True
)
```

```
Overall weighted modularity: 0.6837
```

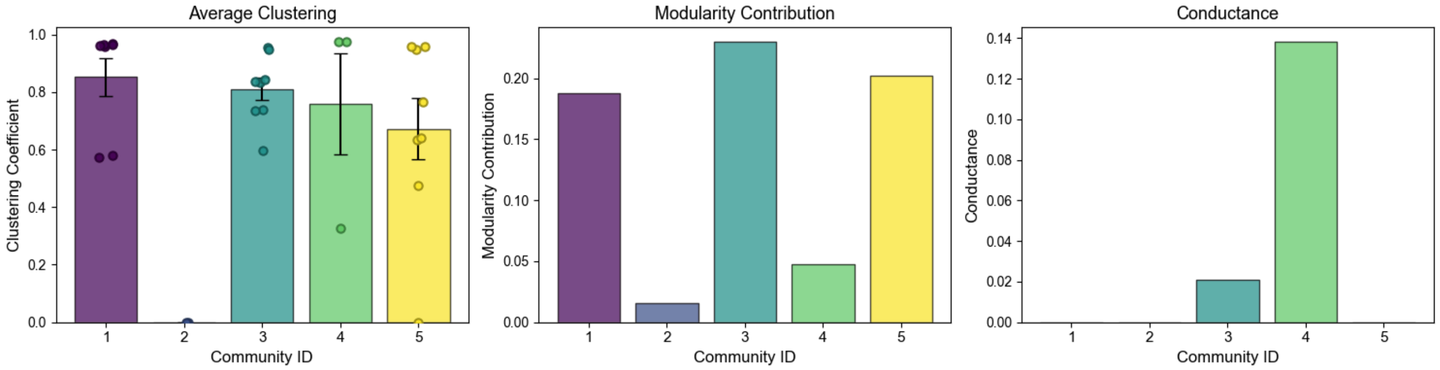
