## Supplementary material for "Time-resolved volatile organic compound profiling enables non-invasive detection of phenological progression in soybean": Supplemetal Materials: Supplementary Data S6.pdf

### Supplementary Data S6 The script and outputs for predicting soybean developmental stages from VOC profiles using TabPFN.

This Python script implements a reproducible classification workflow for imbalanced tabular data using KMeansSMOTE hyperparameter optimization and a post-hoc TabPFN ensemble. The pipeline comprises: (1) loading `Train and Validation_2_woBlank_wClass.csv`, label encoding, and a 75/25 train–validation split with stratification by `Subclass` (DAS); (2) tuning KMeansSMOTE (`k_neighbors`, `n_clusters`, `cluster_balance_threshold`) via Optuna (TPE; 300 trials) to maximize silhouette score on the resampled feature space, followed by a robust, multi-step fallback resampling routine; (3) standardization fitted on the resampled training set and applied to validation, full, and external ("unseen") sets; (4) fitting an `AutoTabPFNClassifier` (`presets="best_quality"`, `max_time=3000`, `eval_metric="roc_auc_ovo_macro"`, `balance_probabilities=True`) with CUDA; (5) evaluation on the 25% hold-out validation and an independent external set `Unseen_2_woBlank_wClass.csv`: classification reports, confusion matrices, and key metrics (accuracy, balanced accuracy, macro-precision/recall/F1, macro ROC-AUC, log loss); (6) ROC and PR curves per class plus macro-average on the external set; and (7) 95% stratified bootstrap confidence intervals (n = 1,000) for external-set metrics.

In [1]:
```
import os
os.environ.pop("SKLEARN_ARRAY_API", None)
os.environ["SCIPY_ARRAY_API"] = "1"
os.environ["LOKY_MAX_CPU_COUNT"] = "1"

import warnings
warnings.filterwarnings("ignore")
from tqdm import TqdmWarning
warnings.filterwarnings("ignore", category=TqdmWarning)

import json
import logging
from collections import defaultdict

import numpy as np
import pandas as pd
import matplotlib.pyplot as plt
import seaborn as sns

import optuna
from optuna.samplers import TPESampler
optuna.logging.set_verbosity(optuna.logging.WARNING)

from sklearn.preprocessing import LabelEncoder, StandardScaler, Normalizer, label_binarize
from sklearn.model_selection import train_test_split
from sklearn.metrics import (
    accuracy_score, balanced_accuracy_score, precision_recall_fscore_support,
    roc_auc_score, log_loss, classification_report, confusion_matrix,
    roc_curve, auc, precision_recall_curve, silhouette_score
)

from sklearn.cluster import KMeans
from imblearn.over_sampling import KMeansSMOTE
from tabpfn_extensions.post_hoc_ensembles.sklearn_interface import AutoTabPFNClassifier

plt.rcParams["font.family"] = "Arial"
```

In [2]:
```
# ------------------------------
# 1) Output directories
# ------------------------------
output_dir = "results"
plots_dir  = os.path.join(output_dir, "plots")
csv_dir    = os.path.join(output_dir, "csv")
os.makedirs(plots_dir, exist_ok=True)
os.makedirs(csv_dir, exist_ok=True)
```

In [3]:
```
# ------------------------------
# 2) Data loading & basic preprocessing
# ------------------------------
# Expect columns: "Class", optional "Subclass", and many "AID_*" feature columns
df = pd.read_csv("Train and Validation_2_woBlank_wClass.csv").dropna(subset=["Class"])

# Label encoding for target
le = LabelEncoder()
df["Class_encoded"] = le.fit_transform(df["Class"])
class_names = le.classes_.astype(str).tolist()
n_classes   = len(class_names)

# Feature matrix
feature_cols = [c for c in df.columns if c.startswith("AID_")]
X_raw = df[feature_cols].astype(np.float32).values
y_raw = df["Class_encoded"].values

# Stratification key (DAS)
stratify_key = df["Subclass"].values
```

In [4]:
```
# ------------------------------
# 3) Train/Validation split (75/25)
# ------------------------------
X_tr_raw, X_val_raw, y_tr, y_val = train_test_split(
    X_raw, y_raw, test_size=0.25, stratify=stratify_key, random_state=42
)
```

In [5]:
```
# ------------------------------
# 4) KMeansSMOTE hyperparameter tuning with Optuna
# ------------------------------
# Objective: maximize silhouette score on resampled data (unsupervised structure)
def _minority_count(y):
    """
    Compute the minimum class count in a label vector.

    Parameters:
```

```python
        y (array-like of int): Class labels of shape (n_samples,).

    Returns:
        int: The size of the smallest class. Returns 0 if `y` is empty.
    """
    _, counts = np.unique(y, return_counts=True)
    return int(counts.min()) if counts.size else 0

def objective(trial):
    """
    Objective function for Optuna optimization of KMeansSMOTE hyperparameters.
    It resamples the training data using the candidate parameters and evaluates
    cluster structure by the silhouette score (higher is better).

    Parameters:
        trial (optuna.trial.Trial): Optuna trial providing parameter suggestions.

    Returns:
        float: Silhouette score on the resampled feature space. Returns -1.0
               when resampling fails or yields a degenerate label set.
    """
    if _minority_count(y_tr) < 2:
        return -1.0

    # Search space
    k_neighbors = trial.suggest_int("k_neighbors", 1, 15)
    n_clusters  = trial.suggest_int("n_clusters", 1, 15)
    cbt         = trial.suggest_float("cluster_balance_threshold", 0.01, 0.30)

    # Build KMeansSMOTE
    kmeans = KMeans(n_clusters=n_clusters, random_state=42)
    smote  = KMeansSMOTE(
        sampling_strategy="auto",
        k_neighbors=k_neighbors,
        kmeans_estimator=kmeans,
        cluster_balance_threshold=cbt,
        random_state=42,
    )

    # Attempt resampling; if it fails or results in a degenerate set, penalize
    try:
        X_res, y_res = smote.fit_resample(X_tr_raw, y_tr)
    except Exception:
        return -1.0
    if len(np.unique(y_res)) < 2:
        return -1.0

    # Silhouette on KMeans labels in the resampled space
    labels = KMeans(n_clusters=len(np.unique(y_res)), random_state=42).fit_predict(X_res)
    return silhouette_score(X_res, labels)

# Run Optuna (TPE; 300 trials)
study = optuna.create_study(direction="maximize", sampler=TPESampler(seed=42))
study.optimize(objective, n_trials=300, show_progress_bar=True)

best = study.best_params
print("Best SMOTE params:", best)

# Save Optuna trials and best params for provenance
trials_df = pd.DataFrame(
    [{"trial": t.number, "value": t.value, **t.params} for t in study.trials]
)
trials_df.to_csv(os.path.join(csv_dir, "optuna_kmeanssmote_trials.csv"), index=False)
with open(os.path.join(csv_dir, "optuna_kmeanssmote_best_params.json"), "w", encoding="utf-8") as f:
    json.dump(best, f, ensure_ascii=False, indent=2)
```

```
Best trial: 9. Best value: 0.724438: 100%|██████████████████████████████████████████| 300/300 [00:04<00:00, 70.72it/s]
Best SMOTE params: {'k_neighbors': 8, 'n_clusters': 9, 'cluster_balance_threshold': 0.023470619688799934}
```

In [6]:

```python
# -------------------------------
# 5) Robust resampling routine with safety fallbacks
# -------------------------------
# Gradually relax constraints if KMeansSMOTE errors occur
def _fit_resample_kmeanssmote_safe(X, y, params):
    """
    Apply KMeansSMOTE with safety bounds and staged fallbacks to avoid common
    failure modes on small/imbalanced datasets.

    Strategy:
      1) Bound k_neighbors and n_clusters by feasibility given the minority size.
      2) Try the bounded best params; if it fails, set cluster_balance_threshold=0.0.
      3) If still failing, decrement n_clusters stepwise down to 1.
      4) If still failing, decrement k_neighbors stepwise (recomputing the
         feasible n_clusters upper bound at each step).
      5) Raise ValueError if all attempts fail.

    Parameters:
        X (np.ndarray): Feature matrix of shape (n_samples, n_features).
        y (array-like of int): Class labels of shape (n_samples,).
        params (dict): Candidate parameters with keys
                       {"k_neighbors", "n_clusters", "cluster_balance_threshold"}.

    Returns:
        Tuple[np.ndarray, np.ndarray]: (X_resampled, y_resampled).

    Raises:
        ValueError: If the minority class has < 2 samples or all fallback
                    attempts fail to produce a valid resample.
    """
    n_minority = _minority_count(y)
    if n_minority < 2:
```

```python
        raise ValueError("KMeansSMOTE cannot run: minority class has < 2 samples.")

    # Safety bounds only (値は極力維持しつつ下限/上限の整合を取る)
    k   = max(1, min(int(params["k_neighbors"]), n_minority - 1))
    mxc = max(1, n_minority // (k + 1))
    ncl = max(1, min(int(params["n_clusters"]), min(10, mxc)))
    cbt = float(params["cluster_balance_threshold"])

    def _make(cbt_, ncl_, k_):
        km = KMeans(n_clusters=int(ncl_), random_state=42)
        return KMeansSMOTE(
            sampling_strategy="auto",
            k_neighbors=int(k_),
            kmeans_estimator=km,
            cluster_balance_threshold=float(cbt_),
            random_state=42,
        )

    # Try best (after safety bounding)
    try:
        return _make(cbt, ncl, k).fit_resample(X, y)
    except ValueError as e:
        # Known upstream bug pattern: retry with cbt=0.0
        if "cannot convert float NaN to integer" not in str(e):
            raise

    # Fallback 1: cbt -> 0.0
    try:
        return _make(0.0, ncl, k).fit_resample(X, y)
    except ValueError:
        pass

    # Fallback 2: gradually reduce n_clusters
    for nc in range(ncl - 1, 0, -1):
        try:
            return _make(0.0, nc, k).fit_resample(X, y)
        except ValueError:
            continue

    # Fallback 3: gradually reduce k_neighbors and re-cap n_clusters accordingly
    for kk in range(k - 1, 0, -1):
        mxc = max(1, n_minority // (kk + 1))
        nc  = min(ncl, mxc)
        try:
            return _make(0.0, nc, kk).fit_resample(X, y)
        except ValueError:
            continue

    raise ValueError("KMeansSMOTE failed after parameter relaxation steps.")

# --- Handle NaN/inf by column-mean imputation on the training block only ---
X_fit  = np.asarray(X_tr_raw, dtype=float)
finite = np.isfinite(X_fit)
if not finite.all():
    with np.errstate(all="ignore"):
        col_mean = np.nanmean(np.where(finite, X_fit, np.nan), axis=0)
    col_mean = np.where(np.isfinite(col_mean), col_mean, 0.0)
    rows, cols = np.where(~finite)
    X_fit[rows, cols] = col_mean[cols]

# --- Apply robust KMeansSMOTE ---
X_tr_sm, y_tr_sm = _fit_resample_kmeanssmote_safe(X_fit, y_tr, best)
```

In [7]:
```python
# -------------------------------
# 6) Data preprocessing
# -------------------------------
scaler = StandardScaler()
X_tr   = scaler.fit_transform(X_tr_sm)
X_val  = scaler.transform(X_val_raw)
X_full = scaler.transform(X_raw)

# --- Metrics helper ---
def metrics(y_true, y_pred, y_prob):
    """
    Compute core classification metrics with macro-averaging.

    Parameters:
        y_true (array-like of int): True labels of shape (n_samples,).
        y_pred (array-like of int): Predicted labels of shape (n_samples,).
        y_prob (np.ndarray): Predicted class probabilities of shape
                             (n_samples, n_classes).

    Returns:
        dict: {
            "acc": float,       # Accuracy
            "bal_acc": float,   # Balanced accuracy
            "prec": float,      # Macro precision
            "rec": float,       # Macro recall
            "f1": float,        # Macro F1-score
            "auc": float or np.nan,  # Macro ROC-AUC (OvR); np.nan if single-class
            "ll": float         # Log loss
        }
    """
    prec, rec, f1, _ = precision_recall_fscore_support(
        y_true, y_pred, average="macro", zero_division=0
    )
    if len(np.unique(y_true)) == 1:
        auc_macro = np.nan
    else:
        auc_macro = roc_auc_score(
            y_true, y_prob,
```

```python
            labels=np.arange(n_classes),
            multi_class="ovr",
            average="macro",
        )
    return dict(
        acc     = accuracy_score(y_true, y_pred),
        bal_acc = balanced_accuracy_score(y_true, y_pred),
        prec    = prec,
        rec     = rec,
        f1      = f1,
        auc     = auc_macro,
        ll      = log_loss(y_true, y_prob, labels=np.arange(n_classes)),
    )
```

In [8]:
```python
# ------------------------------
# 8) Train AutoTabPFN (post-hoc ensemble)
# ------------------------------
auto_clf = AutoTabPFNClassifier(
    balance_probabilities=True,
    max_time=3000,
    presets="best_quality",
    eval_metric="roc_auc_ovo_macro",
    device="cuda",
    phe_init_args={"verbosity": 0},
    phe_fit_args={"learning_curves": True},
    random_state=42,
)

auto_clf.fit(X_tr, y_tr_sm)
```

Out[8]:

```
AutoTabPFNClassifier

AutoTabPFNClassifier(balance_probabilities=True, device='cuda',
                     eval_metric='roc_auc_ovo_macro', max_time=3000,
                     phe_fit_args={'learning_curves': True},
                     phe_init_args={'verbosity': 0}, presets='best_quality',
                     random_state=42)
```

In [9]:
```python
# ------------------------------
# 9) Validation evaluation (25% hold-out)
# ------------------------------
vp_val = auto_clf.predict(X_val)
pb_val = auto_clf.predict_proba(X_val)
val_metrics = metrics(y_val, vp_val, pb_val)

# Classification Report
print("\n=== Classification Report ===")
print(classification_report(y_val, vp_val, target_names=class_names))

# Save textual report
with open(os.path.join(csv_dir, "validation_classification_report.txt"), "w", encoding="utf-8") as f:
    f.write(classification_report(y_val, vp_val, target_names=class_names))

# Confusion matrix (Blues)
cm_val = confusion_matrix(y_val, vp_val)
fig, ax = plt.subplots(figsize=(6, 5))
sns.heatmap(
    cm_val,
    annot=True, fmt="d",
    cmap="Blues",
    xticklabels=class_names, yticklabels=class_names,
    annot_kws={"size": 20},
    ax=ax
)
ax.set_xlabel("Predicted", fontsize=20)
ax.set_ylabel("True", fontsize=20)
ax.set_title("Validation Confusion Matrix", fontsize=20)
ax.set_xticklabels(ax.get_xticklabels(), fontsize=15)
ax.set_yticklabels(ax.get_yticklabels(), fontsize=15)
cbar = ax.collections[0].colorbar
cbar.ax.tick_params(labelsize=15)
plt.savefig("./results/plots/ConfusionMatrix_validation.png", dpi=300, bbox_inches="tight")
plt.show()

# Bar chart of metrics
metric_names = [
    "Accuracy", "Balanced Accuracy", "macro Precision",
    "macro Recall", "macro F1-Score", "macro ROC-AUC", "Log Loss"
]
metric_keys  = ["acc", "bal_acc", "prec", "rec", "f1", "auc", "ll"]
values = [val_metrics[k] for k in metric_keys]

fig, ax = plt.subplots(figsize=(4, 4))
bars = ax.barh(metric_names, values, color="C0")
ax.invert_yaxis()
for bar, v in zip(bars, values):
    ax.text(
        v - 0.01,
        bar.get_y() + bar.get_height() / 2,
        f"{v:.3f}",
        ha="right", va="center",
        fontsize=15
    )
ax.set_xlabel("Score", fontsize=20)
ax.set_title("Validation Metrics", fontsize=20)
ax.tick_params(axis="x", labelsize=15)
ax.tick_params(axis="y", labelsize=15)
```

```python
plt.savefig("./results/plots/EvaluationMetrics_validation.png", dpi=300, bbox_inches="tight")
plt.show()
```

```
=== Classification Report ===
              precision    recall  f1-score   support

           1       0.90      0.86      0.88        22
           2       0.67      0.80      0.73         5
           3       1.00      0.89      0.94        19
           4       0.75      0.86      0.80         7
           5       0.91      0.95      0.93        22

    accuracy                           0.89        75
   macro avg       0.85      0.87      0.86        75
weighted avg       0.90      0.89      0.90        75
```

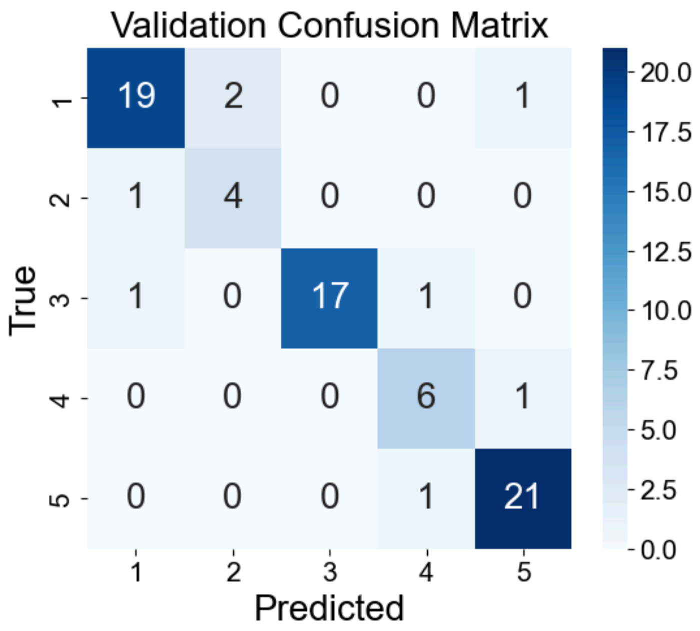


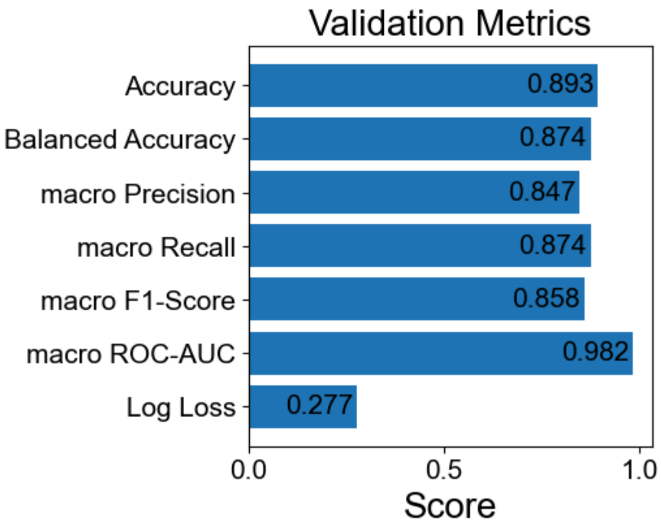


In [10]:

```python
# ------------------------------
# 10) External ("unseen") evaluation
# ------------------------------
df_unseen = pd.read_csv("Unseen_2_woBlank_wClass.csv").dropna(subset=["Class"])
df_unseen["Class_encoded"] = le.transform(df_unseen["Class"])

X_unseen = scaler.transform(df_unseen[feature_cols].astype(np.float32).values)
y_unseen = df_unseen["Class_encoded"].values

pred_u = auto_clf.predict(X_unseen)
prob_u = auto_clf.predict_proba(X_unseen)
unseen_metrics = metrics(y_unseen, pred_u, prob_u)

# Classification Report
print("\n=== Classification Report ===")
print(classification_report(y_unseen, pred_u, target_names=class_names))

# Save textual report
with open(os.path.join(csv_dir, "unseen_classification_report.txt"), "w", encoding="utf-8") as f:
    f.write(classification_report(y_unseen, pred_u, target_names=class_names))

# Confusion matrix (Oranges)
cm_u = confusion_matrix(y_unseen, pred_u)
fig, ax = plt.subplots(figsize=(6, 5))
sns.heatmap(
    cm_u,
    annot=True, fmt="d",
    cmap="Oranges",
    xticklabels=class_names, yticklabels=class_names,
    annot_kws={"size": 20},
    ax=ax
```

```python
)
ax.set_xlabel("Predicted", fontsize=20)
ax.set_ylabel("True", fontsize=20)
ax.set_title("Unseen Confusion Matrix", fontsize=20)
ax.set_xticklabels(ax.get_xticklabels(), fontsize=15)
ax.set_yticklabels(ax.get_yticklabels(), fontsize=15)
cbar = ax.collections[0].colorbar
cbar.ax.tick_params(labelsize=15)
plt.savefig("./results/plots/ConfusionMatrix_unseen.png", dpi=300, bbox_inches="tight")
plt.show()

# Bar chart of metrics
values = [unseen_metrics[k] for k in metric_keys]
fig, ax = plt.subplots(figsize=(4, 4))
bars = ax.barh(metric_names, values, color="C1")
ax.invert_yaxis()
for bar, v in zip(bars, values):
    ax.text(
        v - 0.01,
        bar.get_y() + bar.get_height() / 2,
        f"{v:.3f}",
        ha="right", va="center",
        fontsize=15
    )
ax.set_xlabel("Score", fontsize=20)
ax.set_title("Unseen Data Metrics", fontsize=20)
ax.tick_params(axis="x", labelsize=15)
ax.tick_params(axis="y", labelsize=15)
plt.savefig("./results/plots/EvaluationMetrics_unseen.png", dpi=300, bbox_inches="tight")
plt.show()
```

```
=== Classification Report ===
              precision    recall  f1-score   support

           1       0.89      0.89      0.89        18
           2       0.67      0.50      0.57         4
           3       0.94      1.00      0.97        16
           4       0.86      1.00      0.92         6
           5       1.00      0.94      0.97        16

    accuracy                           0.92        60
   macro avg       0.87      0.87      0.86        60
weighted avg       0.91      0.92      0.91        60
```

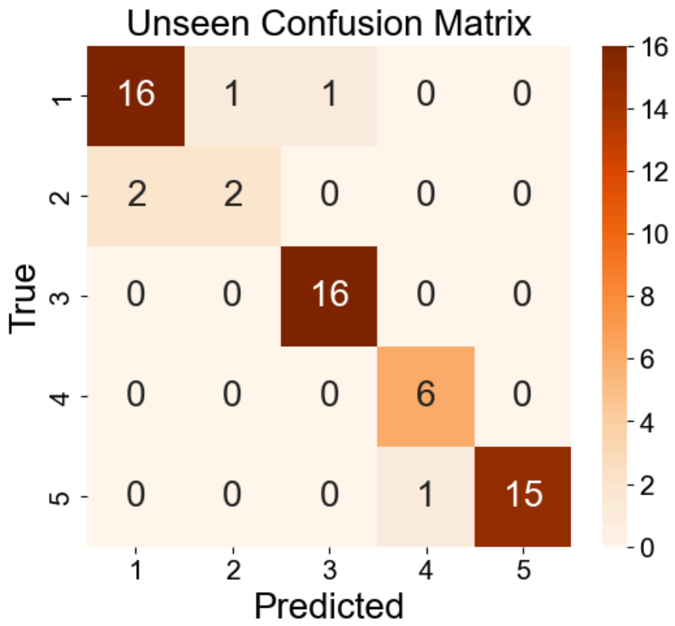


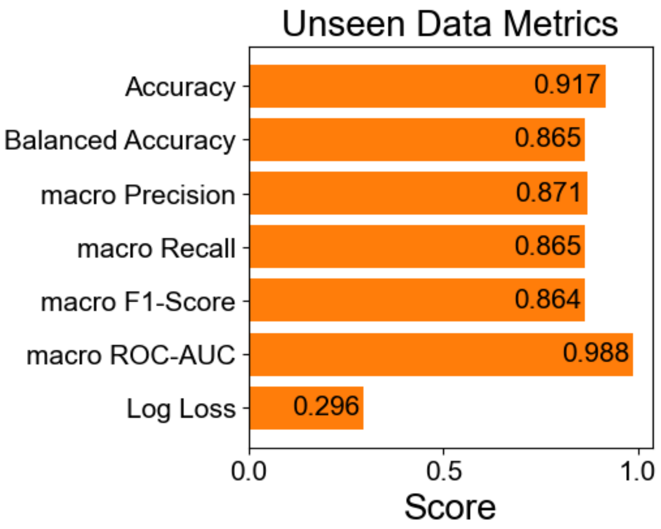


In [11]:
```python
# ------------------------------
# 11) ROC & PR curves on external set
# ------------------------------
```

```python
# ROC curve
y_bin = label_binarize(y_unseen, classes=np.arange(n_classes))
fpr, tpr, roc_auc = {}, {}, {}
for i in range(n_classes):
    fpr[i], tpr[i], _ = roc_curve(y_bin[:, i], prob_u[:, i])
    roc_auc[i] = auc(fpr[i], tpr[i])
fpr["macro"], tpr["macro"], _ = roc_curve(y_bin.ravel(), prob_u.ravel())
roc_auc["macro"] = auc(fpr["macro"], tpr["macro"])

fig, ax = plt.subplots(figsize=(6, 5))
colors = plt.cm.viridis(np.linspace(0, 1, n_classes))
for i, color in zip(range(n_classes), colors):
    ax.plot(
        fpr[i], tpr[i],
        label=f"{class_names[i]} (AUC={roc_auc[i]:.3f})",
        color=color, linewidth=2
    )
ax.plot([0, 1], [0, 1], linestyle="--", linewidth=2, color="grey")
ax.plot(
    fpr["macro"], tpr["macro"],
    label=f"Macro-average (AUC={roc_auc['macro']:.3f})",
    linestyle="--", linewidth=3, color="C1"
)
ax.set_xlabel("FPR", fontsize=20)
ax.set_ylabel("TPR", fontsize=20)
ax.set_title("ROC Curve - Unseen Data", fontsize=20)
ax.tick_params(axis="x", labelsize=15)
ax.tick_params(axis="y", labelsize=15)
ax.legend(loc="lower right", fontsize=10)
plt.savefig("./results/plots/ROCcurve_unseen.png", dpi=300, bbox_inches="tight")
plt.show()

# PR curve
precision, recall, pr_auc = {}, {}, {}
for i in range(n_classes):
    precision[i], recall[i], _ = precision_recall_curve(y_bin[:, i], prob_u[:, i])
    pr_auc[i] = auc(recall[i], precision[i])
precision["macro"], recall["macro"], _ = precision_recall_curve(y_bin.ravel(), prob_u.ravel())
pr_auc["macro"] = auc(recall["macro"], precision["macro"])

fig, ax = plt.subplots(figsize=(6, 5))
for i, color in zip(range(n_classes), colors):
    ax.plot(
        recall[i], precision[i],
        label=f"{class_names[i]} (AUC={pr_auc[i]:.3f})",
        color=color, linewidth=2
    )
ax.plot(
    recall["macro"], precision["macro"],
    label=f"Macro-average (AUC={pr_auc['macro']:.3f})",
    linestyle="--", linewidth=3, color="C1"
)
ax.set_xlabel("Recall", fontsize=20)
ax.set_ylabel("Precision", fontsize=20)
ax.set_title("Precision-Recall Curve - Unseen Data", fontsize=20)
ax.tick_params(axis="x", labelsize=15)
ax.tick_params(axis="y", labelsize=15)
ax.legend(loc="lower left", fontsize=10)
plt.savefig("./results/plots/PRcurve_unseen.png", dpi=300, bbox_inches="tight")
plt.show()
```

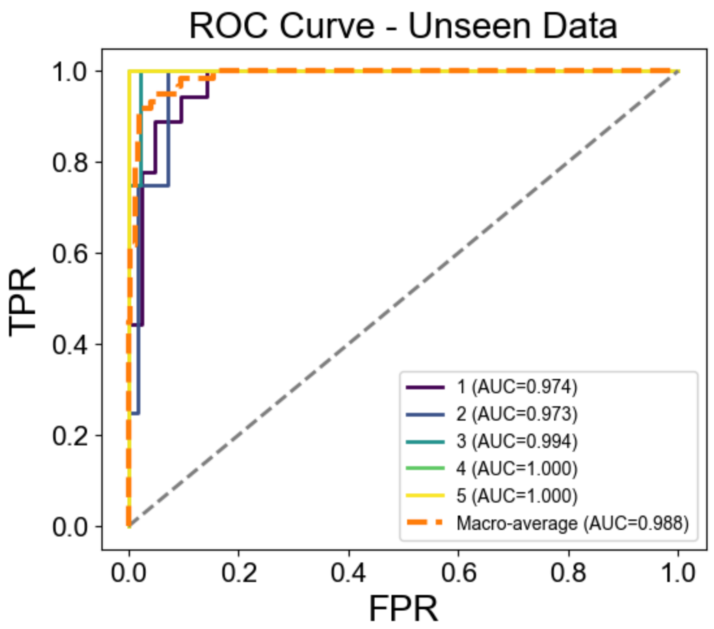

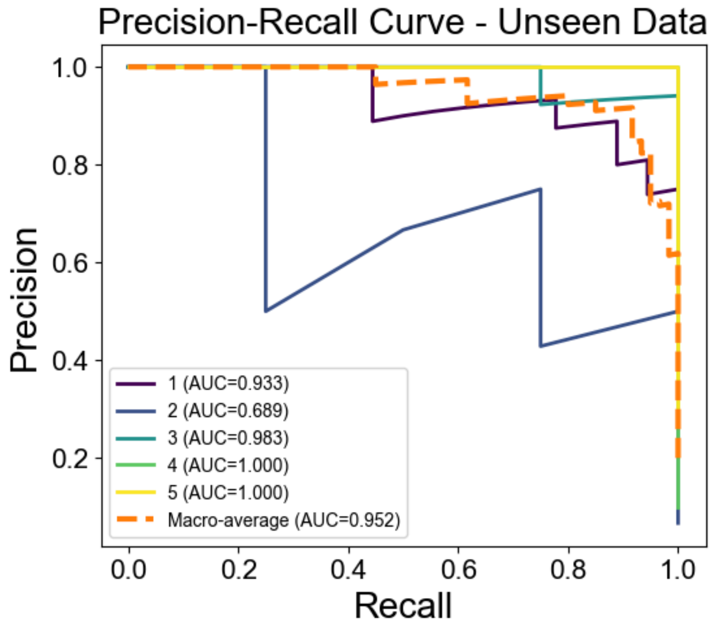


In [12]:

```
# ------------------------------
# 12) 95% stratified bootstrap CIs for external metrics
# ------------------------------
n_boot = 1000
rng    = np.random.RandomState(42)
cls_idx = {c: np.where(y_unseen == c)[0] for c in np.unique(y_unseen)}

boot_scores = defaultdict(list)
for _ in range(n_boot):
    # Stratified resampling per class
    sampled = []
    for c, idx_pool in cls_idx.items():
        sampled.extend(rng.choice(idx_pool, size=len(idx_pool), replace=True))
    sampled = np.asarray(sampled)

    m_bs = metrics(y_unseen[sampled], pred_u[sampled], prob_u[sampled])
    for k in metric_keys:
        boot_scores[k].append(m_bs[k])

# Compute and save CI table
ci_rows = []
print("\n=== Unseen Data Metrics 95% CI (stratified bootstrap) ===")
for k in metric_keys:
    vals = np.asarray(boot_scores[k])
    lower, upper = np.percentile(vals, [2.5, 97.5])
    ci_rows.append({"Metric": k, "Lower95": lower, "Upper95": upper})
    print(f"{k:<10s}: {lower:.3f} – {upper:.3f}")

pd.DataFrame(ci_rows).to_csv(os.path.join(csv_dir, "unseen_metrics_95CI.csv"), index=False)
```

```
=== Unseen Data Metrics 95% CI (stratified bootstrap) ===
acc       : 0.850 – 0.967
bal_acc   : 0.765 – 0.967
prec      : 0.722 – 0.980
rec       : 0.765 – 0.967
f1        : 0.738 – 0.966
auc       : 0.974 – 0.999
ll        : 0.142 – 0.473
```

In [ ]:

In [ ]:

In [ ]:
