## Supplementary material for "Time-resolved volatile organic compound profiling enables non-invasive detection of phenological progression in soybean": Supplemetal Materials: VOC_Supplementary Figs and Tables_ver11.pptx

### Slide 1
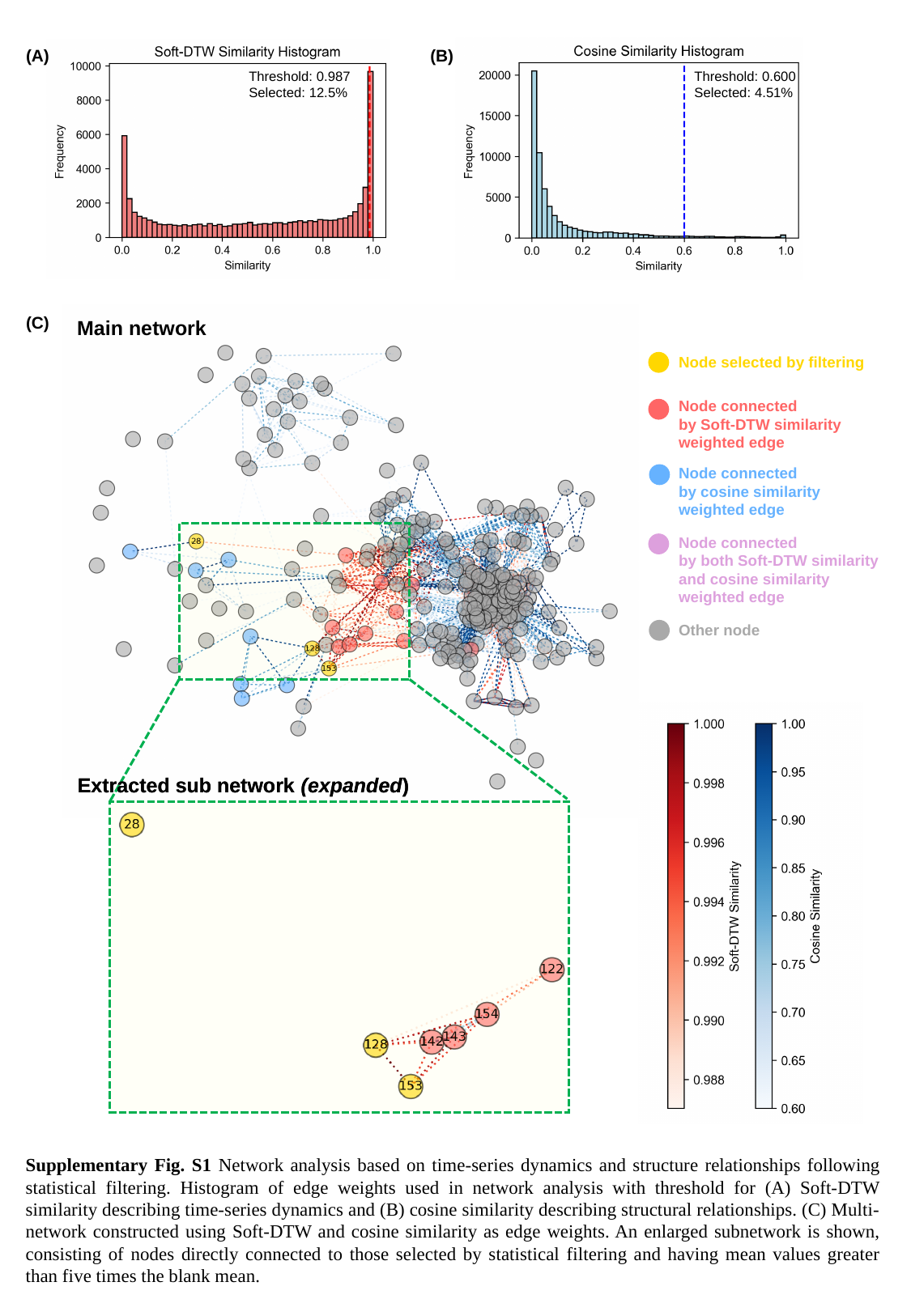

(B)
(A)
Threshold: 0.987
Selected: 12.5%
Threshold: 0.600
Selected: 4.51%
(C)
Main network
Node selected by filtering
Node connected
by Soft-DTW similarity
weighted edge
Node connected
by cosine similarity
weighted edge
Node connected
by both Soft-DTW similarity and cosine similarity
weighted edge
Other node
Extracted sub network (expanded)
Extracted sub network (expanded)
Supplementary Fig. S1 Network analysis based on time-series dynamics and structure relationships following statistical filtering. Histogram of edge weights used in network analysis with threshold for (A) Soft-DTW similarity describing time-series dynamics and (B) cosine similarity describing structural relationships. (C) Multi-network constructed using Soft-DTW and cosine similarity as edge weights. An enlarged subnetwork is shown, consisting of nodes directly connected to those selected by statistical filtering and having mean values greater than five times the blank mean.

### Slide 2
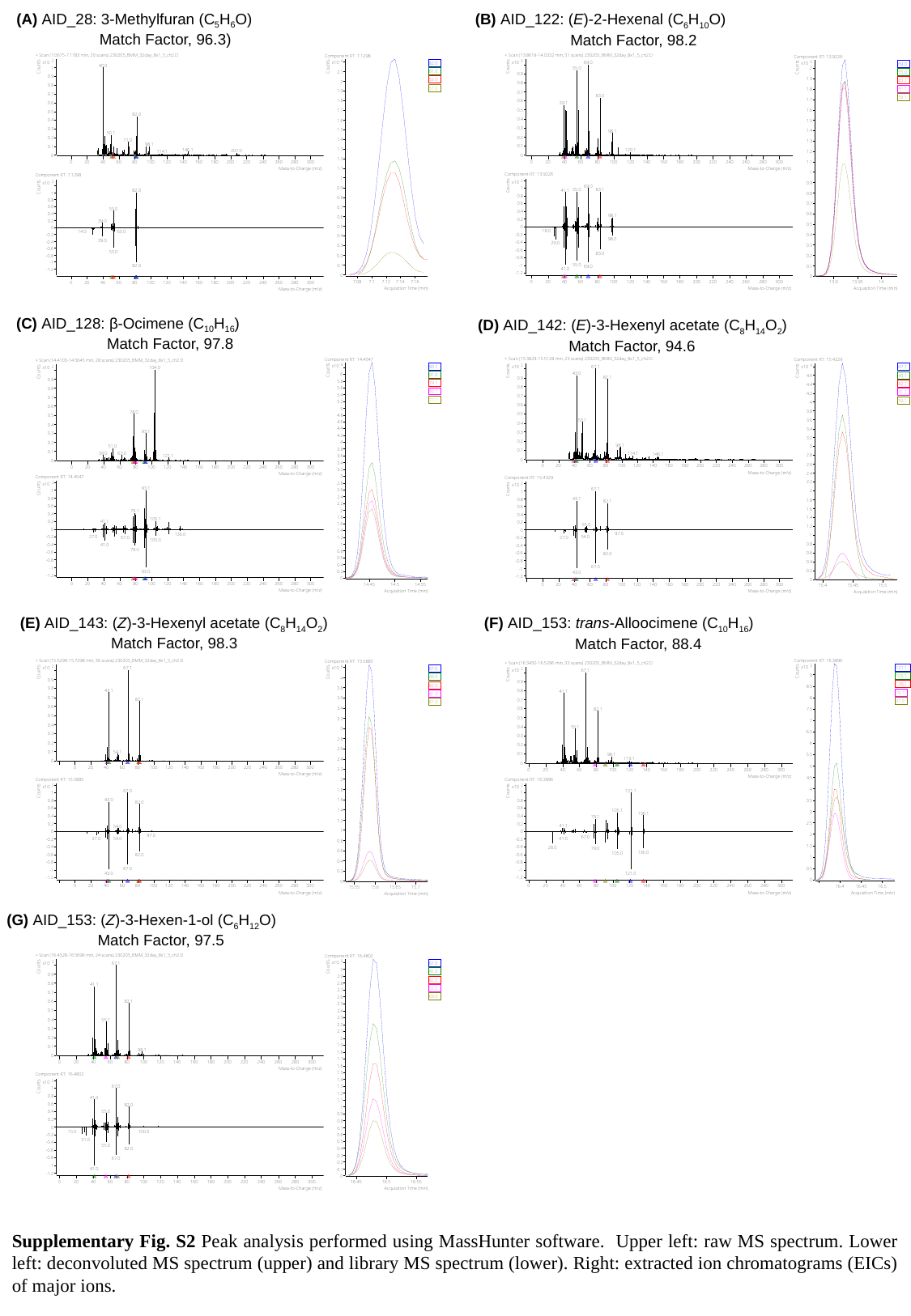

(A) AID_28: 3-Methylfuran (C_5_H_6_O)
Match Factor, 96.3)
(B) AID_122: (E)-2-Hexenal (C_6_H_10_O)
Match Factor, 98.2
(C) AID_128: β-Ocimene (C_10_H_16_)
Match Factor, 97.8
(D) AID_142: (E)-3-Hexenyl acetate (C_8_H_14_O_2_)
Match Factor, 94.6
(E) AID_143: (Z)-3-Hexenyl acetate (C_8_H_14_O_2_)
Match Factor, 98.3
(F) AID_153: trans-Alloocimene (C_10_H_16_)
Match Factor, 88.4
(G) AID_153: (Z)-3-Hexen-1-ol (C_6_H_12_O)
Match Factor, 97.5
Supplementary Fig. S2 Peak analysis performed using MassHunter software. Upper left: raw MS spectrum. Lower left: deconvoluted MS spectrum (upper) and library MS spectrum (lower). Right: extracted ion chromatograms (EICs) of major ions.

### Slide 3
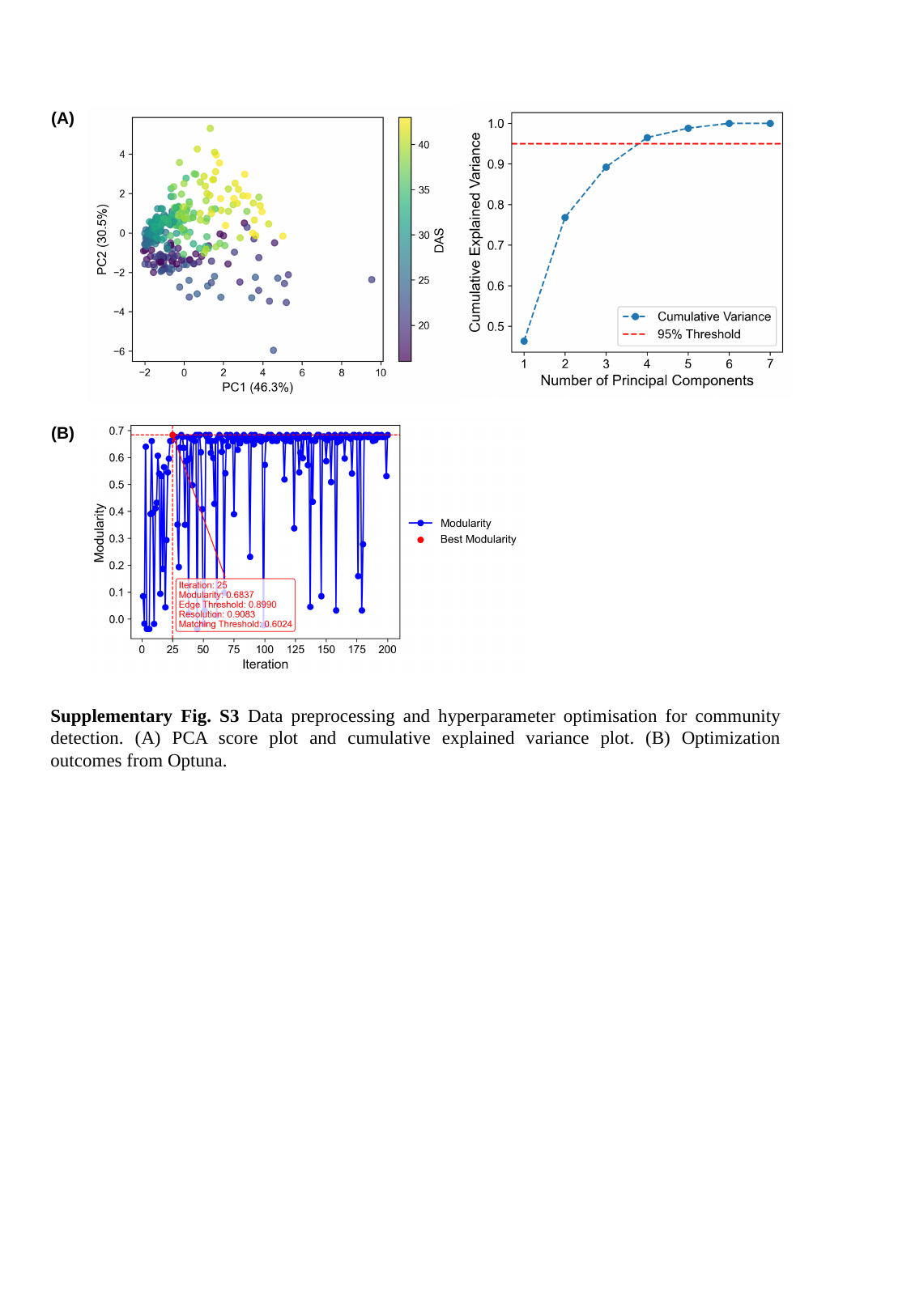

(A)
(B)
Supplementary Fig. S3 Data preprocessing and hyperparameter optimisation for community detection. (A) PCA score plot and cumulative explained variance plot. (B) Optimization outcomes from Optuna.

### Slide 4
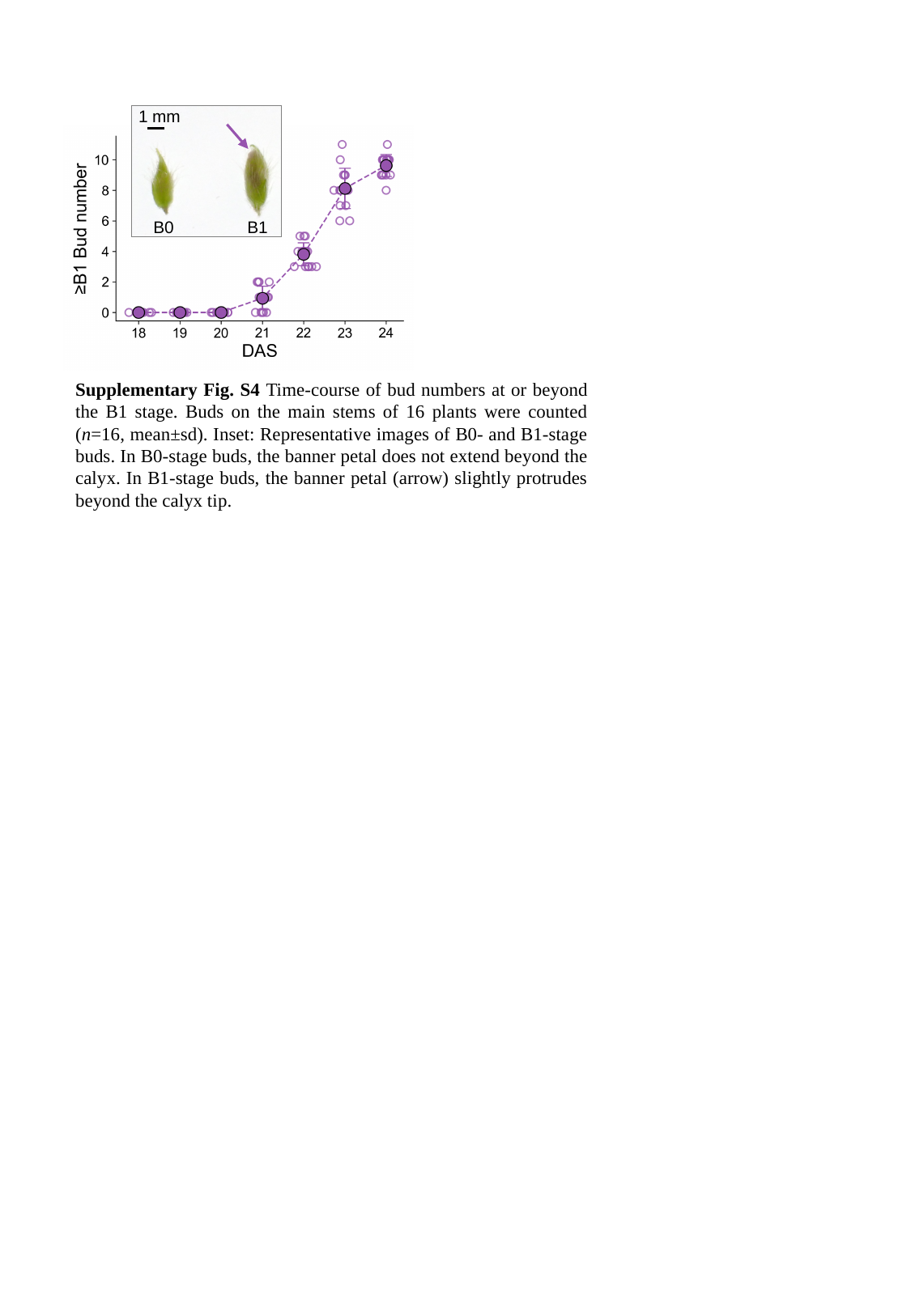

1 mm
B0
B1
Supplementary Fig. S4 Time-course of bud numbers at or beyond the B1 stage. Buds on the main stems of 16 plants were counted (n=16, mean±sd). Inset: Representative images of B0- and B1-stage buds. In B0-stage buds, the banner petal does not extend beyond the calyx. In B1-stage buds, the banner petal (arrow) slightly protrudes beyond the calyx tip.

### Slide 5
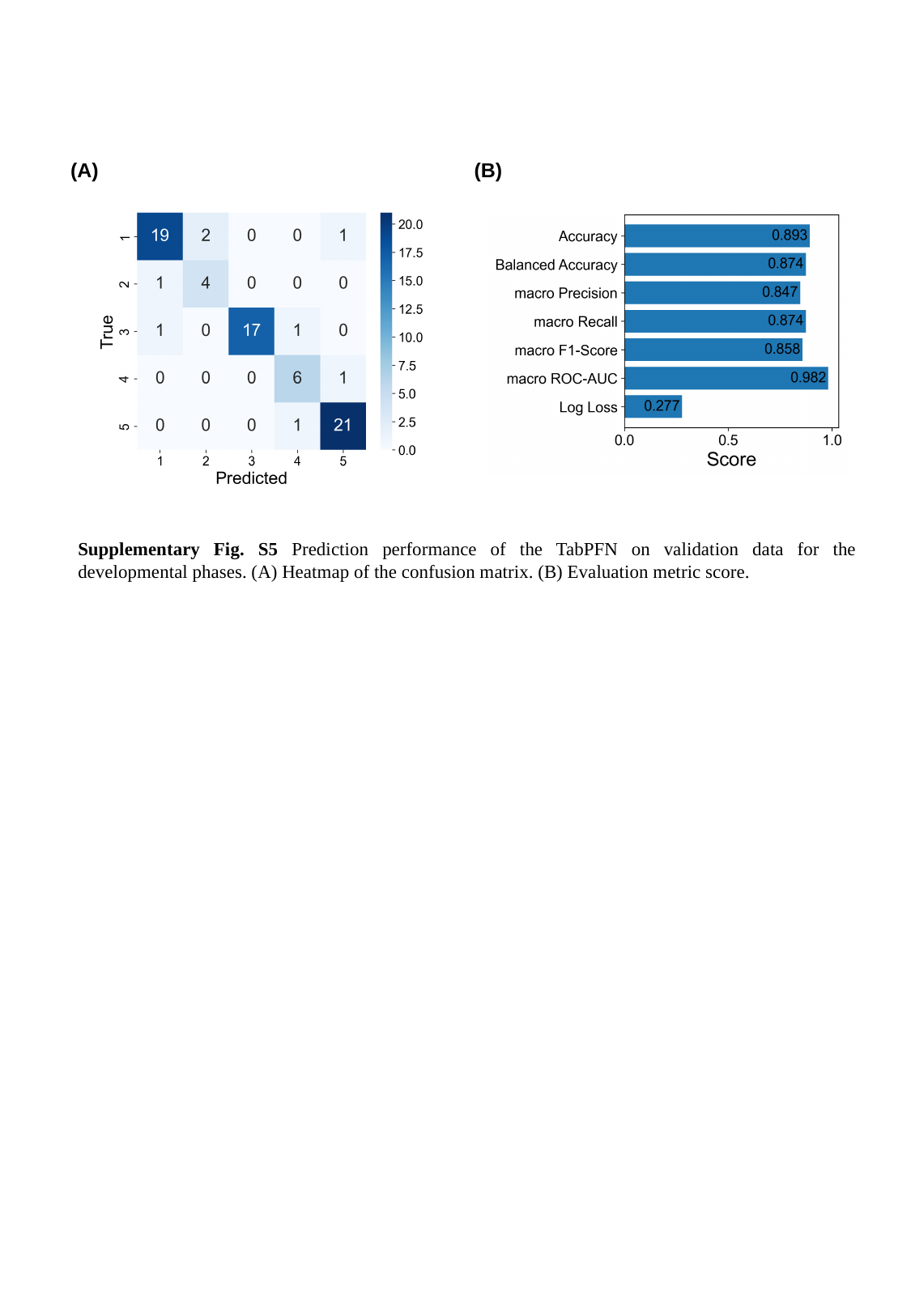

(A)
(B)
Supplementary Fig. S5 Prediction performance of the TabPFN on validation data for the developmental phases. (A) Heatmap of the confusion matrix. (B) Evaluation metric score.

### Slide 6
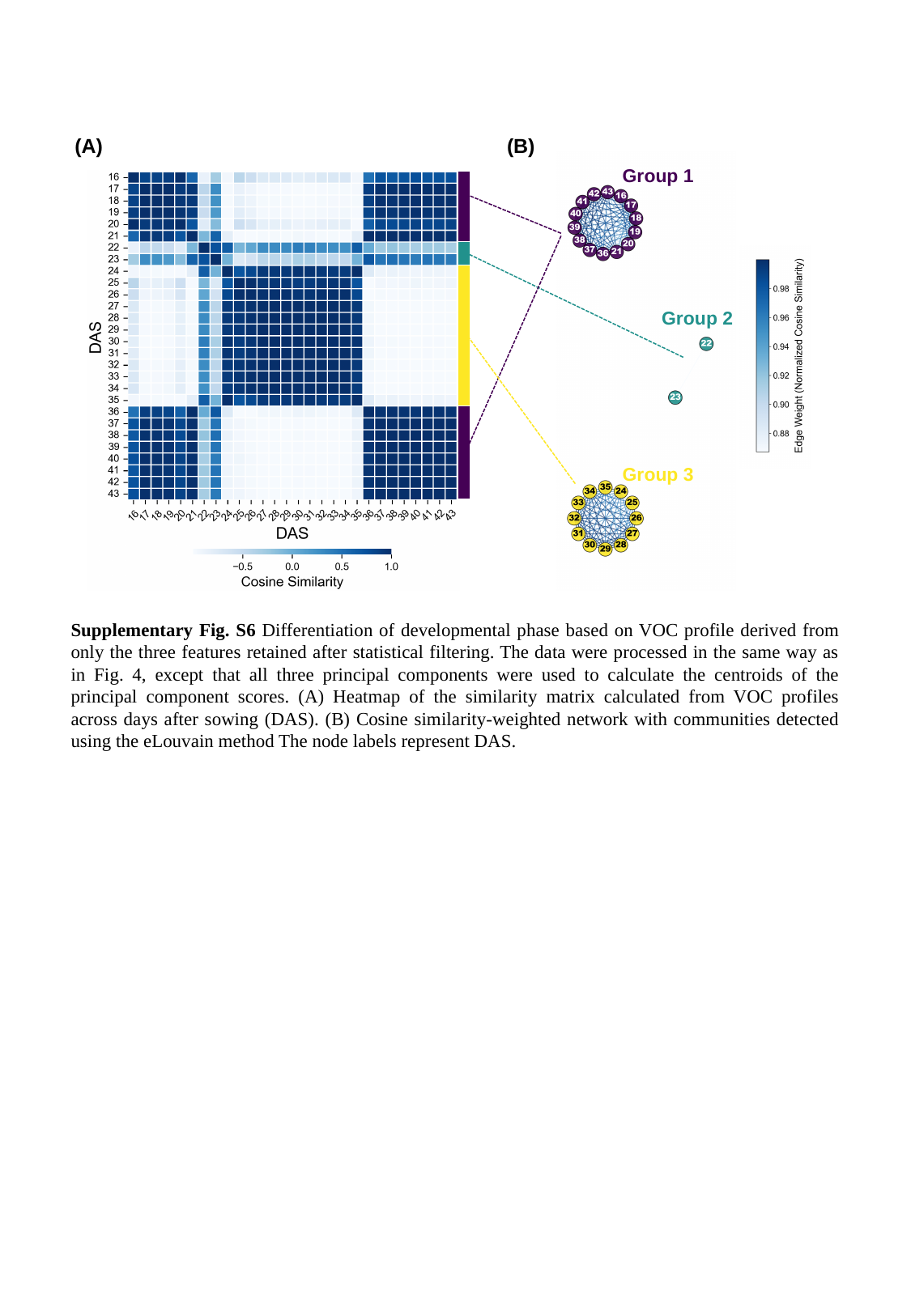

(A)
(B)
Group 1
Group 2
Group 3
Supplementary Fig. S6 Differentiation of developmental phase based on VOC profile derived from only the three features retained after statistical filtering. The data were processed in the same way as in Fig. 4, except that all three principal components were used to calculate the centroids of the principal component scores. (A) Heatmap of the similarity matrix calculated from VOC profiles across days after sowing (DAS). (B) Cosine similarity-weighted network with communities detected using the eLouvain method The node labels represent DAS.

### Slide 7
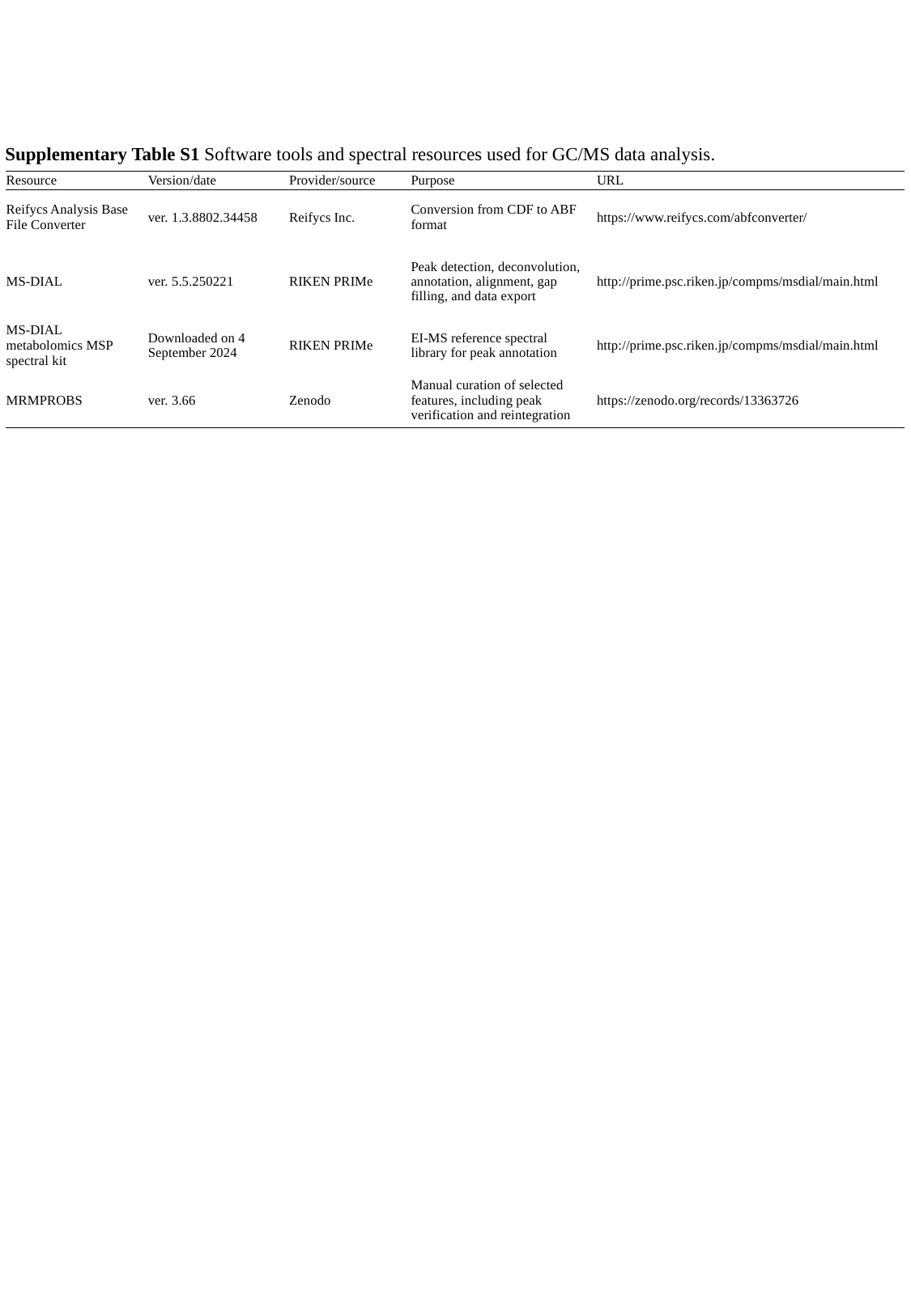

Supplementary Table S1 Software tools and spectral resources used for GC/MS data analysis.
| Resource | | Version/date | | Provider/source | | Purpose | | URL |
| --- | --- | --- | --- | --- | --- | --- | --- | --- |
| Reifycs Analysis Base File Converter | | ver. 1.3.8802.34458 | | Reifycs Inc. | | Conversion from CDF to ABF format | | https://www.reifycs.com/abfconverter/ |
| MS-DIAL | | ver. 5.5.250221 | | RIKEN PRIMe | | Peak detection, deconvolution, annotation, alignment, gap filling, and data export | | http://prime.psc.riken.jp/compms/msdial/main.html |
| MS-DIAL metabolomics MSP spectral kit | | Downloaded on 4 September 2024 | | RIKEN PRIMe | | EI-MS reference spectral library for peak annotation | | http://prime.psc.riken.jp/compms/msdial/main.html |
| MRMPROBS | | ver. 3.66 | | Zenodo | | Manual curation of selected features, including peak verification and reintegration | | https://zenodo.org/records/13363726 |

### Slide 8
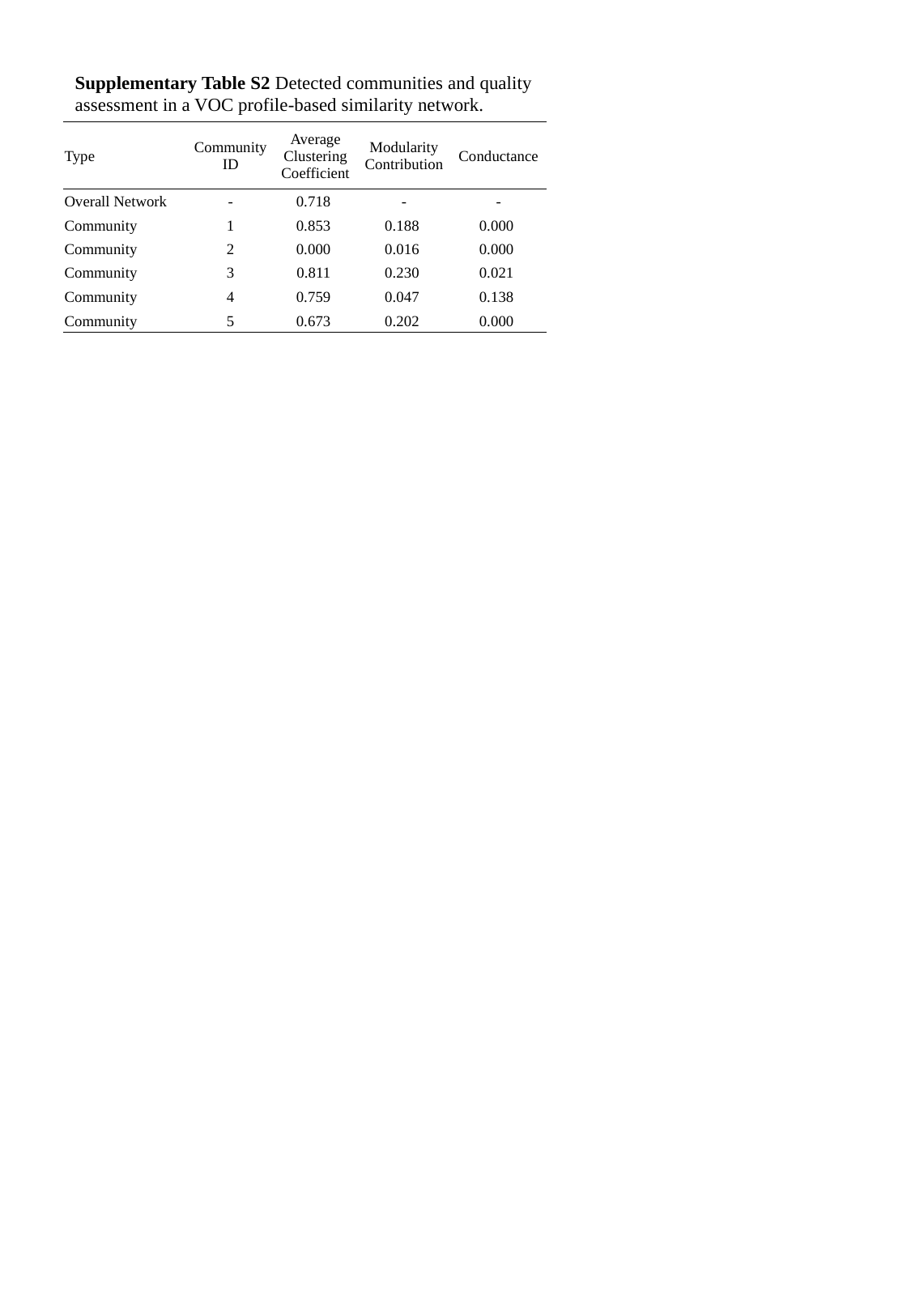

Supplementary Table S2 Detected communities and quality assessment in a VOC profile-based similarity network.
| Type | Community ID | Average Clustering Coefficient | Modularity Contribution | Conductance |
| --- | --- | --- | --- | --- |
| Overall Network | - | 0.718 | - | - |
| Community | 1 | 0.853 | 0.188 | 0.000 |
| Community | 2 | 0.000 | 0.016 | 0.000 |
| Community | 3 | 0.811 | 0.230 | 0.021 |
| Community | 4 | 0.759 | 0.047 | 0.138 |
| Community | 5 | 0.673 | 0.202 | 0.000 |

### Slide 9
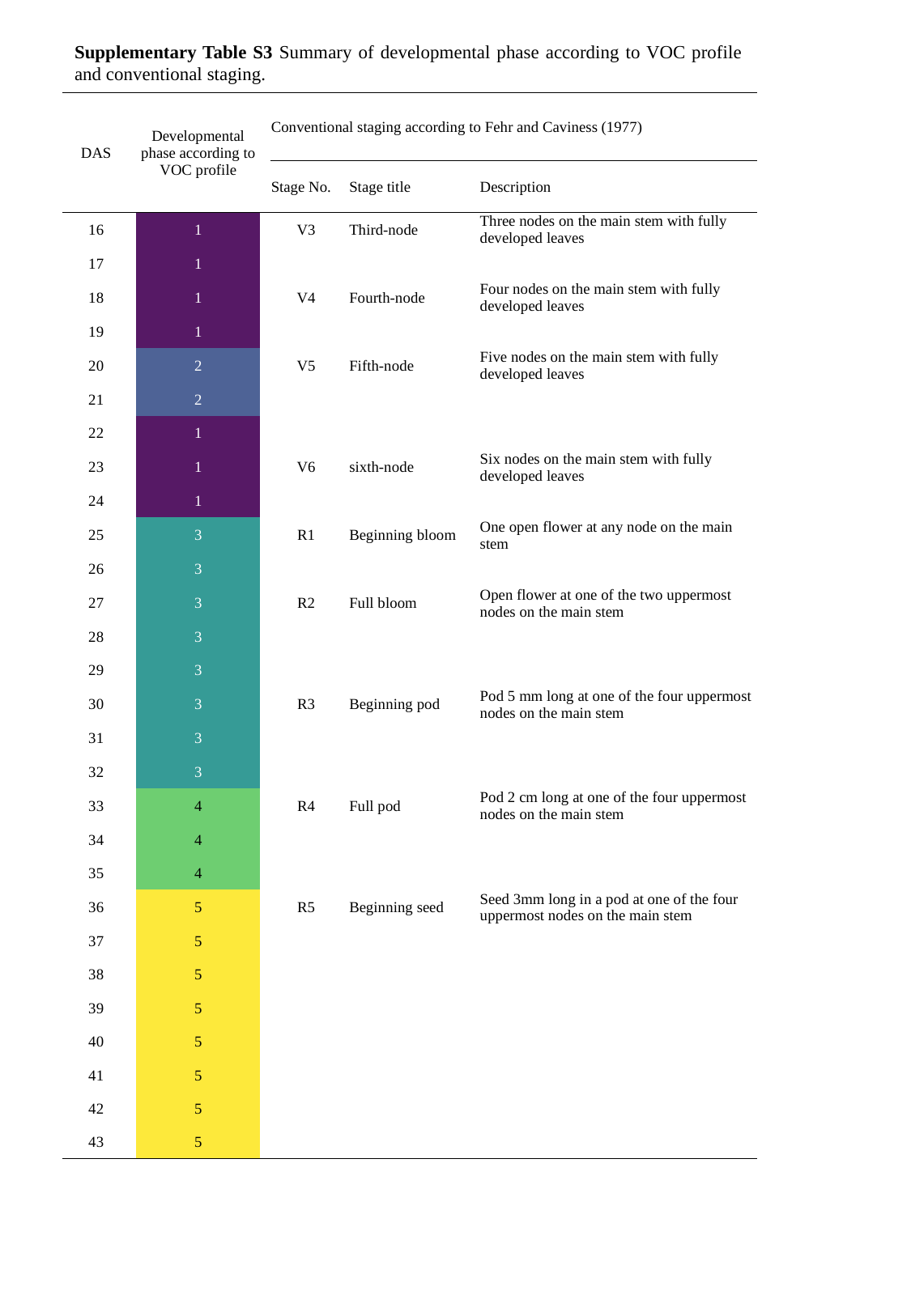

Supplementary Table S3 Summary of developmental phase according to VOC profile and conventional staging.
| DAS | | Developmental phase according to VOC profile | | Conventional staging according to Fehr and Caviness (1977) | | | | |
| --- | --- | --- | --- | --- | --- | --- | --- | --- |
| | | | | Stage No. | | Stage title | | Description |
| | | | | Stage No. | | Stage title | | Description |
| 16 | | 1 | | V3 | | Third-node | | Three nodes on the main stem with fully developed leaves |
| 17 | | 1 | | | | | | |
| 18 | | 1 | | V4 | | Fourth-node | | Four nodes on the main stem with fully developed leaves |
| 19 | | 1 | | | | | | |
| 20 | | 2 | | V5 | | Fifth-node | | Five nodes on the main stem with fully developed leaves |
| 21 | | 2 | | | | | | |
| 22 | | 1 | | | | | | |
| 23 | | 1 | | V6 | | sixth-node | | Six nodes on the main stem with fully developed leaves |
| 24 | | 1 | | | | | | |
| 25 | | 3 | | R1 | | Beginning bloom | | One open flower at any node on the main stem |
| 26 | | 3 | | | | | | |
| 27 | | 3 | | R2 | | Full bloom | | Open flower at one of the two uppermost nodes on the main stem |
| 28 | | 3 | | | | | | |
| 29 | | 3 | | | | | | |
| 30 | | 3 | | R3 | | Beginning pod | | Pod 5 mm long at one of the four uppermost nodes on the main stem |
| 31 | | 3 | | | | | | |
| 32 | | 3 | | | | | | |
| 33 | | 4 | | R4 | | Full pod | | Pod 2 cm long at one of the four uppermost nodes on the main stem |
| 34 | | 4 | | | | | | |
| 35 | | 4 | | | | | | |
| 36 | | 5 | | R5 | | Beginning seed | | Seed 3mm long in a pod at one of the four uppermost nodes on the main stem |
| 37 | | 5 | | | | | | |
| 38 | | 5 | | | | | | |
| 39 | | 5 | | | | | | |
| 40 | | 5 | | | | | | |
| 41 | | 5 | | | | | | |
| 42 | | 5 | | | | | | |
| 43 | | 5 | | | | | | |
